# A time resolved interaction analysis of Bem1 reconstructs the flow of Cdc42 during polar growth

**DOI:** 10.1101/723171

**Authors:** Sören Grinhagens, Alexander Dünkler, Yehui Wu, Lucia Rieger, Philipp Brenner, Thomas Gronemeyer, Nils Johnsson

## Abstract

Cdc42 organizes cellular polarity and directs the formation of cellular structures in many organisms. By locating Cdc24, the source of active Cdc42, to the growing edge of the yeast cell, the scaffold protein Bem1 is instrumental in shaping the cellular gradient of Cdc42. This gradient instructs bud formation, bud growth, or cytokinesis through the actions of a diverse set of effector proteins. To address how Bem1 participates in this transformation we systematically mapped its protein interactions in time and space. SPLIFF analysis defined a unique ensemble of Bem1 interaction-states for each cell cycle stage. The characterization of mutants of Bem1 that interact with a discrete subset of the interaction partners allowed to assign specific functions to different interaction states and identified the determinants for their cellular distributions. The analysis characterizes Bem1 as a cell cycle specific shuttle that distributes active Cdc42 from its source to its effectors and helps to convert the PAKs Cla4 and Ste20 into their active conformation.

## Introduction

Yeast cells sculpt their shape by controlling site, direction and rate of cell expansion. Bud formation, growth and cell separation are the visible consequences of polar cell growth in the budding yeast (Howell and Lew, 2012; Bi and Park, 2012). The many interactions that occur between the involved cell polarity proteins have the potential to serve as inbuilt switches that drive these morphological alterations. Accordingly, changes in the composition and structure of their interaction network should correlate with the different phases of cell growth. This assumption was not yet tested.

Yeast cells initiate bud formation at a predetermined site, expand the bud preferentially at its tip, switch in large buds to an isotropic growth, and finally reorient the growth axis during mitosis and cell separation (Howell and Lew, 2012). The Rho-like GTPase Cdc42 influences local cell expansion in all cell cycle phases by binding in its active, GTP-bound state to different effector proteins (Chiou et al., 2017). Among the many processes that are controlled by Cdc42_GTP_ are the organization of the septin- and actin cytoskeleton, the spatial organization of exocytosis, mating, osmolarity sensing, the mitotic exit, and the regulation of cell separation during cytokinesis (Pruyne et al., 2004; Bi and Park, 2012). The GEF Cdc24 and a variety of GAPs adjust the concentration of Cdc42_GTP_ at the cortex (Smith et al., 2002). This concentration changes over the cell cycle and peaks at G1/S, and at anaphase (Atkins et al., 2013).

Bem1 is the central scaffold for proteins that organize polarized growth in yeast (Matsui et al., 1996; Peterson et al., 1994; Bender et al., 1996; Chenevert et al., 1992). Bem1 binds Cdc24, Cdc42_GTP_, and several Cdc42_GTP_ effector proteins (Bose et al., 2001; Irazoqui et al., 2003). The specific functions of Bem1 are controversially discussed.

Bem1 was shown to determine the localization of Cdc24 at sites of polar growth and to modestly stimulate its activity (Woods et al., 2015; Smith et al., 2013; Rapali et al., 2017). Bem1 might also provide a positive feedback for polarity establishment and might serve as a platform for a negative feedback during later stages of the cell cycle (Kozubowski et al., 2008; Gulli et al., 2000; Kuo et al., 2014; Rapali et al., 2017). The protein is also part of the polarity cap during cell separation, cell mating and fusion. In addition Cdc42 in conjunction with Bem1 operate as upstream regulators of the pheromone response-, the filamentous growth- and the high osmolarity MAPK pathways (Leberer et al., 1997; Winters and Pryciak, 2005; Tanaka et al., 2014; Lyons et al., 1996).

Bem1 consists of two N-terminally located SH3 domains (SH3_a_, SH3_b_), a lipid- and membrane-binding PX domain, and a C-terminal PB domain (PB_Bem1_) (Bender et al., 1996; Matsui et al., 1996). SH3_a_ binds to the exocyst component Sec15. However, the removal of SH3_a_ hardly affects secretory vesicle fusion or other aspects of polarized growth (France et al., 2006). SH3_b_ interacts with well-characterized PxxP motives in the PAKs Cla4 and Ste20, and the polarity proteins Boi1 and Boi2 (Winters and Pryciak, 2005; Bender et al., 1996; Bose et al., 2001; Gorelik and Davidson, 2012). SH3_b_ harbors a C-terminal extension (CI) that binds Cdc42_GTP_ without impairing the interaction with its PxxP ligands (Takaku et al., 2010; Yamaguchi et al., 2007). The significance of this activity for polarity establishment or polar growth is not clear. PB_Bem1_ binds the C-terminal PB domain of Cdc24 (PB_Cdc24_). The interaction is highly affine and can be reconstituted with bacterially expressed proteins *in vitro* (Ogura et al., 2009; Ito et al., 2001). The PB_Bem1_-PB_Cdc24_ interaction localizes Cdc24 to sites of polar growth during all cell cycle stages (Woods et al., 2015; Witte et al., 2017; Butty et al., 2002). Although being intensely studied the molecular mechanisms behind Bem1’s effect on polarization as well its precisely regulated cellular distributions are not fully understood (Atkins et al., 2008; Kozubowski et al., 2008; Li and Wedlich-Soldner, 2009).

Here we probe the interaction network of Bem1 during different stages of polar growth to correlate changes in composition and architecture of the network with changes in cellular morphology and the activities of its binding partners. Our work describes Bem1 as a selective co-activator of the PAKs and other Cdc42-dependent processes.

## Results

### A Split-Ub based protein interaction map of Bem1

We searched for binding partners of Bem1 by performing a systematic Split-Ubiquitin (Split-Ub) screen of Bem1CRU against 548 N_ub_ fusion proteins known or suspected to be involved in different aspects of polarized growth in yeast (Hruby et al., 2011; Johnsson and Varshavsky, 1994). The screen confirmed known binding partners, and additionally identified Bud6, Msb1, Ras1, Ras2, Rga2, Nba1, Spa2, Cdc11, Fks1, and Bem1 as novel interaction partners of Bem1 (Fig. 1A).

**Figure 1.**
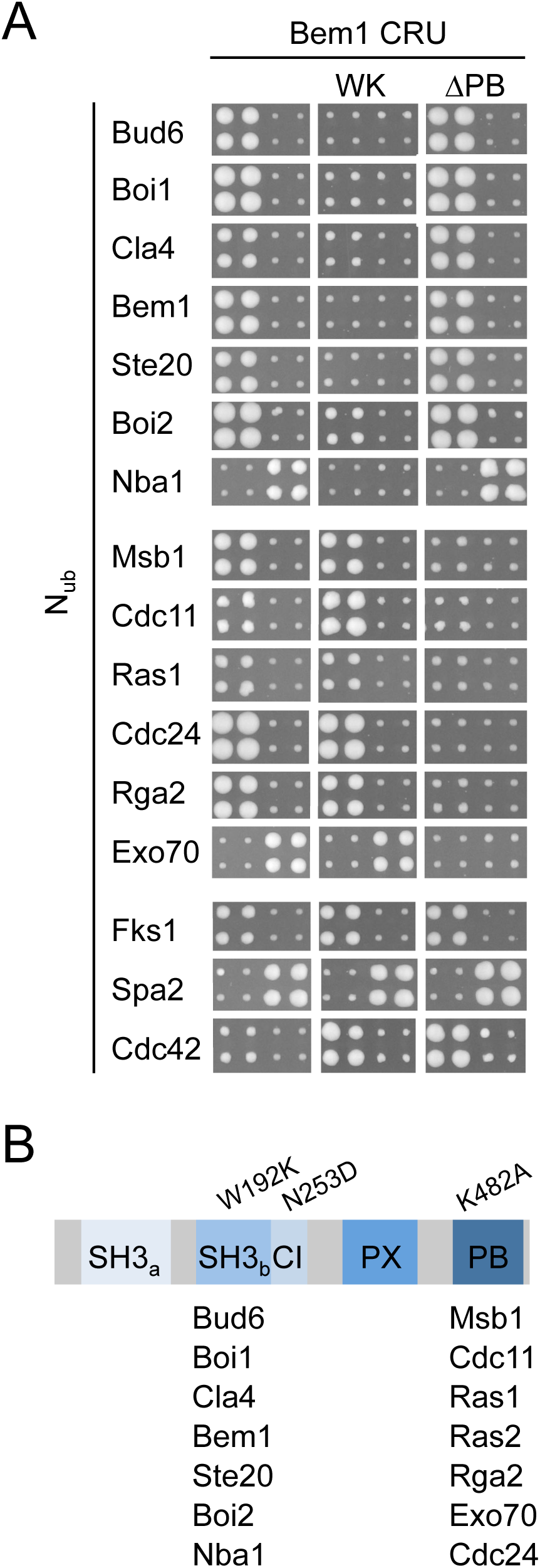
Interaction partners of Bem1. A. Yeast cells carrying Bem1CRU or either of its two mutants were each independently mated four times with the N_ub_-array and tested for interactions by growth on SD medium containing 5-FOA. Interaction is indicated by growth of the four matings. Shown are the cut outs of the quadruplets expressing the N_ub_ fusion of the interacting protein on the left, next to a fusion that does not interact, except for the cells expressing N_ub_-Cdc11, -Spa2, and -Exo70, that are shown on the right of cells expressing non-interacting N_ub_ fusions. B) Schematic presentation of Bem1 showing its different domains and the positions of the residue exchanges of the *bem1*-alleles used in this work. The domain-specific interaction partners of A) are listed below their respective domains.

To map the contact sites of the binding partners on the structure of Bem1, we repeated the screen with mutants of Bem1 that either carried the well-characterized W192K exchange in SH3_b_ (Bem1_WK_CRU) or lacked the C-terminal PB domain and thus the binding site to Cdc24 (Bem1_ΔPB_CRU) (Fig. 1B). A comparison with the interaction profile of the wild type CRU fusion revealed that Bem1_WK_ lost its interactions to Boi1, Ste20, Cla4, Bud6, Nba1 and Bem1, showed a strongly reduced binding to Boi2 but retained its interactions with Exo70, Cdc24, Cdc42, Cdc11, Rga2, Msb1, Ras2, and Ras1 (Fig. 1A, B). Deleting the PB domain in Bem1_ΔPB_CRU removed or strongly reduced the interactions of Bem1 to N_ub_-Cdc24, -Cdc11, -Rga2, -Msb1, -Ras1, -Ras2, and -Exo70 (Fig. 1A, B). As the interactions of Bem1 with Fks1, Cdc42, and Spa2 were not visibly affected by mutations in SH3_b_ or Bem1_PB_ we were unable to further resolve the Split-Ub measured interactions of Bem1 with those proteins (Fig. 1A, B).

### Physical dissection of the Bem1 interaction network

The Split-Ub assay detects direct as well as indirect protein-protein interactions(Hruby et al., 2011; Johnsson, 2014). The Split-Ub measured interactions between Bem1 and Boi1/2, Nba1 or Bud6 depend on SH3_b_ of Bem1 (Fig. 1A, B). Boi1/2 bind directly to SH3_b_ of Bem1 and were shown by Split-Ub analysis to also bind to Bud6 and Nba1(Kustermann et al., 2017; Bender et al., 1996). To test whether Boi1/2 might link Bem1 with Nba1 or Bud6 we introduced Bem1CRU together with N_ub_-Bud6, or N_ub_-Nba1 in a strain that carried either a deletion of *BOI1* or *BOI2*, or a deletion of *BOI2* on top of a mutation in the binding site of Boi1 for Bem1 (*Δboi2 boi1_ΔPxxP_*). Split-Ub assays in these strains confirmed, that the interactions between Bem1 and Bud6, or Bem1 and Nba1 clearly depend on Boi1/2 (Fig. 2A). Furthermore, the ability of Boi1/2 to homo- and to hetero-dimerize could explain the SH3_b_-dependency of the Split-Ub measured Bem1 dimerization (Fig. 1A, B) (Kustermann et al., 2017). The loss of interaction signal between Bem1CRU and N_ub_-Bem1 in a strain lacking *BOI2* and the binding site of Boi1 for Bem1 (*boi1_ΔPxxP_*) strongly suggests that the Split-Ub measured Bem1-Bem1 interaction is mediated by the multimerization of the Boi-proteins (Fig. 2A). In contrast to the Nba1-Boi1/2-Bem1 or the Bem1-Boi1/2-Bem1 complex, the Split-Ub detected interaction between Bud6 and Bem1 is already lost upon deletion of either *BOI1* or *BOI2* (Fig. 2A).

**Figure 2.**
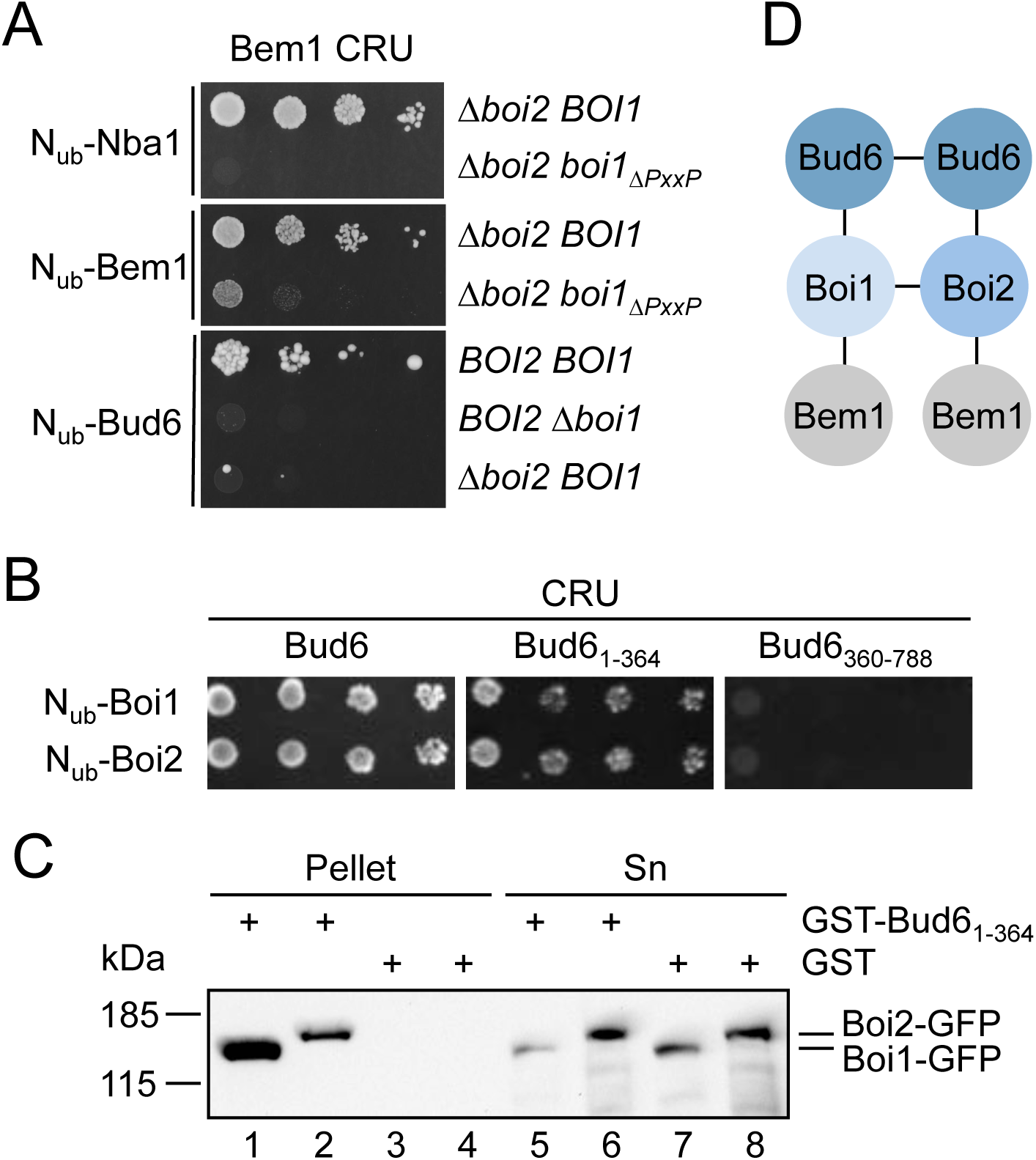
Characterization of the Bem1-Bud6 interaction state. A) Yeast cells carrying deletions of *BOI1,* or *BOI2,* or carrying a *BOI1* deletion together with the *boi1_ΔPxxP_* allele, and co-expressing CRU fusions to Bem1 together with the indicated N_ub_ fusions were grown to an OD_600_ of 1 and spotted in 10-fold serial dilutions onto medium containing 5-FOA. Interactions are indicated by the growth of the yeast cells. B) As in A) but with yeast cells co-expressing CRU fusions to Bud6_1-364_ or Bud6_360-788_, together with N_ub_ fusions to Boi1 and Boi2. C) Extracts of yeast cells expressing either Boi1-GFP (lanes 1, 3, 5, or Boi2-GFP (lanes 2, 4, 6, 8) were incubated with GST-(lanes 3, 4, 7, 8) or GST-Bud6_1-364_ (lanes 1, 2, 5, 6) -immobilized sepharose beads. Bound (lanes 1-4) and unbound (lanes 5-8) fractions were separated by SDS-PAGE and analyzed by anti-GFP antibodies after transfer onto nitrocellulose. D) Model of a potential actin nucleating complex. Bud6 is known to homo-dimerize, whereas Boi1 and Boi2 can either homo- or heterodimerize. The suggested interaction of a Boi1/2 dimer with a Bud6 dimer is experimentally not proven but supported by the observation that the deletion of Boi1 weakens the interaction between Boi2 and Bud6 and vice versa.

Bud6 nucleates together with the yeast formin Bni1 linear actin filaments (Graziano et al., 2011; Graziano et al., 2013). Full activity of Bni1 requires its association with a Rho-GTPase (Dong et al., 2003; Evangelista et al., 1997). Bud6 consists of a C-terminal Actin- and Bni1-binding domain and a N-terminal region of unknown function (Tu et al., 2012). Testing N_ub_-Boi1 and N_ub_-Boi2 against CRU fusions to the N and C-terminal fragments of Bud6 locate the binding sites for Boi1/2 to its N-terminal 364 residues (Fig. 2B). The GST-fusion to this fragment precipitates Boi1- and Boi2-GFP from yeast extracts, suggesting that the interactions between Boi1/2 and Bud6 are direct (Fig. 2C) (Glomb et al., 2019). A cartoon of a tentative actin nucleation complex at the cell cortex is given in Figure 2D.

Nba1 was reported to down-regulate the concentration of active Cdc42 during cytokinesis (Meitinger et al., 2014). To understand how Boi1/2 mediates the interaction between Bem1 and Nba1 we tested N- and C-terminal fragments of Boi1 and Boi2 as N_ub_ fusions against Nba1CRU. The assay identified the SH3 domains of both proteins as contact sites for Nba1 (Fig. 3A). Introducing single residue exchanges into the SH3 domains of Boi1 and Boi2 (Boi1_WK_, Boi1_PL_, Boi2_WK_, Boi2_PL_) that are known to generally disrupt the binding to their PxxP targets also abolished the interactions of the respective N_ub_ fusions with Nba1CRU (Fig. 3B) (Larson and Davidson, 2000). Testing a C-terminal fragment of Nba1 as N_ub_ fusion against Boi1- and Boi2CRU restricted the binding motif for both SH3 domains to a site between residues 256 and 501 of Nba1 (Fig. 3B, C). This region harbors a consensus-binding motif for both SH3 domains of Boi1 and Boi2 (Tonikian et al., 2009). Removing this PxxP site in Nba1 (Nba1_ΔPxxP_, Fig. 3C) impaired the interaction between the corresponding N_ub_-Nba1_ΔPxxP_ and the C_ub_ fusions of Boi1/2 (Fig. 3C). Consistently, N_ub_-Nba1_ΔPxxP_ also failed to interact with the CRU fusion of Bem1 but not with the CRU fusion of Nba1 (Fig. 3B, C). Plasmon resonance spectrometry determined the K_D_s between a fragment of Nba1 containing the PxxP site (6xHIS-Nba1_202-289_SNAP) and SH3_Boi1_ and SH3_Boi2_ to approximately 0,74 µM (n=3) and 1,97 µM (n=2) respectively (Fig. 3D).

**Figure 3.**
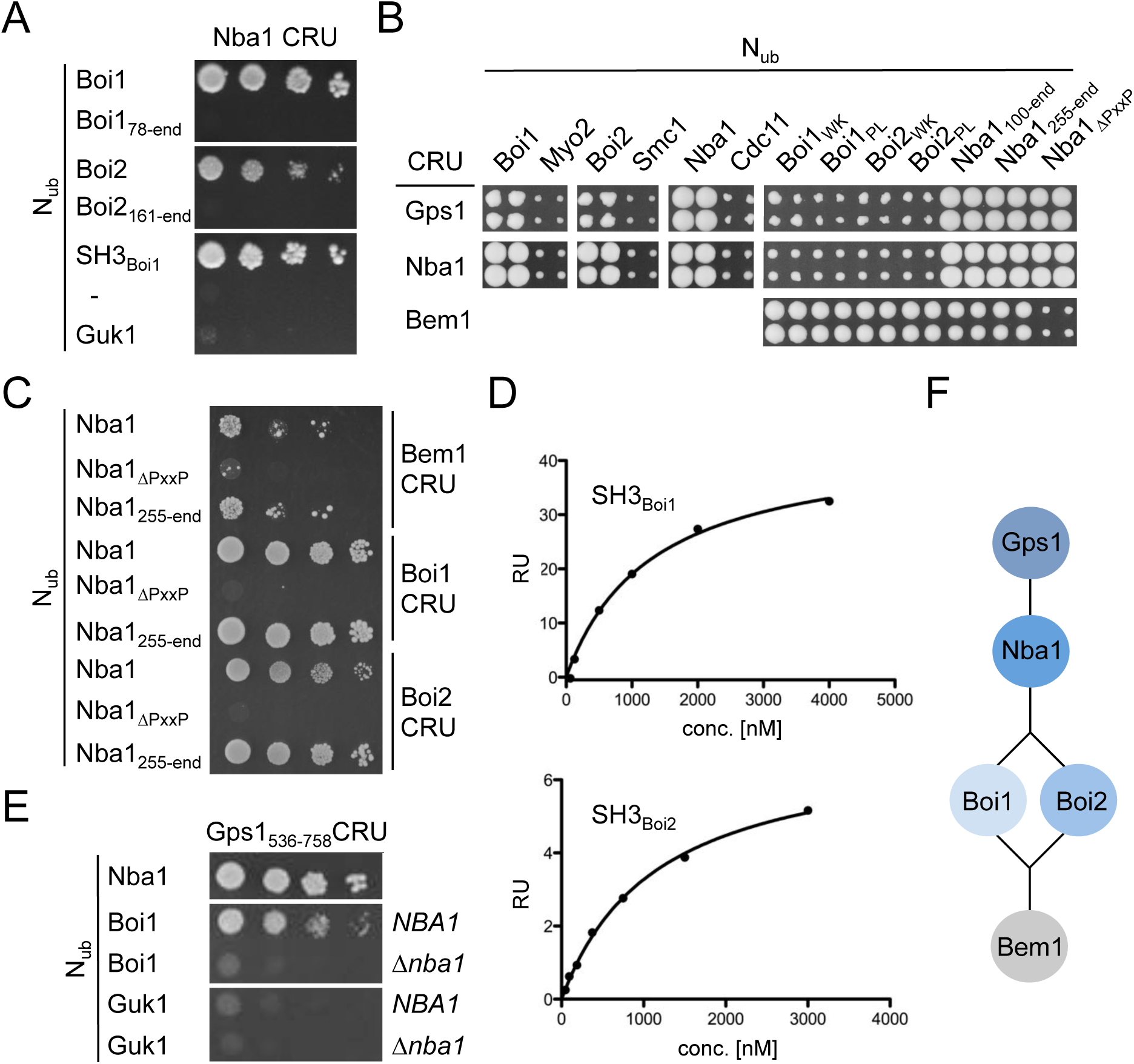
Characterization of the Bem1-Nba1 interaction state. A) Yeast cells co-expressing CRU fusions to Nba1 together with the indicated N_ub_ fusions were grown to an OD_600_ of 1 and spotted in 10-fold serial dilutions onto medium containing 5-FOA. Interactions are indicated by the growth of the yeast cells. N_ub_-Guk1 served as protein that is assumed not to interact with Nba1. B) a-yeast cells expressing the indicated CRU fusions were mated with alpha-yeast cells expressing the indicated N_ub_ fusion and tested for growth on medium containing FOA. Growing quadruplets indicate interaction between the corresponding N_ub_- and C_ub_ fusion. C) As in A) but with yeast cells co-expressing CRU fusions to Bem1, Boi1, or Boi2 together with N_ub_ fusions to Nba1 or its mutants. D) As in A) but with yeast cells containing or lacking *NBA1* and co-expressing a CRU fusion to a C-terminal fragment of Gps1 together with the indicated N_ub_ fusions. E) SPR analysis of the interaction between 6xHIS-Nba1_202-289_-SNAP and the immobilized GST-fusions to the SH3 domains of Boi1 or Boi2. Shown are representative plots of the SPR signals as response units (RU) against the concentrations of 6xHIS-Nba1_202-289_-SNAP including the fitting curves. F) Cartoon of the Nba1-Bem1 interaction state. The cartoon implies an indirect interaction between Bem1 and Gsp1. This interaction was not experimentally observed but inferred from the Nba1-dependent interaction between Boi1/2 and Gps1, and the Boi1/2-dependent interaction between Nba1 and Bem1, as well as the Gps1-dependent neck localization of Bem1.

Nba1 is attached to the bud neck through a direct interaction with Gps1 (Meitinger et al., 2014). Split-Ub analysis reproduced the interactions of Gps1CRU with the N_ub_-fusions of Nba1, Nba1_ΔPxxP_, and revealed novel interactions between Gps1CRU and N_ub_-Boi1/2 (Fig. 3B). Mutations in the SH3 domains of Boi1 and Boi2 abolished the interactions of the respective N_ub_ fusions with Gps1CRU (Fig. 3B) suggesting that Boi1/2 interact with Gps1 through Nba1. A C-terminal fragment of Gps1 still interacts as CRU fusion with N_ub_-Nba1 and N_ub_-Boi1 (Fig. 3E). The interaction between Gps1_537-758_CRU and N_ub_-Boi1 is lost in a *Δnba1*-strain thus supporting the existence of a protein complex that connects Bem1 with Gps1 through Boi1/2, and Nba1 (Fig. 3E, F).

Bem1 interacts with Cdc11 through its PB domain (Fig. 1). We have previously shown that Cdc11 interacts directly with Cdc24 and thus conclude that the Split-Ub-detected interaction between Cdc11 and Bem1 reflects the Bem1-Cdc24-Cdc11 complex (Chollet et al., 2019).

### Functional dissection of Bem1 interaction network

As central scaffold protein of the Cdc42 pathway Bem1 is thought to coordinate the activities of its members by bringing them into close spatial proximity. To define which combination of binding sites and -partners constitute the essential configurations of the Bem1-complex we first tested mutants and fragments for their ability to complement a deletion of *BEM1. BEM1* is not essential in each yeast strain but required for cell survival in the strain JD47 (Fig. 4A) (Dowell et al., 2010). *Δbem1*-cells can be rescued by the simultaneous deletion of the Cdc42 GAP Bem3 but not by the deletion of the Cdc42 GAP Bem2 (Fig. 4A) (Laan et al., 2015). The ability to rescue depends on Bem3’s GAP activity and a functional PH domain (Fig. S1). The CRIB domain of Gic2 (Gic2_PBD_) captures Cdc42_GTP_ and is used as RFP fusion to monitor active Cdc42 in living yeast cells (Atkins et al., 2013; Brown et al., 1997; Okada et al., 2013; Orlando et al., 2008). Overexpression of Gic2_PBD_ in *Δbem1 Δbem3*-cells eliminates the positive effect of the *BEM3* deletion on the cell survival of *Δbem1-*cells (Fig. 4A). The results imply that Gic2_PBD_ reduces the free pool of Cdc42_GTP_ at the cell cortex and that Bem1 is needed to stimulate the synthesis and/or to improve the effective use of this pool. A fragment of Bem1 (Bem1_145-551_) that covers SH3_b_CI, the PX- and the PB domain and thus keeps the majority of all detected interactions, complements *Δbem1*-cells (Figs. 4B, S2). This region can be further divided into two independently complementing fragments: The SH3_b_CI domain (Bem1_145-268_) with its binding sites for Cdc42 and for its PxxP ligands Ste20, Cla4, Boi1/2, and the C-terminal fragment containing the PX- and the PB domain (Bem1_268-551_) with its binding sites for Cdc24, lipids and the other PB-domain ligands (Figs. 1A, 4B, S2). Any further reduction of either fragment abolishes their ability to complement. The autonomy of the central SH3_b_CI-domain was unexpected. Single mutations that interrupt the binding of SH3_b_CI either to the PxxP ligands (SH3_bWK_CI) or to Cdc42_GTP_ (SH3_b_CI_ND_) interfere with the fragment’s ability to rescue *Δbem1* cells (Figs. 4D, S2) (Gorelik and Davidson, 2012; Yamaguchi et al., 2007).

**Figure 4.**
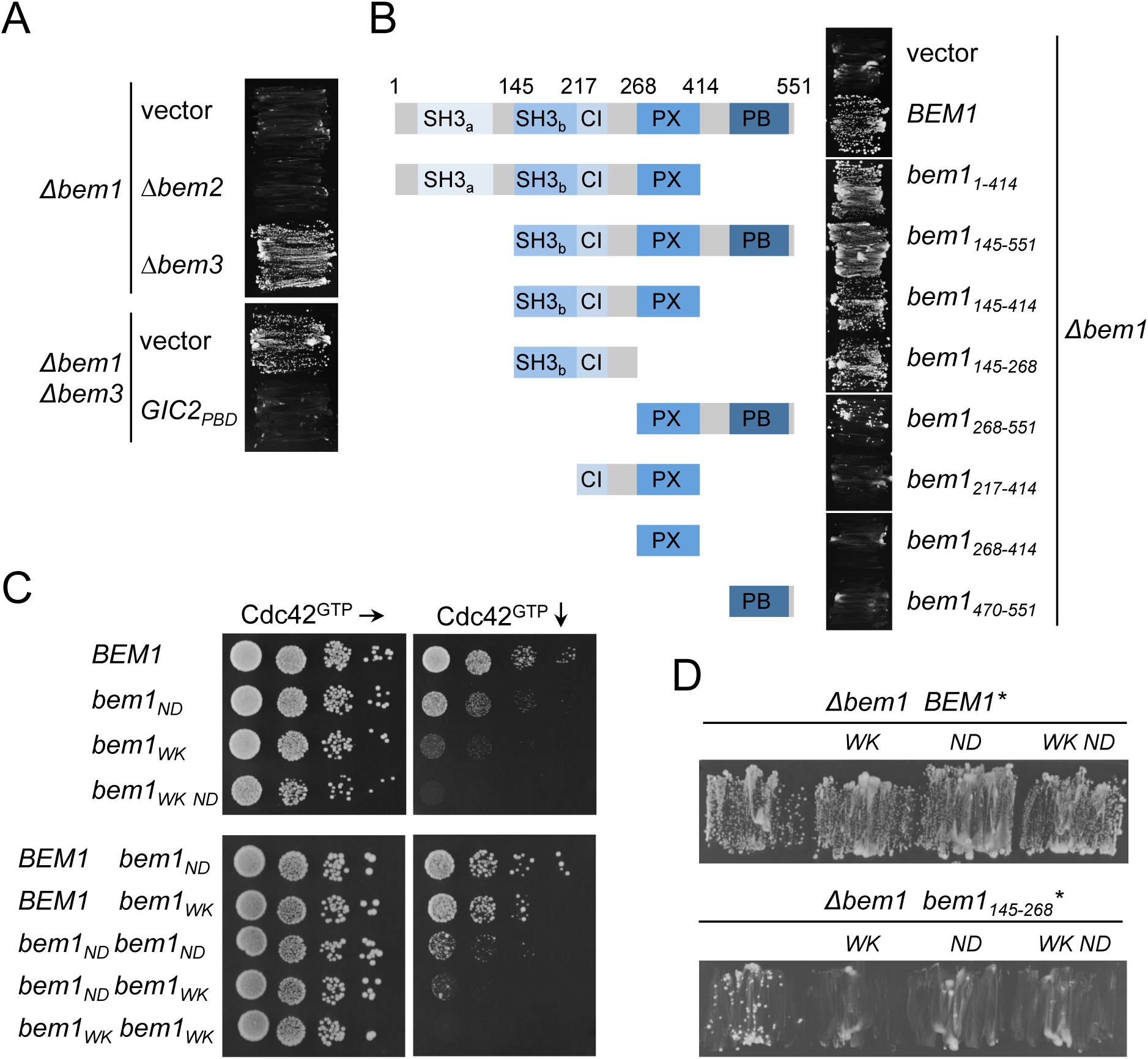
Bem1 contains two functionally independent regions. (A) *Δbem1*-cells carrying a vector-encoded *BEM1* and carrying an additional gene deletion as well as an empty vector, or a vector expressing Gic2_PBD_, were incubated on media selecting against the presence of the plasmid-encoded *BEM1*. (B) *Δbem1*-cells carrying a vector-encoded *BEM1* and a vector expressing Bem1 or the indicated fragments of *BEM1* (left panel) were incubated on medium selecting against the vector-encoded *BEM1* (right panel). (C) Haploid cells (upper panel), or diploid cells (lower panel) carrying the indicated alleles of *BEM1* and expressing Gic2_PBD_ were incubated in 10-fold serial dilutions on media moderately (left panels) or fully (right panels) expressing Gic2_PBD_. (D) *Δbem1*-cells carrying a vector-encoded *BEM1* and additionally expressing the full length *BEM1* with the indicated residue exchanges, or fragments of *BEM1* with the indicated residue exchanges, were incubated on medium selecting against the vector-encoded *BEM1*.

The existence of two independently complementing regions explains why none of the single interaction-interfering mutations in the SH3_b_CI domain or the deletion of the PB domain alone eliminate the functionality of the otherwise full-length Bem1 (Fig. 4B, C). A low concentration of active Cdc42 requires the presence of Bem1 for cell survival (Fig. 4A). We artificially reduced the pool of free Cdc42_GTP_ by expressing increasing amounts of Gic2_PBD_ in *bem1_WK_*-, *bem1_KA_*-, *bem1_ND_*- or *bem1_WK ND_*-cells (Fig. 4D). All tested interaction interfering mutations drastically decreased the tolerance of the cells towards Gic2_PBD_ overexpression. The allele *bem1_WK ND_* bearing both mutations in the SH3_b_CI domain confers a higher sensitivity than the singly mutated *bem1_WK_*- or *bem1_ND_*-allele. Overexpressing Bem3 and thus reducing Cdc42_GTP_ at the cortex by different means has a similar impact on these mutants (Fig. S1).

To test whether the closely spaced SH3_b_ and CI operate independently of each other, we introduced the WK and ND mutations in the different *BEM1* copies of the diploid genome to co-express Bem1_WK_ and Bem1_ND_ in one cell. The undiminished sensitivity of these cells toward Gic2_PBD_ overexpression suggests that both binding sites operate in cis and have to contact Cdc42_GTP_ and one of the SH3_b_-ligands at the same time (Fig. 4C).

### Functional annotation of Bem1-PAK interaction states

All four SH3_b_-ligands bind Cdc42_GTP_. The functional linkage between SH3b and CI suggest that Bem1 shuttles active Cdc42 from Cdc24 directly to these effectors. Which of the four known SH3_b_-interactions become critical under conditions of limited Cdc42_GTP_? None of the four SH3_b_-ligands are essential. Cells however do not tolerate the simultaneous loss of both PAKs, or of both Boi-proteins (Bender et al., 1996; Cvrckova et al., 1995). We introduced mutations in *CLA4* (*cla4_F15AAA/PPF451L_* = *cla4_PPAFL_*), *STE20* (*ste20_F470L P475T_ = ste20_FLPT_*), and *BOI1* (*boi1_Δ_*_PxxP_) that specifically reduce their affinities to SH3_b_ (Gorelik and Davidson, 2012; Bender et al., 1996; Kozubowski et al., 2008) (Fig. S3). The mutations in *STE20* and *CLA4* were selected to not impair their interaction with Nbp2, a further ligand of their PxxP motifs (Fig. S3) (Winters and Pryciak, 2005; Gorelik and Davidson, 2012; Hruby et al., 2011). Gic2_PBD_ overexpression killed cells lacking *CLA4* and the SH3_b_-binding motif of Ste20, or cells lacking *STE20* and the SH3_b_-binding motifs of Cla4, or cells co-expressing *cla4_PPAFL_* with *ste20_FLPT_* (Fig. 5A). Cells lacking *BOI2* and the Bem1-binding site in Boi1 were not affected by Gic2_PBD_ -overexpression (Fig. 5A). The PH domains of Boi1 (PH_Boi1_) and Boi2 (PH_Boi2_) bind Cdc42_GTP_ (Bender et al., 1996; Kustermann et al., 2017). Similar to Gic2_PBD_, overexpression of PH_Boi1_ kills cells lacking the physical connection of Bem1 to Ste20, and Cla4 but does not affect cells lacking the connection of Bem1 to Boi1/2 (Fig. S1D). A simultaneous overexpression of Cdc42 suppresses the toxic effect of PH_Boi1_ on *Δste20 cla4_PPAAFL_*-cells (Fig. S1E).

**Figure 5.**
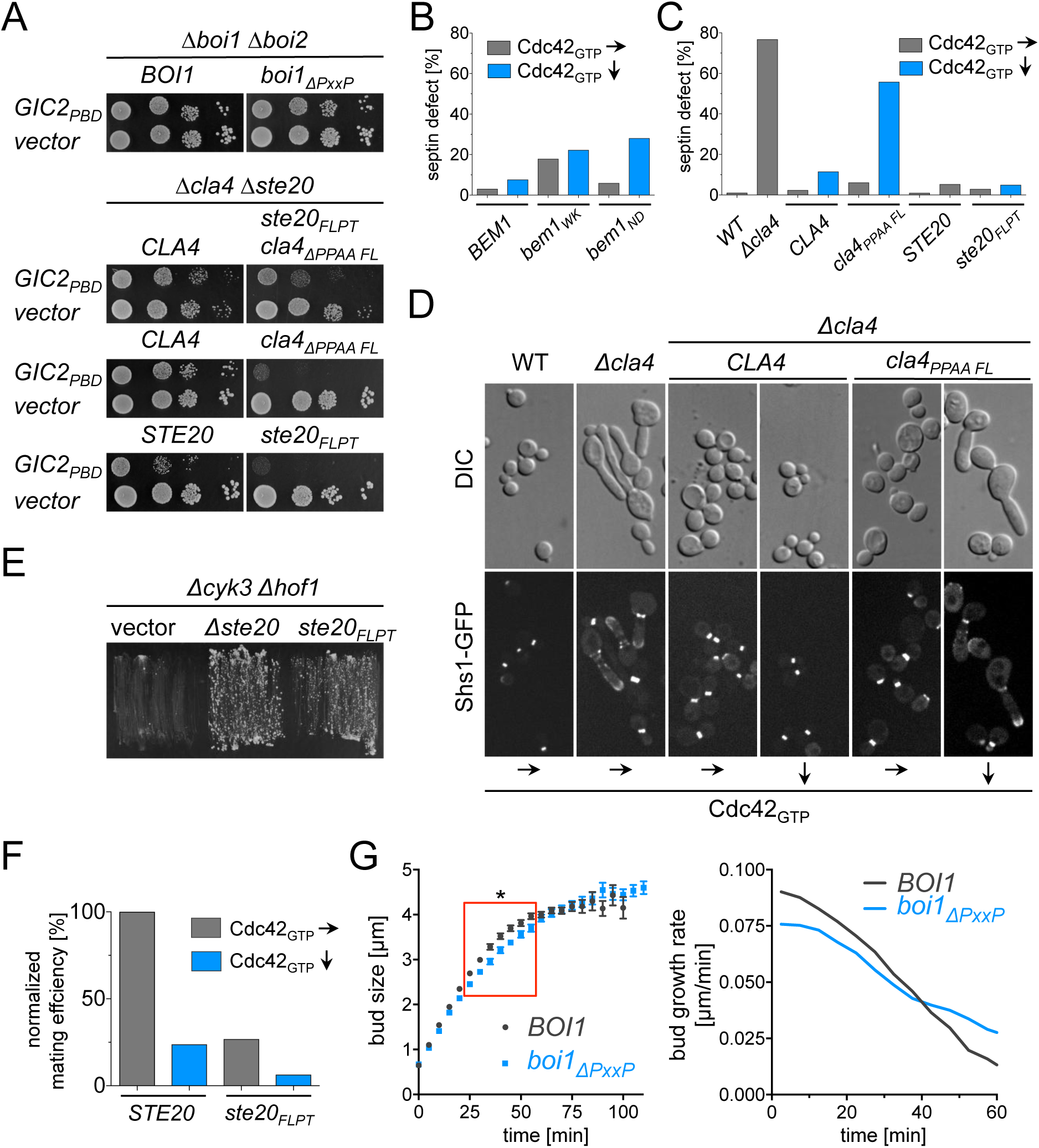
The connection between Bem1 and the PAKs is essential under limiting amounts of active Cdc42. (A) Yeast cells carrying the indicated alleles and either Gic2_PBD_ (upper panels) or an empty vector (lower panels) were spotted in 10 fold serial dilutions on medium inducing low (left panels) or high (right panels) expression of Gic2_PBD_. (B) Cells expressing a GFP fusion to Shs1 and carrying the indicated alleles of *BEM1,* and a vector expressing Gic2_PBD_ were incubated under conditions of low (70 µM Met, black bars) or high expression (0 Met, striped bars) of Gic2_PBD_. Cells (500<n<600) inspected under the fluorescence microscope were classified according to their native-like or abnormal distribution of the septin Shs1-GFP. (C) As in (B) but with cells (500<n<600) carrying the indicated alleles of *STE20* or *CLA4*. (D) The morphologies and the Shs1-GFP distributions of the cells of (C) were visualized by DIC and fluorescence microscopy under conditions of low and high Gic2_PBD_ expression. (E) *Δcyk3 Δhof1* cells expressing *HOF1* from an extra-chromosomal vector and carrying the indicated alleles of *STE20* or an empty vector were incubated on media selecting against the *HOF1* containing vector. (F) a- and alpha-cells containing the same alleles of *STE20* and expressing Gic2_PBD_ were incubated together in media inducing low (70 µM Met, black bars) or high expression (0 Met, grey bars) of Gic2_PBD_ and plated on media selecting for the presence of diploid cells. (G) The bud length of cells (n=38; SEM) carrying the indicated alleles of *BOI1* and co-expressing Bem1-GFP and Shs1-mCHERRY were measured every five min starting with a bud length of 0.65 µm and ending at the time point of septin splitting (left panel). Red box indicates the time window with a significant difference between *Δboi2 BOI1-* and *Δboi2 boi1_ΔPxxP_*-cells. Right panel shows the first derivatives of the growth curves of the left panel and compares the rate of bud growth in µm/min. Scale bar indicates 3µM.

Cells without Cla4 or its kinase activity do not correctly assemble septins at the incipient bud site and as a consequence display mis-localized septin patches and elongated buds (Weiss et al., 2000; Holly and Blumer, 1999). These phenotypes were recapitulated in *bem1_WK_*- or in *bem1_ND_*-cells, or in *cla4_PPAAFL_*-cells upon overexpression of Gic2_PBD_ (Fig. 5). In contrast, Gic2_PBD_ overexpression did not affect morphology or the septin cytoskeleton of *ste20_FLPT_*- or *Δboi2 boi1_ΔPxxP_*-cells (Figs. 5C, D, S4). The experiments prove that the Cla4-Bem1-Cdc42_GTP_ complex is important during incipient bud site- and septin-assembly, whereas the Ste20-Bem1-Cdc42_GTP_ or the Boi1/2-Bem1-Cdc42_GTP_ complexes do not measurably contribute to this activity.

A deletion of *STE20* rescues a strain that is arrested at cytokinesis by the simultaneous loss of the cytokinesis factors Hof1 and Cyk3 (Onishi et al., 2013). It is speculated that Cdc42 inhibits secondary septum (ss) formation through activation of Ste20. The loss of Ste20 might prematurely activate ss-formation thus compensating for the loss of the primary septum. We confirmed and extended this observation by showing that also the *ste20_FLPT_* allele rescues *Δhof1Δcyk3* cells (Fig. 5E). Accordingly, the Ste20-Bem1 complex is functionally relevant during cell separation. The same interaction is important for the fusion of cells during mating (Fig. 5F) (Winters and Pryciak, 2005). Again, the reduction of the freely available pool of Cdc42_GTP_ potentiates the effect of the *ste20_FLPT_* allele (Fig. 5F).

The Boi1 proteins stimulate the fusion of secretory vesicles with the plasma membrane (Kustermann et al., 2017; Masgrau et al., 2017). To test whether the Bem1-Boi1/2 complex contributes to this activity we compared the kinetics of tip growth of *Δboi2*-cells either containing or lacking the binding site of Boi1 for Bem1. By monitoring the distance from bud neck (Shs1-mCherry) to bud tip (Bem1-GFP) we observed for *Δboi2 boi1*_ΔPxxP_-cells a reduced rate of tip elongation during the first 40 min into bud growth. Consequently, *Δboi2 boi1_ΔPxxP_*-cells needed more time to reach the critical size for cell separation (Fig. 5G).

### The PAK kinases do not anchor Bem1 at the bud tip

Bem1 localizes at the growth zone of the cell to effectively shuttle Cdc42 to its effectors. Before identifying the interactions that determine the cell cycle-specific distribution of Bem1 we first investigated the cellular localization of its different, functionally characterized fragments. SH3_b_CI-GFP is the minimal GFP-labeled fragment of Bem1 that fully recapitulates the cellular distribution of the full-length protein (Fig. 6A). The cortical targeting of SH3_b_CI requires the ligands of SH3_b_ but not Cdc42_GTP_, as SH3_b_CI_ND_-GFP is still concentrated at bud neck and tip, whereas SH3_bWK_CI-GFP stays cytosolic throughout the cell cycle (Fig. 6A). To find out which of the four SH3_b_-ligands influences its distribution we expressed SH3_b_-CI-GFP in cells each lacking a specific SH3_b_ binding site. The analysis identified Boi1/2 as the exclusive receptor for SH3_b_CI at cortex and bud neck (Fig. 6B).

**Figure 6.**
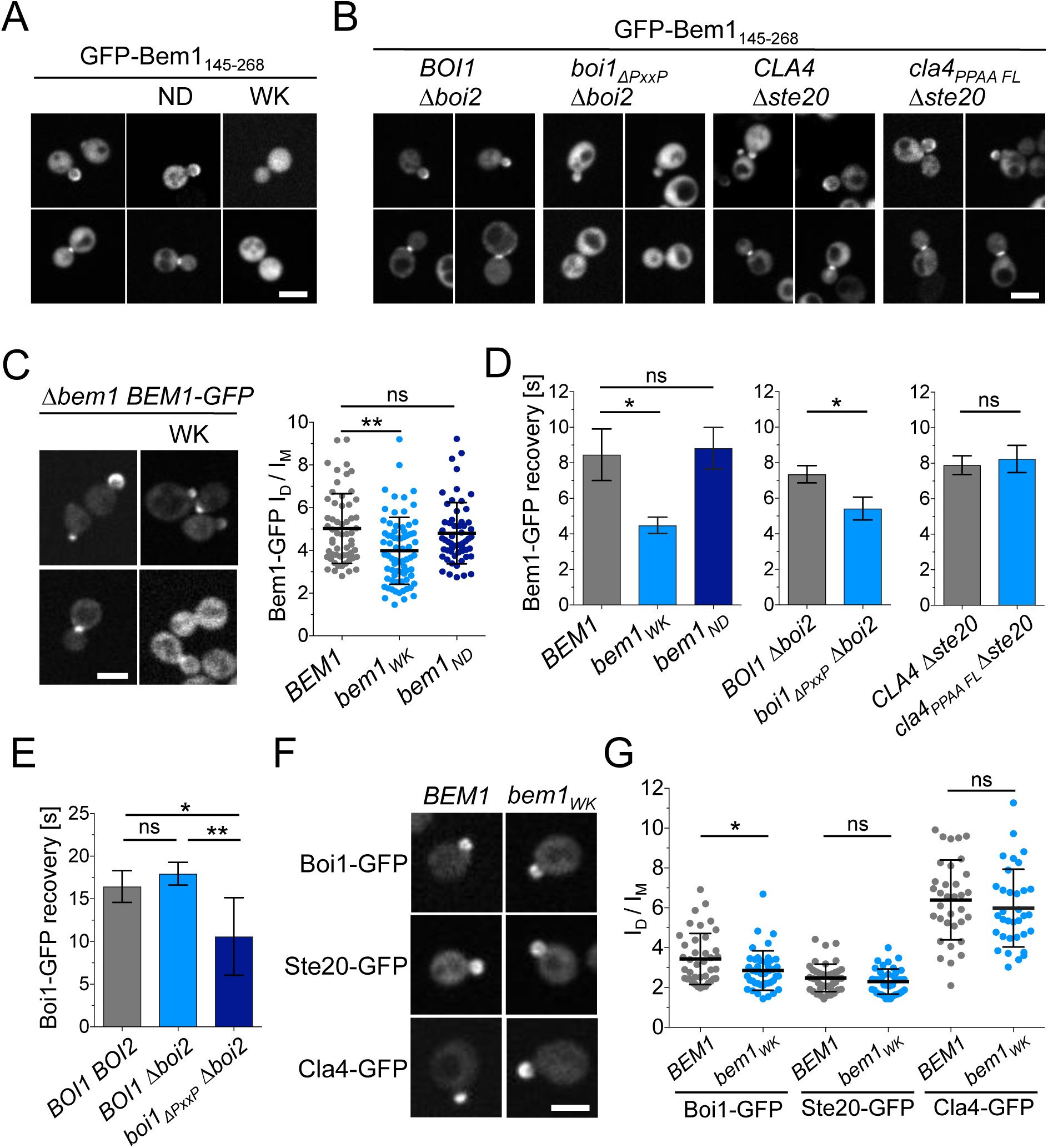
Boi1 and Boi2 localize Bem1-Cdc24 at bud tip and neck. (A) Cells expressing GFP-Bem1_145-268_ (left panel) and carrying the ND (middle panel) or WK (right panel) exchange were inspected by fluorescence microscopy. The Bem1-fragment carrying the WK exchange does not localize at bud tip and neck any longer. (B) Cells of the indicated genotypes expressing GFP-Bem1_145-268_ were inspected by fluorescence microscopy (left panel). Only *Δboi2 boi1_ΔPxxP_*-cells show a clear misdistribution of GFP-Bem1_145-268_. (C) Quantification of the ratio of the Bem1-GFP-(n= 58), Bem1_WK_-GFP-(n= 72), and Bem1_ND_-GFP (n=57) intensities in bud and mother cell reveal a higher fraction of Bem1_WK_-GFP in the mother cell. (D) Left panel: Half-times of fluorescence recovery after photo-bleaching the bud of the cells expressing Bem1-GFP (n=11) Bem1_WK_-GFP (n= 18) or Bem1_ND_-GFP (n= 24). Middle panel: Half-times of fluorescence recovery after photo-bleaching the bud of *Δboi2 BOI1* cells (n=20) or *Δboi2 boi1_ΔPxxP_-*cells (n=16) expressing Bem1 GFP. Right panel: Half-times of fluorescence recovery after photo-bleaching the bud of *Δste20 CLA4*-cells (n= 16) or *Δste20 cla4_PPAAFL_*-cells (n=14) expressing Bem1-GFP. (E) Half-times of fluorescence recovery after photo-bleaching the bud of *BOI1 BOI2*-cells (n=23), *Δboi2 BOI1*-cells (n=24), or *Δboi2 boi1_ΔPxxP_-*cells (n=22) expressing GFP fusions to the *BOI1* alleles. (F) Upper panel: Cells of the indicated genotypes and expressing GFP fusions to Boi1 (left panel), Ste20 (middle panel), and Cla4 (right panel), were inspected by fluorescence microscopy. Lower panel: The ratios of the fluorescence intensities of bud and mother cells were quantified in *BEM1*-(n= 40) and *bem1_WK_*-cells (n=43) expressing Boi1-GFP, in *BEM1*-(n= 43) and *bem1_WK_*-cells (n=40) expressing Ste20-GFP, and in *BEM1*-(n=34) and *bem1_WK_*-cells (n= 34) expressing Cla4-GFP. ns= not significant. * = p <0.05, ** = p <0.01. Scale bars indicate 3µM.

Boi1/2 are necessary and sufficient to attach also the full length Bem1 to the bud neck (Fig. 6C, see also Fig. 9) whereas the fluorescence signal of Bem1_WK_-GFP is only reduced but not abolished at the tip of small and large buds (Fig. 6C). To obtain a quantitative measure of tip affinity we compared the FRAPs between the cortex-localized Bem1-GFP, Bem1_WK_-GFP, and Bem1_ND_-GFP (Fig. 6D). The halftime of recovery was not changed by the N253D mutation in Bem1 (t_1/2_ = 8.82 ±1.12 sec.) whereas the W192K exchange reduces t_1/2_ to 4.48 ± 0.46 sec (Fig. 6D). A comparable reduction in t_1/2_ is observed when the FRAPs of Bem1-GFP are compared between *Δboi2*-cells (t_1/2_ = 7.35 ± 0.49 sec) and *Δboi2*-cells lacking in addition the Bem1-binding sites in Boi1 (t_1/2_ = 5.42 ± 0,49 sec) (Fig. 6D). The FRAP of the cortex-localized Bem1 was not changed in cells where the interaction between Bem1 and the PAKs were eliminated (*Δste20 cla4_PPAAFL_*) (Fig. 6D).

As expected for the cortical anchor of Bem1, Boi1 has slower FRAP than Bem1 (Fig. 6E). Boi1 and Boi2 are recruited to the bud cortex mainly through their Cdc42_GTP_- and lipid-binding PH domains. Bem1 contains in addition to its interaction with Boi1/2 multiple phospholipid-binding sites (Hallett et al., 2002; Meca et al., 2019). The physical connection between Bem1 and Boi1/2 might thus reciprocally influence the strength and selectivity of their cortex attachment. Indeed, Boi1_ΔPxxP_-GFP carrying a mutated Bem1-binding site was significantly more mobile than the native protein (t_1/2_ = 16,43 ± 1,86 sec versus t_1/2_ = 10.58 ± 0.97 sec) (Fig. 6E). In accordance, Boi1-GFP was also less focused at the bud cortex of *bem1_WK_*-cells, whereas *bem1_WK_* did not detectably influence the cortical localization of Ste20-GFP or Cla4-GFP (Fig. 6F) (Winters and Pryciak, 2005). The experiments provide strong evidence for a cooperative recruitment of Boi1/2 and Bem1 to the bud cortex. The cortex-localized PAK kinases do not measurably contribute to Bem1’s distribution during the cell cycle.

### Nba1 is the primary attachment site for Boi1/2-Bem1 at the bud neck

Bem1 leaves the cortex during mitosis and arrives at the bud neck shortly before the acto-myosin ring contraction is completed (Fig. S5). With a FRAPt_1/2_ of 6 s (t_1/2_ = 6.01 ± 0.63; n=23) Bem1 is slightly more mobile at the neck than at the bud tip. Boi1/2 link Bem1 and Cdc24 to the neck (Fig. 6B, 7C). Mutations in the SH3 domains of Boi1 (Boi1_WK_), and Boi2 (Boi2_WK_) remove the Boi proteins from the neck (Fig. 7C) (Hallett et al., 2002). Nba1 is a potential docking site for Boi1/2 as its binds to both SH3 domains and arrives at the neck at roughly the same time as Bem1 (Fig. 3) (Meitinger et al., 2014). Accordingly, a deletion of *NBA1* or of its Boi1/2-binding site (Nba1_ΔPxxP_) removes Boi2-GFP completely, and 55% of Boi1-GFP from the neck (Fig. 7B). The Nba1-mediated interaction between Gps1 and Boi1/2 suggests that Gps1 might anchor the Nba1-Boi1/2-Bem1-Cdc24 complex at the cell division site (Fig. 3F). *Δgps1*-cells lack Nba1-GFP at the neck, and reduce neck localizations of Boi1-GFP and Boi2-GFP to a similar extent as *Δnba1*-cells (Fig. 7A, B) (Meitinger et al., 2014). We conclude that the entire Boi2 is attached by Nba1-Gps1, whereas 45% of Boi1 are anchored by still unknown binding partner(s). The bud neck localization of the isolated SH3_Boi1_ or SH3_Boi2_ mirror the SH3-dependencies of the corresponding full-length proteins (Fig. 7B). However, t_1/2_ of FRAP of SH3_Boi1_-GFP is significantly shorter than t_1/2_ of the full-length Boi1-GFP (t_1/2_ = 0.7 s versus t_1/2_ = 12 s; Fig. 7D). The difference points to regions outside of SH3_Boi1_ that contribute to its binding to neck. The binding site for Bem1 is a reasonable candidate as a direct interaction between Bem1 and Nba1 was reported (Meitinger et al., 2014). However, Boi1_ΔPxxP_-GFP lacking the binding site to Bem1 stills displays a t_1/2_ of 10 s at the bud neck that is very similar to the t_1/2_ of FRAP of the wild type protein (Fig. 7D). This observation supports the conclusion that Bem1 recruitment to the neck is distinct from its synergistic recruitment at the cortex.

**Figure 7.**
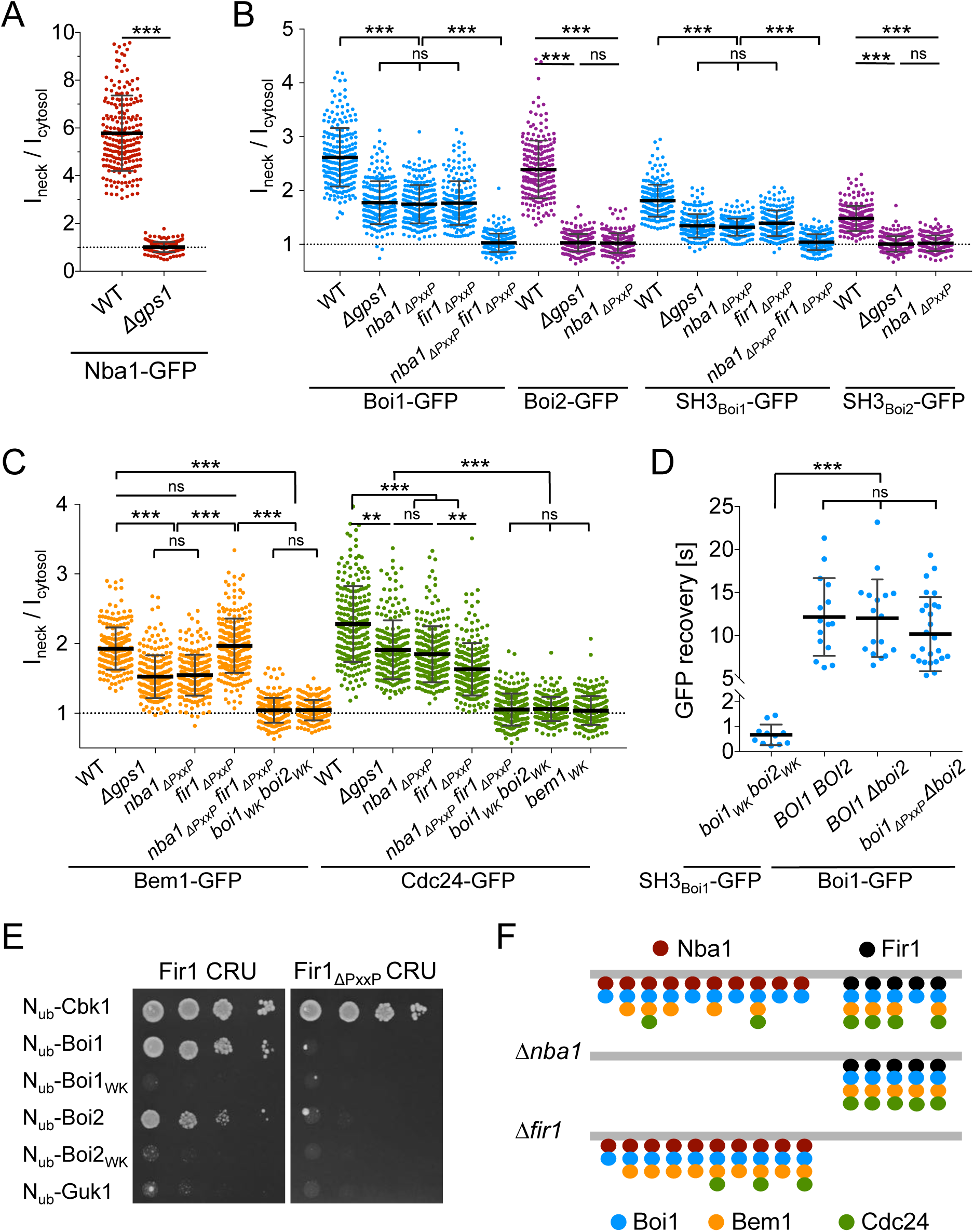
Two receptor systems attach Bem1-Cdc24 to the bud neck. (A) Ratios of bud neck to cytosolic fluorescence intensities of Nba1-GFP in wild type- and *Δgps1-*cells. (B) As in (A) but in cells of the indicated genotypes expressing Boi1-GFP, Boi2-GFP, SH3_Boi1_-GFP, SH3_Boi2_-GFP. (C) As in (A) but in cells of the indicated genotypes expressing Bem1-GFP, or Cdc24-GFP. (D) Half time of fluorescence recovery after photo bleaching the bud neck of *boi1_WK_ boi2_WK_*-cells expressing SH3_Boi1_-GFP (n=), of cells expressing Boi1-GFP (n= 15), of *Δboi2*-cells expressing Boi1-GFP (n= 17), and of *Δboi2*-cells expressing Boi1*_ΔPxxP_*-GFP (n= 27). (E) Split Ubiquitin analysis of cells expressing Fir1CRU or Fir1_ΔPxxP_CRU together with N_ub_ fusions to Boi1, Boi2, Boi1_WK_, Boi2_WK_, CbK1 as positive control, and Guk1 as non-interacting protein. Cells were grown to an OD_600_ of 1 and spotted in 10-fold serial dilutions onto medium containing 5-FOA. Interactions are indicated by the growth of the yeast cells. (F) Cartoon of the Bem1-Cdc24 complex at the bud neck. Nba1 and Fir 1 recruit Boi1 through its SH3 domain to the bud neck (for simplicity only Boi1 is shown), which recruits Bem1 through its SH3b domain. Cdc24 is attached to Bem1 through its PB domain. The concentration of the proteins at the neck is decreasing in the order of their appearance. To explain the distribution of the proteins in wild type (upper panel), *Δnba1* (middle panel), and *Δfir1* (lower panel) strains we assumed that Nba1 outnumbers Fir1 at the neck, that all Nba1 and Fir1 are bound by Boi1/2, that the number of Nba1 equals the number of Bem1, and that Nba1 reduces the affinity between Cdc24 and Bem1.

A complete loss of Bem1-GFP- or Cdc24-GFP intensity at the neck occurs only in cells expressing the SH3 mutations in both Boi proteins (*boi1_WK_ boi2_WK,_* Fig. 7C). In contrast, *nba1_ΔPxxP_*-, or *Δgps1*-cells still keep approximately 69% of Cdc24-GFP and 58% of Bem1-GFP at the neck (Fig. 7C)

### Fir1 is the alternative anchor for Boi1-Bem1-Cdc24 at the neck

Which receptor couples the unaccounted fraction of the Cdc24-Bem1-Boi1/2 complex to the cell division site? Fir1 relocates at the end of cytokinesis to the bud neck (Brace et al., 2019). A two-hybrid analysis reported the interaction between Boi1 and Fir1 and predicted a binding site for the SH3 domains of Boi1 and Boi2 between residues 523 and 533 of Fir1 (Tonikian et al., 2009). A Split-Ubiquitin analysis confirmed the interaction between Boi1/2 and Fir1 and could further demonstrate that the interactions depends on the functional SH3_Boi1_ or SH3_Boi2_ and the potential SH3_Boi1/2_-binding motif in Fir1 (Fig. 7E). To measure the influence of Fir1 on the bud neck localization of Boi1, SH3_Boi1_, and the Cdc24-Bem1 complex we introduced the corresponding GFP fusions in strains lacking *FIR1* (*Δfir1*), lacking the Boi1-binding motif in *FIR1 (fir1_ΔPxxP_*), or in cells lacking the motifs in *FIR1* and *NBA1 (nba1_Δpxxp_ fir1_Δpxxp_)* (Fig. 7B, C). Quantifying the intensities of the GFP signals proved that Fir1 recruits the Boi1-Bem1-Cdc24 complex independently of Nba1 to the bud neck. Boi1 is equally distributed between Nba1 and Fir1 (Fig. 7B). In contrast, the distribution of the Bem1-Cdc24 complex between Nba1 and Fir1 is not symmetrical (Fig. 7C). Proportional more of Cdc24 is anchored through Fir1 than through Nba1, whereas the amount of neck-anchored Bem1 does not change upon removal of the Boi1-binding site in Fir1 (Fig. 7C). We conclude that the Nba1- and Fir1-bound Boi1 and Boi2 are in excess over Bem1 at the neck. By assuming that the Nba1-Boi1-Bem1 complex binds Cdc24 less well than the Fir1-Boi1-Bem1 complex, we can explain why interrupting the interaction between Boi1 and Fir1 has a larger impact on the amount of neck-localized Cdc24 than interrupting the interaction between Boi1/2 and Nba1 (Fig. 7C). This assumption is strengthened by the finding that Nba1 interferes with Bem1-Cdc24 complex formation *in vitro* (Meitinger et al., 2014).

### Temporal dissection of the Bem1-interaction network

Scaffolds concentrate different activities at a certain time to a specific site. Split-Ub analysis provides a static projection of a measured interaction from all cell cycle stages. To correlate a certain interaction with a cell cycle specific function or localization we had to resolve this static projection into its successive temporal layers. We thus characterized the time dependency of a subset of Bem1 interactions through SPLIFF analysis (Moreno et al., 2013). The inducible *P_MET17_* promoter was integrated in front of the chromosomal Bem1CCG to obtain enough fluorescence for a one cell cycle-lasting time-lapse analysis. With exception of N_ub_-Exo70 and N_ub_-Cdc42, the *P_CUP1_* promoters of the genomically integrated N_ub_ fusions were replaced by their respective native promoters (Fig. S5C). We also included N_ub_-Rsr1 in our analysis as Rsr1 is known to bind to Bem1 and Cdc24 and as N_ub_-Rsr1 generates under its native expression levels a strong interaction signal with Bem1CRU (Fig. S5C) (Park et al., 1997).

After mating of the N_ub_- and Bem1CCG fusion protein-expressing cells, Bem1CCG accumulates at the site of cell fusion (PCDI), relocated to the presumptive bud site shortly before bud growth and stayed at the bud cortex below the plasma membrane till G2/mitosis (PCDII) to finally reappear at the bud neck shortly before contraction of the acto-myosin ring was completed (PCDIII) (Fig. S5). Green and red fluorescence were measured at PCDI and II every five and at the PCDIII every two minutes. The fluorescence intensities were plotted as percentage of N_ub_-induced conversion of Bem1CCG to Bem1CC against time after cell fusion (Fig. 8, Table S1). A net increase of conversion after two (PCDII) or three (PCDIII) consecutive time points was taken as evidence for the presence of the interaction during this time interval. No increase or a decrease in the relative amount of conversion was considered as absence of interaction. The interaction partners of Bem1 fall into three categories: Ste20, Cdc24, Boi1, Boi2, Cdc42 interact with Bem1 during all three phases (Fig. 8A). Cla4 interacts with Bem1 only during PCDI and II. Rsr1, Bud6, Rga2, Exo70 and Bem1 interact with Bem1 only during bud formation and growth (PCDII), and Nba1 interacts with Bem1 exclusively during cytokinesis (PCDIII) (Fig. 8). We can further differentiate between Bem1 interactions that last through the entire PCDII (Bem1, Boi1, Exo70, Cdc42, Cdc24) and those that are detectable in small buds only (Bud6, Rsr1) (Fig. 8). The interaction between Bem1 and Rga2 stalls during bud formation and picks up after 10 min into bud growth to continue as long as Bem1 remains at the cortex (Fig. 8). The interaction signals between Bem1 and Boi1, Boi2, Cal4 and Ste20 reach a plateau after approximately 20 min into bud growth. The slight increase of conversion was considered as sign of a continuing yet diminished interaction between Bem1 and Boi1 during the remaining phase of bud growth (Fig. 8). The decrease in the ratio of converted Bem1CCG in the N_ub_-Ste20, N_ub_-Cla4 and N_ub_-Boi2 expressing cells might already indicate a loss of interaction between Bem1 and the N_ub_-fusions during the transition from bud growth to mitosis. However, it has to be noted that conversion ratios at or above 80% are very difficult to interpret and that no increase or a slight decrease do not necessarily have to reflect absence of interaction. During cytokinesis only Boi1/2, Ste20, Cdc24, Cdc42 and Nba1 are detectably associated with Bem1 (Fig. 8). Bem1 clearly distinguishes between Ste20 and Cla4 during abscission.

**Figure 8.**
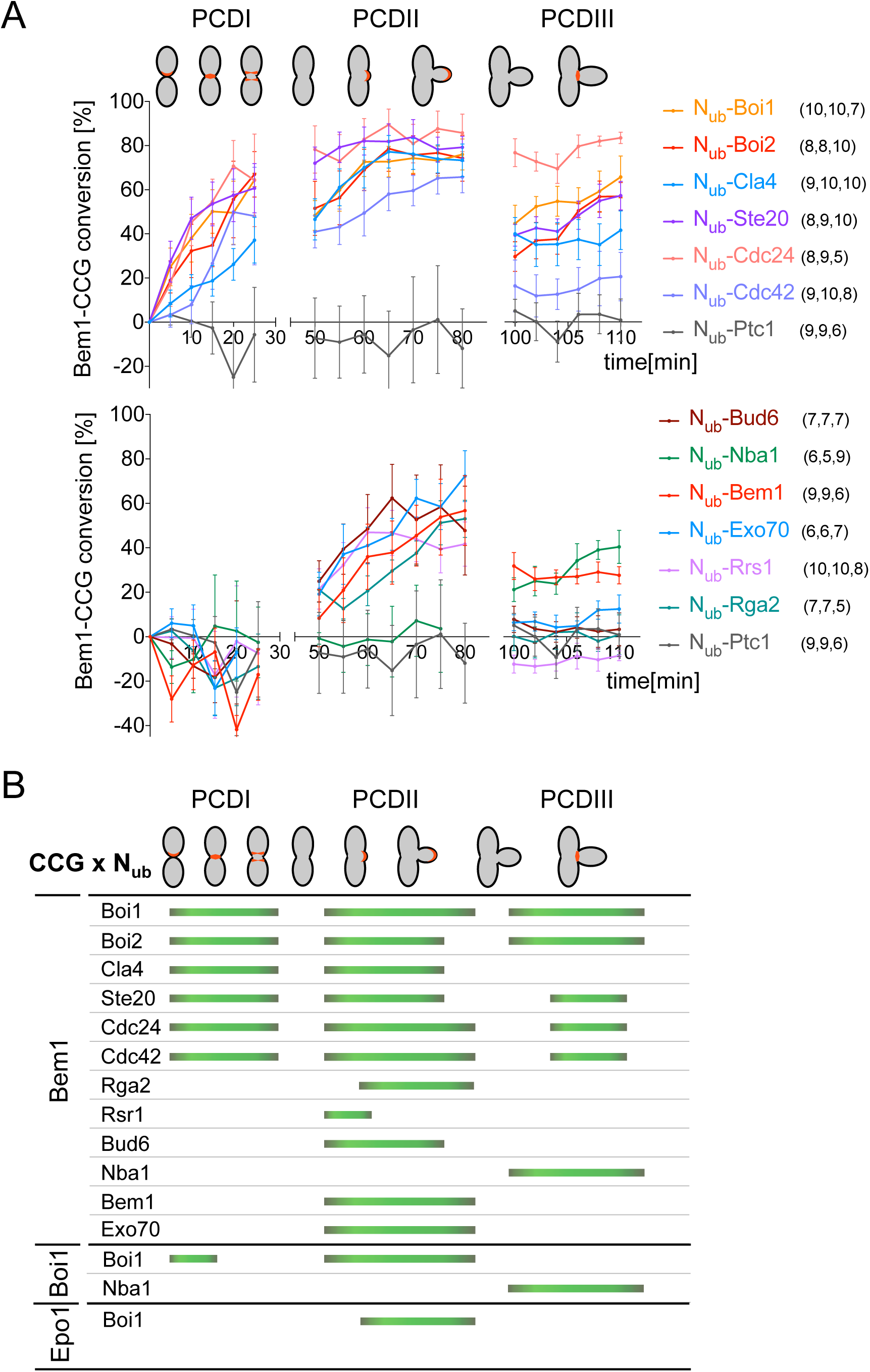
Time resolved analysis of Bem1 interaction states. SPLIFF analysis of yeast zygotes formed by the fusion of a-cells expressing Bem1CCG and alpha-cells expressing the indicated N_ub_-fusions. Plotted are the conversions of Bem1CCG to BemCC (in percent) over time. GFP-and mCherry fluorescence intensities were measured at sites of polarized Bem1 locations as indicated in red in the cartoons in the upper panel. PCDI (Polar cortical domain I) refers to a layer between the two cells after fusion has occurred, PCDII refers to the incipient bud site and to the bud tip during bud growth, and PCDIII refers to the site of cell separation during abscission. Measurements were taken every 5 min during PCDI/II and every 2 min during PCDIII. The numbers in brackets behind the N_ub_ fusions indicate the number of cells measured for PCDI/II and PCDIII. Error bars indicate SEM. (B) Interaction profiles for Bem1, Boi1, and Epo1. Green bars encompass the times where interactions between the respective CCG and N_ub_ fusions were detected. Interactions were indicated when conversion increased over three consecutive time points (see Results and Discussion). The profiles for Boi1CCG and Epo1CCG were derived from Figure 9.

### SH3_Boi1_ can switch between interaction partners during the cell cycle

The detection of the Bem1-Bem1 interaction requires the oligomerization of the Boi-proteins, whereas the proximity between Bem1 and Nba1 is mediated by the simultaneous interactions of both proteins with Boi1/2 (Figs. 2, 3).

We tested the consistency of our SPLIFF analysis by measuring Boi1CCG against N_ub_-Boi1 and N_ub_-Nba1. In agreement with the time dependency of Bem1 oligomerization and the formation of the Nba1-Bem1 complex, Boi1CCG was converted to Boi1CC by N_ub_-Boi1 during bud growth and not during cytokinesis whereas Boi1CCG was converted by N_ub_-Nba1 only during cytokinesis (Figs. 8B, 9C Table S1).

**Figure 9.**
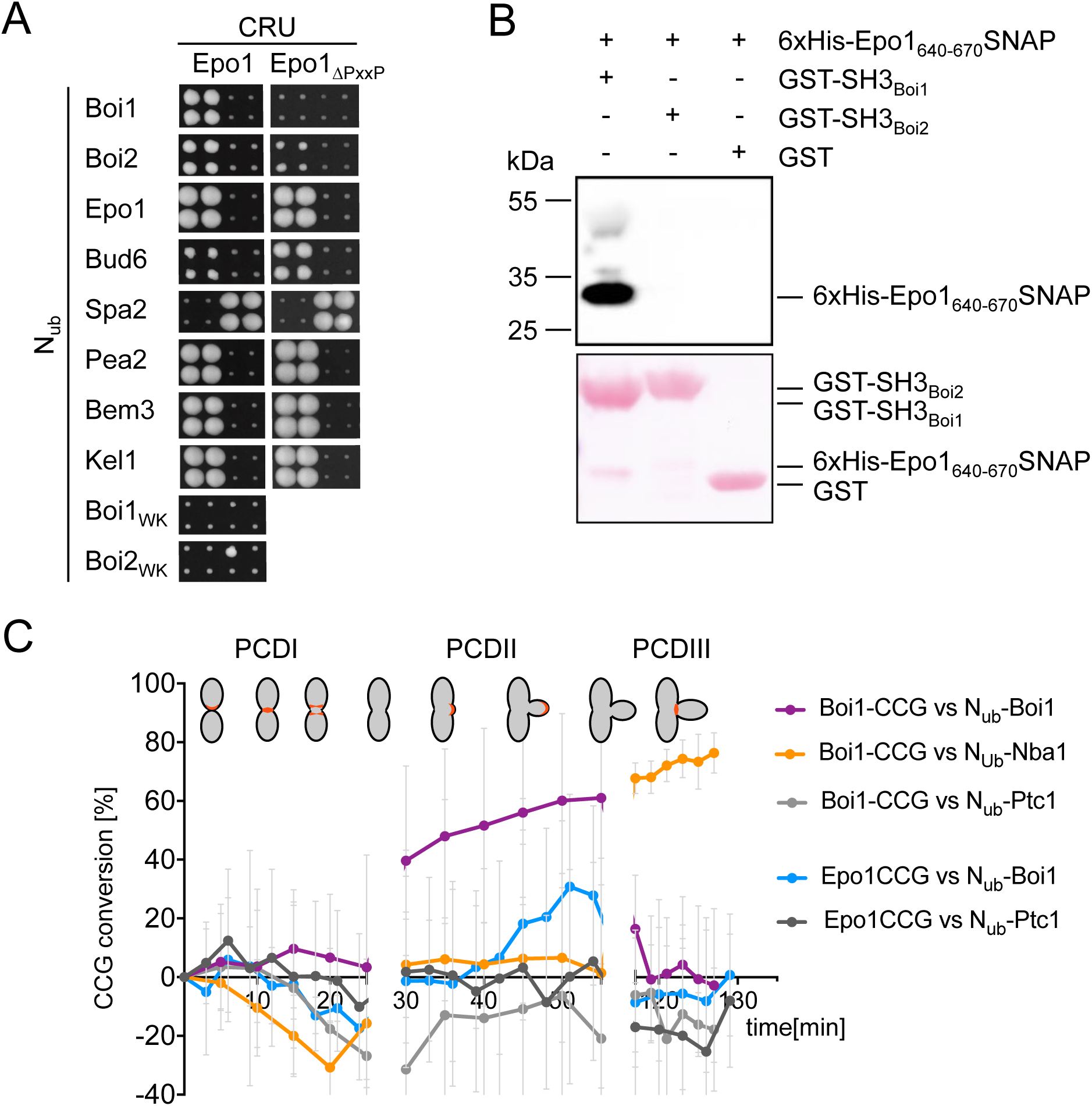
Epo1 interacts with Boi1 in large budded cells. (A) As in (Fig. 1A) but with cells expressing Epo1CRU or Epo1_Δ654-661_CRU. (B) Extracts of *E.coli*-cells expressing 6xHIS SNAP-tag fusion to Epo1_640-670_ were incubated with GST-Boi1-, GST-Boi2- or GST-coupled beads. Beads were eluted with glutathione and eluates were separated on SDS-PAGE, transferred on nitrocellulose, and stained with Ponceau (lower panel) and subsequently with anti-His antibodies (upper panel). (C) SPLIFF analysis of Epo1CCG and Boi1CCG as in Fig. 8. a-cells expressing Epo1CCG or Boi1 CCG were mated with alpha cells expressing N_ub_-Boi1, N_ub_-Ptc1 and additionally N_ub_-Nba1 in case of Boi1CCG cells. Shown are the calculated fractions of converted CCG-fusions in cells expressing the indicated N_ub_ fusions against time (7<n<13 independent matings; error bars, s.e.).

Besides binding to Nba1 and Fir1, SH3_Boi1/2_ interact with additional proteins and might expand the influence of the Bem1-Cdc24 complex to further processes (Kustermann et al., 2017; Tonikian et al., 2009). To test whether other SH3_Boi_ -interactions are also cell cycle specific we investigated the interaction to one of its potential SH3_Boi1_ ligands in detail. Epo1 links the cortical endoplasmic reticulum to the polarisome and was shown to bind Boi1 and Boi2 (Neller et al., 2015). Mutations that inactivate SH3_Boi1_, or a mutation of the predicted Boi1-binding motif in Epo1 abolishes the interaction between both proteins (Fig. 9A). A pull down of this binding motif with a GST-fusion to SH3_Boi1_ proves its direct interaction (Fig. 9B). Epo1 and Nba1 thus compete for the same binding site in Boi1. SPLIFF picks up the interaction between N_ub_-Boi1 and Epo1CCG for the first time in medium/large buds (Fig. 9C, Table S1). The interaction lasts for approximately 27 min into bud growth. No interaction could be recorded for Epo1 during most of its time at the bud neck (Fig. 9C). A small increase in the rate of Epo1CCG conversion occurs between min 135 and 144. We conclude that Boi1 changes its interaction from Epo1 to Nba1 during the cell cycle and thus increases the molecular heterogeneity and possibly the functions of the Bem1 interaction states.

## Discussion

Bem1 is a central scaffold protein for the Cdc42 pathway that is essential in some but not all *S. cerevisiae* strains (Dowell et al., 2010). Eliminating the GAP activity of Bem3 and thus increasing the concentration of active Cdc42 at the cortex especially during bud formation rescues the otherwise lethal deletion of *BEM1* in the strain JD47, whereas the overexpression of a Cdc42_GTP_-and membrane-binding fragment of Gic2 counteracts the positive effect of the *BEM3* deletion (Knaus et al., 2007). Whether yeast cells of a certain strain can live without Bem1 thus seems to depend on the remaining concentration of active Cdc42 at the cortex. We assumed that understanding how Bem1 helps to cope with limiting concentrations of Cdc42_GTP_ will ultimately reveal the mechanisms of its actions. A central fragment of Bem1 harboring the SH3_b_ domain with its neighboring Cdc42_GTP_ binding element, and a C-terminal fragment, containing the PB and PX domain, independently rescue a *Δbem1* strain. The C-terminal fragment binds strongly to Cdc24 but does not connect Cdc24 to Cdc42 effector proteins or to the cortex. We propose, in line with published data, that the C-terminal fragment increases the concentration of Cdc42_GTP_ by stimulating the activity of Cdc24 (Rapali et al., 2017; Smith et al., 2013; Shimada et al., 2004). The central SH3_b_CI fragment interacts with active Cdc42 and four Cdc42 effectors and needs both of its binding sites to rescue *Δbem1*-cells. Our genetic analysis further shows that SH3_b_ and CI have to operate in cis to be functional. This complements the previous finding that also SH3b and PB_Bem1_ operate in cis (Kozubowski et al., 2008). Both observations suggest a chain of binding sites that funnel active Cdc42 from its source to its targets.

We could further show that the CRIB domains of both PAKs are the essential acceptors of Cdc42_GTP._ How can the isolated SH3bCI without connection to Cdc24 still stimulate their activities? A comparison between the binding characteristics of the non-essential Boi1/2- PxxP sites and the essential Ste20/Cla4-PxxP sites suggests a molecular mechanism. Bem1 and all its SH3b-ligands are concentrated at the cell tip during bud formation and growth. The cortex-localizations of the PAKs but not of Boi1/2 strictly depend on Cdc42_GTP_ (Leberer et al., 1997; Peter et al., 1996; Wild et al., 2004; Kustermann et al., 2017). The isolated binding motives of Cla4, and of Boi1/2 bind with similar affinities to SH3_b_ whereas the SH3_b_ binding motif of Ste20 displays a significantly higher *in vitro* affinity (Gorelik and Davidson, 2012). SPLIFF analysis shows that both PAKs and Boi1/2 interact with Bem1 during bud formation and growth. Still and in contrast to Boi1/2 both PAKs do not measurably contribute to the cortical localization of SH3_b_CI or full length Bem1. We thus conclude that Bem1 interacts much stronger with the inactive PAKs than with their cortex-localized Cdc42_GTP_-bound forms. To explain the stimulatory activity of SH3_b_CI we postulate that SH3_b_CI might open the CRIB domains of the PAKs to actively load them with the CI-attached Cdc42_GTP_ (Lamson et al., 2002). The Cdc42_GTP_-bound CRIB domain might then release the auto-inhibition of the kinases and at the same time disrupt the interaction with SH3_b_. The activated PAK consequently dissociates from Bem1. This sequence describes Bem1 not as a passive scaffold but more similar to the kinase scaffold Ste5 as a co-activator that regulates through binding the activity of the PAKs and stimulates the synthesis and the transfer of Cdc42 (Bhattacharyya et al., 2006). Support for our model comes from the observation that Bem1 needs the fully functional SH3bCI to activate Ste20 and Cla4 also during osmo-stress (Tanaka et al., 2014; Chang et al., 1999). A similar activity was proposed for Scd2, the Bem1 homologue from *S.pombe*, that binds with its second SH3 domain the Ste20 homologues Shk1, and increases the auto-phosphorylation activity of the kinase (Tanaka et al., 2014; Chang et al., 1999).

The Bem1-Cdc24 complex is central source and distributor of Cdc42_GTP_. A temporal map of the interaction network of this complex might reveal where at a given time the activated Cdc42 is preferentially directed. Figure 10 summarizes the SPLIFF-derived cellular flow of Cdc42_GTP_ through the cell cycle. During bud site formation and bud growth Cdc42 is channeled directly to Exo70 and possibly from Boi1/2 to the other Cdc42-activated exocyst component Sec3. Bud6 binds and stimulates the Cdc42-activated formin Bni1. The temporal formation of the Bem1-Boi1/2-Bud6 complex might thus boost the Bni1-catalyzed actin filament formation in small buds. As the binding sites of Bem1 for Boi1/2 and Exo70 do not overlap, a super-complex that stimulates and coordinates actin assembly and vesicle fusion during bud assembly and early growth seems plausible (Kustermann et al., 2017; Liu and Novick, 2014). This complex disassembles in large buds and does not detectably form during cytokinesis (Figs. 8, 10).

**Figure 10:**
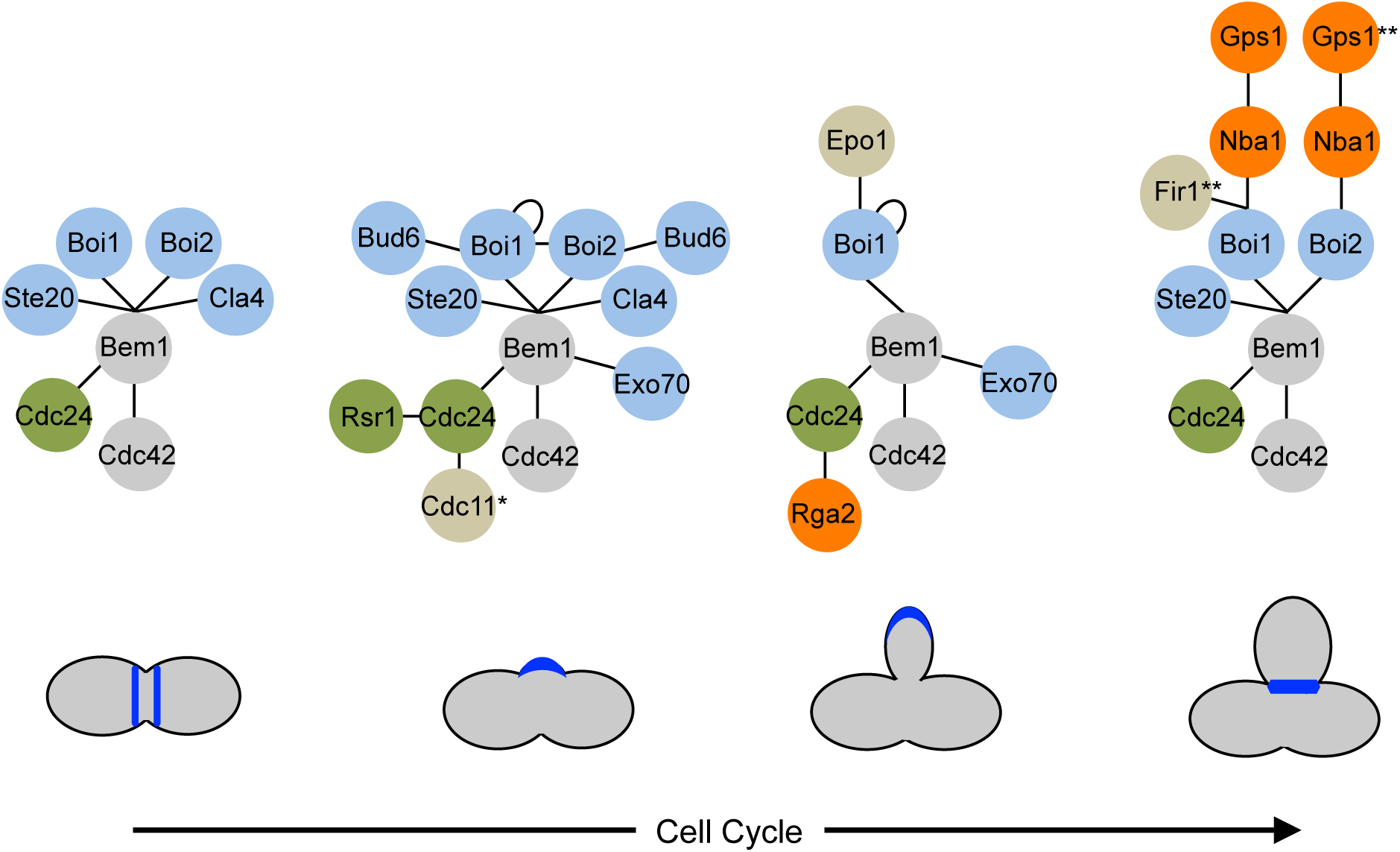
Cartoon of the time resolved interaction network of Bem1. Edges indicate interactions between the connected proteins. Edges that convert on a common point indicate competition between the different ligands. Green color indicates proteins that promote the synthesis of active Cdc42. Orange color indicate proteins that reduce the concentration of active Cdc42. Blue color indicate effectors of active Cdc42, or proteins that bind to effectors (Bud6-Bni1). Epo1 binds to the same site of Boi1 as Nba1 or Fir1 but at a different cell cycle phase.* The interaction between Cdc24 and Cdc11 occurs only during bud site assembly and was determined by SPLIFF analysis in a previous study (Chollet et al. 2019). **The time point of Fir1- and Gps1 binding to Bem1 was not derived by SPLIFF analysis but indirectly through their effects on Bem1 localization (Fig. 7).

The PAKs Ste20 and Cla4 contact Bem1 at the same site as Boi1/2 and form alternative, exclusive interaction states. The Cla4-Bem1-Cdc42-Cdc24 and Ste20-Bem1-Cdc42-Cdc24 interaction states coexist with the Boi1/2-Bem1-Cdc42-Cdc24 interaction states except during cytokinesis where only Ste20-Bem1-Cdc42-Cdc24 and Boi1/Boi2-Bem1-Cdc42-Cdc24 are detectable (Figs. 8, 10). The persistent activation of Ste20 but not of Cla4 during cell separation correlates with its role in cytokinesis and its early, CDK-independent activation in the next cell cycle (Moran et al., 2019). The Ste20-Bem1-Cdc42-Cdc24 interaction state plays also a role during mating, whereas the Cla4-Bem1-Cdc42-Cdc24 interaction state is specific for forming the septin base during bud assembly (Winters and Pryciak, 2005). Both PAKs perform many other functions in addition (Tanaka et al., 2014; Hofken and Schiebel, 2002; Drogen et al., 2000). The Cdc42 sensitivity of cells lacking the Bem1 binding sites in *STE20* and *CLA4,* suggest that the Bem1-Cdc42-Cdc24 interaction states are the operative units for many if not all PAK-activities (Tanaka et al., 2014).

Nba1 binds to Boi1/2-Bem1-Cdc24 exclusively during cytokinesis at the neck of the cell. An alternative docking site for Bem1 at the neck was discovered by the Boi1-Fir1 complex. Cdc24 is unequally distributed between Nba1 and Fir1, with more Cdc24 associated with Fir1 (Fig. 7F). Fir1 is known to delay SS formation by inhibiting the cell separation kinase Cbk1 (Brace et al., 2019). This activity might be supported by the local production of Cdc42_GTP_ and the subsequent activation of Ste20. In contrast, less Cdc24 is bound to Nba1 and thus less active Cdc42 is generated. In addition, the loss of *NBA1* is synthetic lethal to a certain allele of the essential cytokinesis factor IQGAP (in yeast: *IQG1*), suggesting that Nba1 might stimulate abscission (Tian et al., 2014). We therefore propose that the Gps1-Nba1-Boi1/2-Bem1-Cdc24 complex is an abscission promoting complex, whereas the Fir1-Boi1-Bem1-Cdc24 complex might act as an abscission inhibiting complex (Brace et al., 2019).

Our SPLIFF analysis of Bem1 and Boi1 shows that both change the binding partners of their SH3 domains during the cell cycle (Figs. 8, 9, 10). This finding is surprising as the SH3 domains bind to the PxxP motives of their interaction partners with similarly low affinities. By comparing the FRAP of full-length Boi1 at the bud neck with the FRAP of the isolated SH3_Boi1_, we could however show that the SH3_Boi1_-Nba1/Fir1 interactions are cooperatively embedded into a larger binding pocket formed by the PxxP motif and additional contact sites that are still to be defined (Figs. 6, 7). The assumption that this larger pocket is induced upon contact with the SH3 domain and only formed once Nba1/Fir1 assemble at the bud neck could explain why Boi1 interacts during abscission predominantly with Nba1 and Fir1 but not any longer with Epo1. The time dependent selectivity of SH3-PxxP interactions and the herein suggested molecular mechanism might be a general phenomenon whose detection would require the application of time-sensitive techniques as used in this study.

## Material and Methods

### Growth conditions and cultivation of yeast strains

All yeast strains were derivatives of JD47, a descendant from a cross of the strains YPH500 and BBY45 (Dohmen et al., 1995). Cultivation of yeast was performed in standard SD or YPD media at 30°C or the indicated temperatures as described (Guthrie and Fink, 1991). Media for Split-Ubiquitin interaction assay and selection for the loss of centromeric *URA3*-containing plasmids comprised 1 mg/ml 5-fluoro-orotic acid (5-FOA, Formedium, Hunstanton, UK).

### Construction of plasmids, gene fusions and manipulations

Construction of N_ub_ and C_ub_ gene fusions as well as GFP-, mCherry- or mCherry-C_ub_-RGFP (CCG)-fusions was performed as described (Wittke et al., 1999; Dünkler et al., 2012; Neller et al., 2015; Moreno et al., 2013). Bem1CRU/ -GFP-CCG were constructed by genomic in-frame insertions of the *GFP-*, *CRU-*, or *CCG*-modules behind the coding sequences of *BEM1* or its alleles. In brief, a PCR-fragment of the C-terminal region of the respective target gene lacking the stop codon was cloned via *Eag*I and *Sal*I restriction sites in front of the *CRU*-, *GFP*-, *mCherry*-, or CCG-module on a pRS303, pRS304 or pRS306 vector (Sikorski and Hieter, 1989). Plasmids were linearized using unique restriction sites within this sequence and transformed into yeast cells for integration into the genomic target ORF. Colony PCR with diagnostic primer combinations was used to verify the successful genomic integration. Centromeric plasmids expressing different fragments of *BEM1* were obtained by ligation of PCR fragments spanning the respective region of *BEM1* behind the sequence of the *P_MET17_* - GFP module on the pRS313 vector (Table S2) (Sikorski and Hieter, 1989). Mutations in the coding region of *BEM1*, *STE20, CLA4,* or *BOI1* were obtained by overlap-extension PCR using plasmids containing the corresponding ORFs as templates.

Insertion of mutations into the *BEM1*, *STE20, BOI1,* or *CLA4* loci were performed in yeast strains lacking the ORFs of the respective gens but still containing their 5’ and 3’ UTR sequences. Mutations were introduced in the respective genes on an integrative pRS vector containing the up- and down stream sequences of the gene (Sikorski and Hieter, 1989). Yeast strains lacking the corresponding ORF were then transformed with the mutated gene on the integrative vector linearized in the promoter sequence of the gene. Successful integration was verified by diagnostic PCR.

Alternatively, insertion of genomic mutations were achieved by CRISPR/Cas9 (Laughery et al., 2015). To introduce the mutations at the chosen sites guideRNA sequences were cloned into pML plasmids and co-transformed with oligonucleotides harboring the desired mutations. Successful manipulations were verified by PCR product sequencing of the respective genomic ORFs. Details of the introduced mutations are listed in Table S3.

In certain strains the native promoter sequence was replaced by *P_MET17_* through recombination with a PCR fragment generated from pYM-N35 and primers containing sequences identical to the respective genomic locations at their 5’ ends (Janke et al., 2004). GST fusions were obtained by placing the ORF of the respective gene or gene fragment in frame behind the *E. coli GST* sequence on the pGEX-2T plasmid (GE Healthcare, Freiburg, Germany) using BamHI and EcoRI restriction sites. Fusions to the human O6-Alkyl-DNA transferase (SNAP-tag, New England Biolabs, Beverly, MA) were expressed from plasmid pAGT-Xpress, a pET-15b derivative (Schneider et al., 2013). Gene fragments were inserted in frame into a multi-cloning site located between the upstream *6xHIS*-tag coding sequence and the downstream *SNAP*-tag-coding sequence. The *6xHIS*-tag fusions were obtained by placing the ORF of the respective gene or gene fragment behind the *E. coli 6xHIS*-tag sequence on the previously constructed pAC plasmid (Schneider et al., 2013).

Gene deletions were performed by one step PCR-based homologous recombination using pFA6a natNT2, pFA6a hphNT1, pFA6a kanMX6, pFA6a CmLEU2 and pFA6a HISMX6 as templates (Bähler et al., 1998; Longtine et al., 1998; Janke et al., 2004; Schaub et al., 2006). Lists of plasmids and yeast strains used in this study can be found in Tables S2 and S3 of the supplementary information. Plasmid maps can be obtained upon request.

### Split-Ub interaction analysis

Split-Ubiquitin array analysis: A library of 548 different α-strains each expressing a different N_ub_ fusion were mated with a *Bem1*-*C_ub_-R-URA3* (CRU), *Bem1_WK_*-*C_ub_-R-URA3* (CRU), or *Bem1_ΔPB_*-*C_ub_-R-URA3* (CRU) expressing a-strain. Diploids were transferred as independent quadruplets on SD media containing 1 mg/ml 5-FOA and different concentrations of copper to adjust the expression of the N_ub_ fusions (Dünkler et al., 2012). Individual Split-Ub interaction analysis: CRU and N_ub_ expressing strains were mated or co-expressed in haploid cells and spotted onto medium containing 1 mg/ml FOA and different concentrations of copper in four 10-fold serial dilutions starting from OD_600_=1. Growth at 30°C was recorded every day for 3 to 5 days.

### Mating efficiency

Saturated cultures of JD47 cells containing the respective allele of *STE20* and expressing Gic2_PBD_ from a centromeric plasmid under the control of a *P_MET17_* promoter and JD53 cells carrying a Kanamycin-tagged *PTC1* gene and expressing Gic2_PBD_ were resuspended in media containing no or 70 µM methionine and grown for 6 h at 30 °C. Cells were adjusted to an OD_600_ of 1 and equal amounts of JD53 and JD47 cells mixed and incubated for 4 h at 30 °C. Cells were diluted 1/20 and 250 µL of each mating were spread on media selecting for diploids. Colony numbers were counted after 2 d at 30°C.

### Preparation of yeast cell extracts

Exponentially grown yeast cell cultures were pelleted and resuspended in yeast extraction buffer (50 mM HEPES, 150 mM NaCl, 1 mM EDTA) with 1x protease inhibitor cocktail (Roche Diagnostics, Penzberg, Germany). Cells were lysed by vortexing them together with glass beads (3-fold amount of glass beads and extraction buffer to pellet weight) 12 times for one minute interrupted by short incubations on ice. The obtained yeast cell extracts were clarified by centrifugation at 16,000 g for 20 min at 4°C.

### Recombinant protein expression and purification from *E. coli*

All proteins were expressed in *E.coli* BL21DE3 cells. GST-Bud6_1-320_ was expressed at 30 °C for 5 h in LB medium after induction with 1 mM IPTG. GST fusions to SH3 domains of Boi1 and Boi2 and 6xHis-Nba1_202-289_-SNAP were expressed at 18 °C in SB medium for 20 h after induction with 0,1 mM IPTG. Cells were pelleted, washed once with PBS and stored at −80 °C until lysis. All subsequent purifications were carried out on an Äkta Purifier chromatography device (GE Healthcare, Freiburg, Germany). Cells expressing GST fusion proteins were resuspended in PBS containing protease inhibitor cocktail (Roche Diagnostics, Penzberg, Germany) and lysed by lysozyme treatment (1 mg/ml, 30 min on ice), followed by sonication with a Bandelin Sonapuls HD 2070 (Reichmann Industrieservice, Hagen, Germany). Extracts were clarified by centrifugation at 40,000 g for 10 min at 4 °C and the proteins were purified using a 5 ml GSTrap column (GE Healthcare) and subsequent size exclusion chromatography on a Superdex 200 16/60 column vs. HBSEP buffer (10 mM HEPES, 150 mM NaCl, 3 mM EDTA, 0.05% Tween 20, pH 7.4). Purified protein was concentrated and stored on ice. 6xHis-Nba1_202-289_-SNAP was lysed in IMAC buffer (50 mM KH_2_PO_4_, 300 mM NaCl, 20 mM Imidazole containing protease inhibitor cocktail) as described above and purified by IMAC and elution in a linear imidazole gradient on a 5 ml HisTrap HP column (GE healthcare, Freiburg, Germany) followed by size exclusion chromatography as described above. 6xHis-Exo84_121-141_-AGT was purified by IMAC as descried above followed by anion exchange chromatography on a ResourceQ column (GE healthcare, Freiburg, Germany) in TRIS buffer (50 mM TRIS, 50 mM NaCl, pH 7,5). 6xHis-Sec3_481-521_-AGT was directly used after IMAC and buffer exchange to PBS via a PD10 desalting column (GE Healthcare, Freiburg, Germany).

### GST-pulldown assay

GST-tagged proteins were immobilized on glutathione sepharose beads (GE Healthcare, Freiburg, Germany) directly from *E. coli* extracts. After 1 h incubation at 4 °C with either yeast extracts or purified proteins under rotation at 4 °C, the beads were washed 3 times with the respective buffer. Bound material was eluted with GST elution buffer (50mM TRIS, 20mM reduced glutathione) and analyzed by SDS-PAGE followed by Coomassie staining and immunoblotting with anti-His (dilution: 1:5000; Sigma-Aldrich, Steinheim, Germany), or anti-GFP-antibodies (dilution 1:1000; Roche Diagnostics, Penzberg, Germany).

### SPR measurements

Binding affinities were measured using purified and immobilized GST-SH3_Boi1_ or GST-SH3_Boi2_ as ligands on an anti-GST chip on a Biacore X100 device (GE Healthcare, Freiburg, Germany). HBSEP buffer was used as running buffer in all experiments. The chip was prepared by covalent coupling of an anti-GST antibody (GE Healthcare) as capture molecule to the dextran surface of both flow cells of a CM5 Chip (GE Heathcare) using an amine coupling kit (GE healthcare). GST tagged ligand proteins were captured on the detection flow cell of the chip and free GST was captured on the reference flow cell. Purified 6xHis-NBA1_202-289_-SNAP as analyte molecule was prepared in suitable concentrations in running buffer and kinetics were measured with constant flow over the previously prepared chip. Regeneration after each cycle was achieved by a 20 s injection pulse with 10 mM Glycine pH 2.0. The equilibrium binding constant K_D_ was subsequently determined by the X100 evaluation software using background substracted sensograms. All measurements were performed at least as triplicate.

### Fluorescence microscopy

For microscopic inspection yeast cells were grown overnight in SD media, diluted 1:8 in 3-4 ml fresh SD medium, and grown for 3 to 6 h at 30 °C to mid-log phase. About 1 ml culture was spun down, and the cell pellet resuspended in 20-50 µl residual medium. 3 µl were spotted onto a microscope slide, the cells were immobilized with a coverslip and inspected under the microscope. For time-resolved imaging 3 µl of prepared cell suspension was mounted on a SD-agarose pad (1.7% agarose), embedded in a customized glass slide, and sealed by a cover slip fixed by parafilm stripes. Imaging was started after 15 to 30 min recovery at 30 °C. SPLIFF and other time-lapse experiments were observed with a wide-field fluorescence microscope system (DeltaVision, GE Healthcare) provided with a Olympus IX71 microscope, a steady-state heating chamber, a CoolSNAP HQ2 and CascadeII512-CCD camera both by Photometrics, a U Plan S Apochromat 100Å∼ 1.4 NA oil **∞**/0.17/FN26.5 objective and a Photofluor LM-75 halogen lamp (Burlington, VT, USA). Images were visualized using softWoRx software (GE Healthcare) and adapted z series at 30°C. Exposure time was adapted to the intensity of GFP and mCherry signal for every fluorescently labeled protein to reduce bleaching and phototoxicity. For further analyses an Axio Observer spinning-disc confocal microscope (Zeiss, Göttingen, Germany), equipped with an Evolve512 EMCCD camera (Photometrics, Tucson, USA), a Plan-Apochromat 63Å∼/1.4 oil DIC objective, and 488 nm and 561 nm diode lasers (Zeiss, Göttingen, Germany) was used. Images were analyzed with the ZEN2 software (Zeiss).

### Quantitative analysis of microscopy data and SPLIFF measurements

Microscopy data were processed and analyzed using ImageJ64 1.49 software. For standard fluorescence signal quantification three regions of interest (ROIs) were determined, first the signal of interest (e.g. tip, bud neck), second a region in the cytosol and third a randomly chosen position outside of the cell (background). The mean grey values of each ROI (*I_fluorescence_, I_cytosol_, I_background_*) were quantified after z-projection. To compare the fluorescence signals of a protein in certain strains the relative fluorescence (*I_relative_*) signal of the protein was calculated after subtraction of the background.

*I_relative_ = I_fluorescence_-I_background_ / I_cytosol_-I_background_*

SPLIFF analysis for temporal and spatial characteristics of Bem1-, Boi1- and Epo1-CCG interactions was performed as described (Moreno et al., 2013; Dunkler et al., 2015). *P_MET17_* promoter controlled CCG fusions were expressed in SD medium without methionine and mixed with *MAT alpha* cells expressing suitable *P_CUP1_-N_ub_-HA* fusions. Both cell types were immobilized on a SD agarose pad and mating induced interaction monitored by three channel z-stack (5×0.6 µm) microscopy every 2, 3 or 5 min. z-slices with fluorescence signals were projected by SUM-projection. The fluorescence intensities (FI) of mCherry and GFP channels were determined by integrated density measurements of the region of interest and a region within the cytosol. For each time point and channel the intracellular background was subtracted from the localized signal to obtain the localized fluorescence intensity (FI_red_ and FI_green_). The values were normalized to the time point before cell fusion. The resulting relative fluorescence intensity RFI(t) was then used to calculate the conversion FD(t) :

*FD(t)=RFI_red_-RFI_green_/RFI_red_*

FD(t) as a readout of CCG-to CC conversion describes its temporal progress in percent. Excel was utilized for initial calculations, Prism 7.0 software (GraphPad) to plot the final graph. All error bars indicate SEM.

### FRAP experiments

FRAP experiments were performed as described elsewhere with an iMIC digital microscope with a 60X objective (Till Photonics, Munich, Germany) at RT (Phair et al., 2004). Pictures were acquired with a series of 5 z-slices each separated by 0.5 µM. Four images were taken before the ROI was bleached with 100% laser power, a dwell time of 1,2 s/µm^2^, a line overlap of 42% and an experimental loop count of 10 to 20.

Pictures were taken at a constant time interval of 0.9 s after bleaching except SH3_Boi1_-GFP signal recovery (each 0.25 s in a single z-layer). Initial z-slice projection and fluorescence quantification was performed with the software iMIC Offline analysis. Alternatively, an Axio Observer spinning-disc confocal microscope (Zeiss, Göttingen, Germany), equipped with a Zeiss Plan-Apochromat 63Å∼/1.4 oil DIC objective, a 488 nm diode lasers and an UGA-42 photo-manipulation system was used (Rapp OptoElectronic, Wedel, Germany). Initial signal measurements were ImageJ based. Subsequently all data sets were double normalized using Excel. The software Prism 6.0 (GraphPad) was used for the fitting of the double normalized data to a one-phase association curve.

### Statistical evaluation

GraphPad Prism was applied for statistical data evaluation. The distributions of the data sets were analyzed by the D’Agostino and Pearson normality test. Students *t*-tests were used to analyze data following a normal distribution whereas Mann-Whitney-U-tests were used for data that did not pass this criteria. The one-way ANOVA or the Kruskal-Wallis ANOVA tests were used to compare data sets from more than two groups.

## Acknowledgements

We thank Steffi Timmermanns, and Nicole Schmidt for technical assistance. The work was funded by grants from the DFG to N.J. (Jo 187/5-2; Jo 187/9-1). The authors declare no conflicts of interest.

## Author contribution

S. Grinhagnes, A. Dünkler, L. Rieger, Y. Wu, and N. Johnsson designed and analyzed the experiments. S. Grinhagnes, and A. Dünkler performed the microscopic analysis and the photokinetic experiments. A. Dünkler, S. Grinhagens performed the SPLIFF analysis. Y. Wu, and L. Rieger performed the protein interaction screens. L. Rieger, P. Brenner, Y. Wu, and T. Gronemeyer performed the biochemical characterizations of wild type and mutated PxxP-SH3 interactions. N. Johnsson, A. Dünkler, S. Grinhagens, J. Rieger and Y. Wu designed the study. N. Johnsson together with the help of A. Dünkler wrote the manuscript.

