## Supplementary Figures and Tables for "A time resolved interaction analysis of Bem1 reconstructs the flow of Cdc42 during polar growth"

### Supplementary Information

### Supplementary Figures and Tables

#### Figure S1

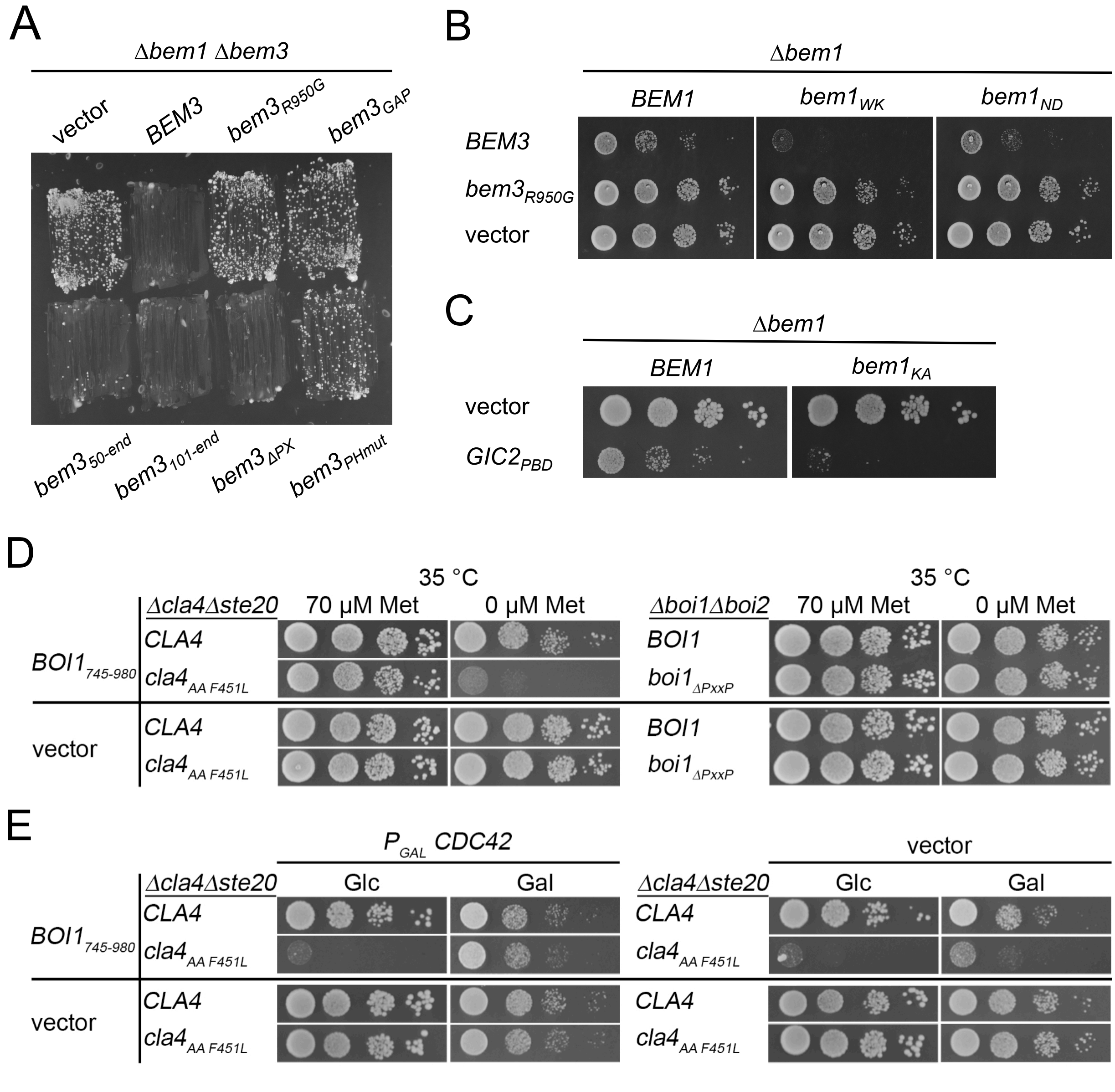

(A) Δ*bem1*Δ*bem3* cells harboring *BEM1* on a centromeric, *URA3* expressing plasmid and containing the indicated GFP fusion alleles of *BEM3* on an additional plasmid were streaked on medium containing 5-FOA and selecting for the presence of the *bem3* containing plasmids. Growth of the cells indicates alleles that do not complement the function of *BEM3*. $bem3_{PHmut}$ harbors the residue exchanges R644S, R645S, K647D that are known to impair the binding of the PH domain to phospholipids. $bem3_{R950G}$ contains an exchange in the catalytic site of the GAP domain that eliminates the activity of Bem3.
(B) Cells containing the indicated *bem1* alleles and an empty vector or expressing Bem3-GFP, or $Bem3_{R950G}$-GFP from a centromeric plasmid under the control of the $P_{MET17}$ promoter were spotted in 10-fold serial dilutions on media without methionine to induce the expression of the *bem3* alleles. Cells were incubated for two days at 37$^o$C.
(C) Δ*bem1* cells expressing *BEM1* or $bem1_{KA}$ from a centromeric plasmid under control of the $P_{MET17}$ promoter, and containing an empty vector (upper lanes), or $Gic2_{PBD}$ expressed from the $P_{MET17}$-promoter (lower lanes), were spotted in 10-fold serial dilutions on medium lacking methionine, and incubated for

two days at 30$^{o}$C. *bem1$_{KA}$* harbors the residue exchanges K482A that is known to impair the binding of Cdc24

(D) Left panel: $\Delta$*ste20* cells expressing *CLA4* or *cla4$_{AAF451L}$* and either the PH domain of Boi1 (Boi1$_{745-980}$), or an empty vector were spotted on plates containing 70µM methionine, or no methionine to fully express Boi1$_{745-980}$. Right panel: $\Delta$*boi2* cells expressing *Boi1* or *boi1$_{\Delta PxxP}$* and either the PH domain of Boi1 (Boi1$_{745-980}$), or an empty vector were spotted on plates containing 70µM methionine, or no methionine to fully express Boi1$_{745-980}$. Cells were incubated for two days at 35$^{o}$C. (E) Cells as in left panel of (D) but additionally expressing Cdc42 on a centromeric vector under the control of the P$_{GAL1}$ promoter (left panel), or containing an empty vector (right panel). Cells were spotted on media containing glucose (Glc) and lacking methionine to overexpress Boi1$_{745-980}$ and repress Cdc42, or on media lacking methionine and containing galactose (Gal) to simultaneously overexpress Boi1$_{745-980}$ and Cdc42. Note the partial rescue of *cla4$_{AAF451L}$*-cells overexpressing Cdc42.

### Figure S2

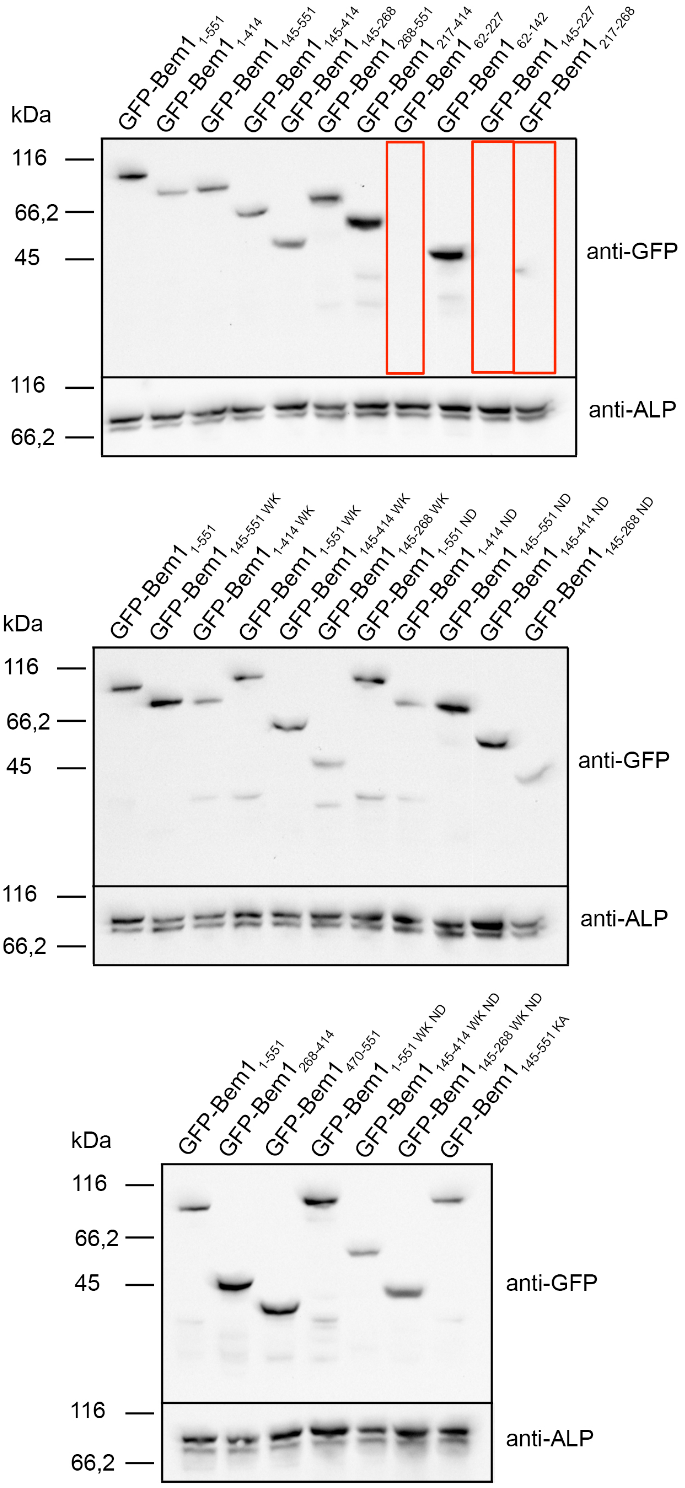

Expression levels of different *BEM1* alleles. N-terminal GFP-tagged fragments of Bem1 and mutants thereof were expressed under control of the $P_{MET17}$ promoter from a centromeric plasmid. Yeast cells were grown in media containing 70µM methionine to an $OD_{600}$ of 1.5 -2. Extracts containing equal amounts of proteins were separated by SDS-PAGE and stained with anti-GFP antibody after transfer onto nitrocellulose. Equal loading was controlled by staining the blot with anti-alkaline phosphatase antibody (anti-ALP). Lanes showing fragments that are not measurably expressed were framed by a red box.

### Figure S3

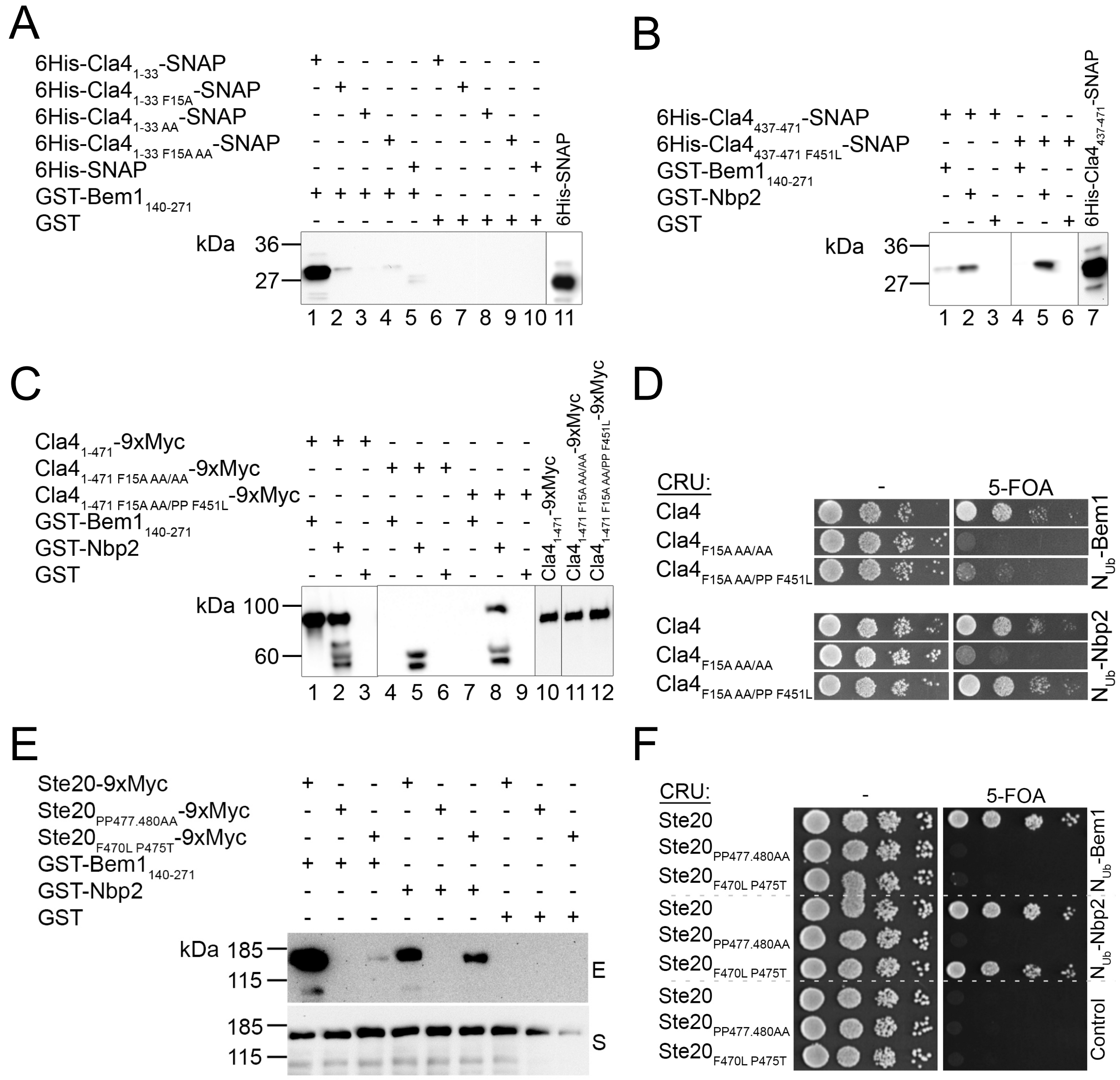

Characterization of mutations in Cla4 and Ste20 that impair the interactions of both proteins with Bem1 but not with Nbp2. (A) Extracts of *E.coli* cells expressing a 6His-SNAP fusion protein of binding site 1 of Cla4 (6His-$Cla4_{1-33}$-SNAP) containing no mutation (lanes 1, 6), a phenylalanine to leucine exchange at position 15 (6His-$Cla4_{1-33F15A}$-SNAP; lanes 2, 7), an alanine to proline exchange at positions 22 and 25 (6His-$Cla4_{1-33AA}$-SNAP; lanes 3, 8), the triple mutations (6His-$Cla4_{1-33F15AAA}$-SNAP; lanes 4, 9), or a 6His-SNAP fusion (lanes 5, 10) were incubated with Glutathione-coated beads exposing GST-$Bem1_{140-271}$ (lanes 1-5), or GST (lanes 6-19). Shown is the anti-His western blot of the glutathione eluates of the beads after SDS-PAGE and transfer onto nitrocellulose. Lane 11 documents the input of the 6His-SNAP fusion. (B) Analysis as in (A) but with extracts of *E.coli* cells expressing a 6His-SNAP fusion protein of binding site 2 of Cla4 (6His-$Cla4_{437-471}$-SNAP) containing no mutation (lanes 1-3), or a phenylalanine to leucine exchange at position 451 (6His-$Cla4_{437-471F451L}$-SNAP; lanes 4-6). Extracts were incubated with beads exposing GST- $Bem1_{140-271}$ (lanes 1, 4), GST-Nbp2 (lanes 2, 5), or GST (lanes 3, 6). The input is shown in lane 7. (C) A MYC-tagged fragment of Cla4 containing both binding sites and no mutations ($Cla4_{1-471}$9xMyc; lanes 1-3), the indicated mutations $Cla4_{1-471F15AAA/AA}$ (lanes 4-6), or $Cla4_{1-471F15AAA/PPF451L}$ (lanes 7-9) were expressed in yeast from a centromeric vector under control of the $P_{MET17}$ promoter. Extracts of these cells were incubated with bead-coupled GST-$Bem1_{140-271}$ (lanes 1, 4, 7), GST-Nbp2 (lanes 2, 5, 8) or GST (lanes 3, 6, 9). The inputs of the Myc-tagged fusions are shown in lanes 10-12. The eluted proteins were separated by SDS-PAGE and stained with anti-Myc antibody after transfer onto nitrocellulose. The protein doublet below 60 kDa is occasionally observed in lanes displaying GST-Nbp2 and might arise by a cross-reactivity of the antibodies. (D) Split-Ub assay of yeast cells co-expressing $N_{ub}$-Bem1 (upper panel) or $N_{ub}$-Nbp2 (lower panel) together with CRU fusions to Cla4 or its indicated mutants, expressed from a centromeric vector under control of the $P_{MET17}$ promoter. 4µl of cells of $OD_{600}$ of 1 were spotted in 10 fold serial dilutions on media lacking (left panels) or containing 5-FOA. Growth on media containing 5-FOA indicates interaction between the respective $N_{ub}$ and $C_{ub}$ fusion. (E) As in (C) but with cells co- expressing 9xMyc fusions to Ste20, to Ste20 containing alanine exchanges for proline at positions 477 and 489 ($Ste20_{PP477,480AA}$), or to Ste20 containing an phenylalanine to leucine exchange at position 470 and a proline to threonine exchange at position 475 ($Ste20_{F470L\ P475T}$). (F) Split-Ub assay as in (D) but with cells co-expressing the indicated $N_{ub}$ fusions with CRU fusions to Ste20, or to the indicated mutants of Ste20. The CRU fusions were expressed from a centromeric vector under control of the $P_{MET17}$ promoter.

**Figure S4**

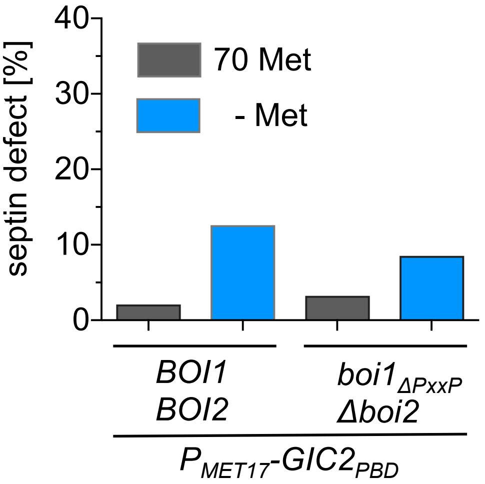

The disruption of the Boi1/2-Bem1 connection does not induce defects in septin organization. Quantification of wild type- and Δ*boi2 boi1*$_{\Delta PxxP}$-cells as in Fig. 5B. 500-600 cells containing a GFP fusion to the septin *SHS1* were observed with the fluorescence microscope under conditions of low (70 µM methionine), or high (no methionine) expression of Gic2$_{PBD}$. No increased septin mis-localization is detectable in Δ*boi2 boi1*$_{\Delta PxxP}$-cells.

**Figure S5**

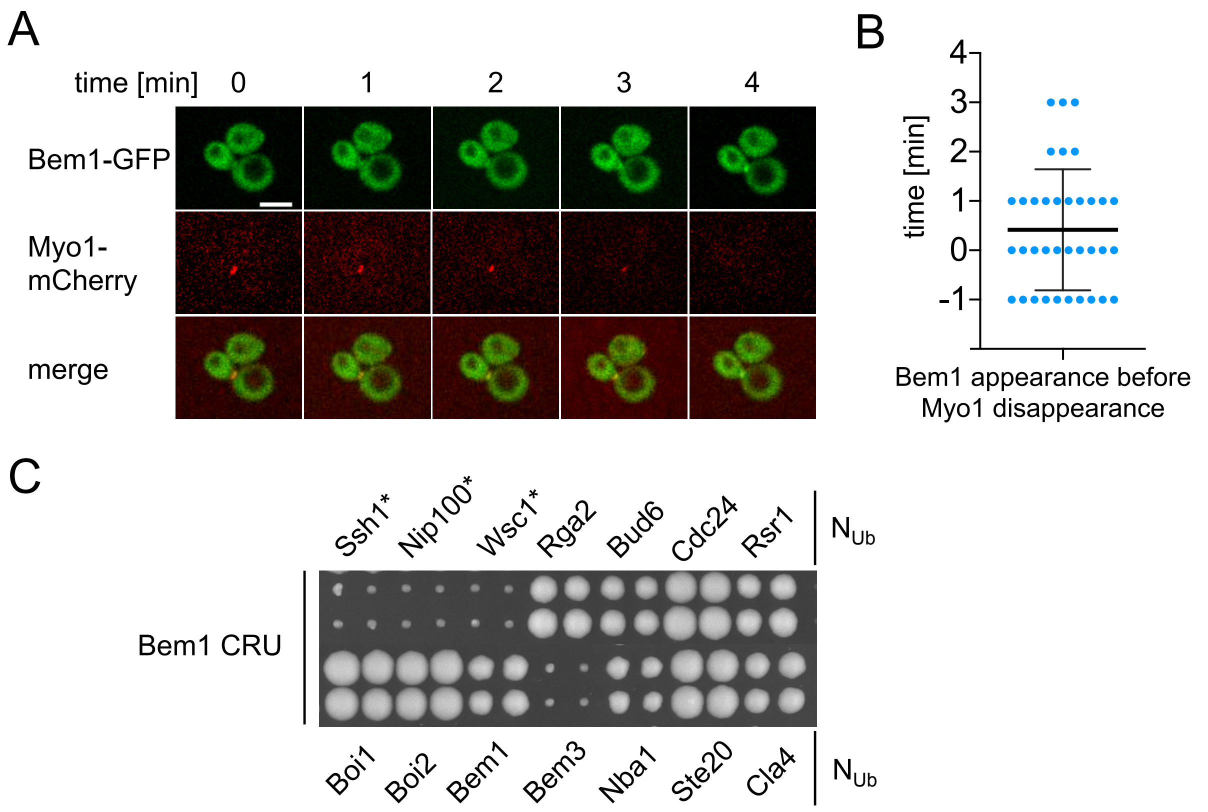

(A) Time-lapse analysis of cells co-expressing Bem1-GFP and Myo1-mCherry. Shown are the frames where Myo1-mCherry (middle panel) contracts at the bud neck during cytokinesis. Bem1-GFP (upper panel) becomes visible at the bud neck shortly after Myo1-mCherry disappears. The merge of both channels is shown in the lower panel. Scale bar indicates 3μM. (B) Quantification of 36 cells. Plotted are the times of Bem1-GFP appearances in relation to the disappearance of Myo1-mCherry. Bem1 appears in average 0.4 min before Myo1 is removed from the site of cell separation.
(C) Split-Ub analysis as in Figure 1 but of cells expressing Bem1CRU together wit the indicated $N_{ub}$ fusions expressed from their native promoters. * indicates $N_{ub}$ fusions expressed from the $P_{CUP1}$ promoter. Interaction was scored after four days of growth on media containing 5-FOA.

### Supplementary Tables

### Table S1

Calculated conversion rates (FD) of CCG- to CC conversion in percent. Listed are FD values of the single cell measurements during mating, bud growth and cytokinesis for SPLIFF substrates Bem1-, Boi1- and Epo1-CCG.

#### Bem1-CCG x $N_{ub}$-Boi1

| cell fusion | cell 1 | cell 2 | cell 3 | cell 4 | cell 5 | cell 6 | cell 7 | cell 8 | cell 9 | cell10 |
|---|---|---|---|---|---|---|---|---|---|---|
| 0 min | 0,0 | 0,0 | 0,0 | 0,0 | | 0,0 | 0,0 | 0,0 | 0,0 | 0,0 |
| 5 min | 17,5 | 9,2 | 0,6 | 36,7 | | 10,0 | 62,1 | 24,7 | -3,2 | 68,0 |
| 10 min | 6,5 | 17,9 | 12,1 | 40,9 | | 21,6 | 59,6 | 100,0 | 12,5 | 71,3 |
| 15 min | 27,8 | | 32,2 | 47,8 | | | 68,5 | 85,4 | 16,1 | 73,3 |
| 20 min | | | 24,7 | | | | 49,7 | | | 74,3 |
| 25 min | | | 59,7 | | | | 63,6 | | | 70,1 |
| **bud growth** | | | | | | | | | | |
| 50 min | 41,8 | 40,5 | 67,8 | 18,7 | 22,0 | 30,2 | 75,8 | 46,9 | 70,3 | 69,4 |
| 55 min | 96,1 | 29,0 | 78,9 | 27,5 | 17,6 | 28,7 | 78,1 | 88,7 | 84,3 | 69,8 |
| 60 min | 100,0 | 51,4 | 99,2 | 44,3 | 52,4 | 57,8 | 81,5 | 94,0 | 68,3 | 77,4 |
| 65 min | 97,9 | 41,2 | 100,0 | 46,1 | 58,1 | 59,4 | 80,9 | 100,0 | 66,1 | 77,8 |
| 70 min | 100,0 | 45,8 | 100,0 | 43,6 | 67,8 | 63,6 | 83,2 | 100,0 | 58,2 | 81,1 |
| 75 min | 95,9 | 39,4 | 100,0 | 47,0 | 66,1 | 59,8 | 85,3 | 100,0 | 59,4 | 79,6 |
| 80 min | 100,0 | 43,8 | 100,0 | 41,9 | 58,9 | 80,5 | 83,8 | 100,0 | 68,8 | 82,3 |
| **cytokinesis** | | | | | | | | | | |
| 0 min | 31,7 | 19,2 | | 51,5 | 66,1 | 50,4 | | | | |
| 2 min | 35,9 | 39,1 | 50,3 | 51,4 | 70,1 | 50,7 | 61,9 | | | |
| 4 min | 38,2 | 23,2 | 68,2 | 57,4 | 73,5 | 52,1 | 63,6 | | | |
| 6 min | 40,8 | 22,5 | 65,7 | 48,8 | 75,7 | 55,7 | 62,5 | | | |
| 8 min | 41,7 | 14,5 | 59,1 | 64,1 | 89,3 | 68,1 | 70,7 | | | |
| 10 min | 41,0 | 38,6 | 70,1 | | 100,0 | 62,7 | 75,0 | | | |

#### Bem1-CCG x $N_{ub}$-Boi2

| cell fusion | cell 1 | cell 2 | cell 3 | cell 4 | cell 5 | cell 6 | cell 7 | cell 8 | cell 9 | cell 10 |
|---|---|---|---|---|---|---|---|---|---|---|
| 0 min | 0,0 | 0,0 | 0,0 | 0,0 | 0,0 | 0,0 | 0,0 | 0,0 | | |
| 5 min | 17,0 | -0,1 | 20,2 | 0,5 | 24,1 | 50,0 | 11,6 | 28,5 | | |
| 10 min | 16,6 | 6,9 | 18,9 | 40,1 | 2,7 | 100,0 | 9,3 | 63,8 | | |
| 15 min | 28,1 | 19,8 | 45,1 | 50,1 | 8,4 | 93,1 | 15,7 | 18,5 | | |
| 20 min | 34,4 | | 100,0 | 65,7 | | | 22,4 | | | |
| 25 min | | | 77,3 | | | | 56,7 | | | |
| **bud growth** | | | | | | | | | | |

| | | | | | | | | | | |
|---|---|---|---|---|---|---|---|---|---|---|
| 50 min | 67,5 | 71,2 | 71,2 | -1,1 | 47,2 | 100,0 | 84,1 | 26,9 | -2,7 | |
| 55 min | 67,6 | 78,4 | 78,4 | 18,3 | 74,1 | 100,0 | 77,8 | -9,7 | 22,7 | |
| 60 min | 60,2 | 100,0 | 100,0 | 35,5 | 60,3 | 100,0 | 80,2 | 24,4 | 60,2 | |
| 65 min | 100,0 | 100,0 | 100,0 | 42,2 | 68,0 | 100,0 | 100,0 | 16,2 | 81,4 | |
| 70 min | 76,1 | 100,0 | 100,0 | 57,1 | 58,2 | 100,0 | 100,0 | 26,9 | 63,1 | |
| 75 min | 100,0 | 100,0 | 100,0 | 34,9 | 48,5 | 100,0 | 100,0 | 54,7 | 52,5 | |
| 80 min | 100,0 | 94,0 | 94,0 | 22,4 | 74,6 | 100,0 | 100,0 | 30,2 | 53,8 | |
| **cytokinesis** | | | | | | | | | | |
| 0 min | 11,9 | 36,9 | 20,5 | 58,1 | 52,1 | 53,1 | 0,0 | 4,2 | 33,4 | 20,9 |
| 2 min | 0,0 | 54,2 | 65,2 | 55,8 | 67,3 | 35,1 | 6,8 | 0,9 | 39,7 | 37,6 |
| 4 min | 8,2 | 13,7 | 61,8 | 46,6 | 68,3 | 43,0 | 46,1 | 7,6 | 41,4 | 31,9 |
| 6 min | 41,0 | 49,8 | 54,3 | 54,6 | 69,5 | 43,2 | 54,8 | 33,0 | 50,4 | 44,8 |
| 8 min | 30,6 | | 79,1 | 67,0 | | 47,1 | | | 61,0 | 49,9 |
| 10 min | 33,4 | | | 68,0 | | | | 60,1 | 64,7 | 53,6 |

### Bem1-CCG x $N_{ub}$-Cla4

| **cell fusion** | **cell 1** | **cell 2** | **cell 3** | **cell 4** | **cell 5** | **cell 6** | **cell 7** | **cell 8** | **cell 9** | **cell 10** |
|---|---|---|---|---|---|---|---|---|---|---|
| 0 min | 0,0 | 0,0 | 0,0 | 0,0 | | 0,0 | 0,0 | 0,0 | 0,0 | 0,0 |
| 5 min | -5,6 | 44,2 | -22,1 | 11,7 | | 17,0 | -2,2 | 12,7 | 15,6 | 4,4 |
| 10 min | -8,5 | 45,0 | 15,6 | 13,9 | | 22,6 | -6,2 | 22,4 | 31,7 | 5,4 |
| 15 min | 1,0 | 51,6 | -5,6 | 13,7 | | 27,4 | 4,8 | 25,1 | 48,8 | 1,9 |
| 20 min | 9,1 | 44,9 | -1,8 | 3,3 | | 46,9 | | 46,2 | 37,1 | 23,8 |
| 25 min | 0,0 | 28,1 | 32,2 | 25,8 | | 57,8 | | 29,6 | 86,4 | |
| **bud growth** | | | | | | | | | | |
| 50 min | 42,6 | 74,3 | 62,8 | 49,4 | -12,6 | 90,1 | 17,1 | 38,8 | 36,8 | 66,3 |
| 55 min | 51,9 | 73,2 | 91,6 | 35,4 | 7,6 | 100,0 | 56,5 | 60,9 | 69,4 | 64,2 |
| 60 min | 100,0 | 92,5 | 89,9 | 39,5 | 9,8 | 100,0 | 69,3 | 72,4 | 75,0 | 52,0 |
| 65 min | 98,1 | 96,8 | 99,1 | 58,0 | 29,1 | 100,0 | 76,5 | 74,7 | 81,7 | 58,5 |
| 70 min | 100,0 | 92,3 | 98,3 | 68,7 | 33,5 | 100,0 | 59,3 | 75,4 | 78,1 | 56,6 |
| 75 min | 100,0 | 90,2 | 88,6 | 59,3 | 38,0 | 100,0 | 63,2 | 69,5 | 69,1 | 61,7 |
| 80 min | 100,0 | 80,2 | 100,0 | 64,1 | 24,1 | 100,0 | 63,3 | 63,3 | 64,1 | 74,1 |

| **cytokinesis** | | | | | | | | | | |
|---|---|---|---|---|---|---|---|---|---|---|
| 0 min | 15,4 | 79,9 | 27,8 | 6,2 | 33,6 | 39,0 | 42,5 | 49,5 | 71,5 | 26,6 |
| 2 min | 13,8 | 79,1 | 20,3 | -12,7 | 12,8 | 51,9 | 41,1 | 45,7 | 80,8 | 11,3 |
| 4 min | 5,7 | 80,0 | 17,5 | 4,8 | 7,9 | 61,2 | 51,1 | 42,5 | 72,4 | 3,0 |
| 6 min | 12,1 | 76,4 | 18,0 | 12,3 | 14,1 | 36,1 | 51,6 | 60,9 | 79,4 | 6,1 |
| 8 min | 1,1 | 80,7 | 14,9 | 7,4 | 20,0 | 37,4 | 54,5 | 56,2 | 69,4 | 2,6 |
| 10 min | | | | 22,8 | 19,7 | 56,4 | 58,5 | 65,0 | | 22,6 |

### Bem1-CCG x $N_{ub}$ Ste20

| **cell fusion** | **cell 1** | **cell 2** | **cell 3** | **cell 4** | **cell 5** | **cell 6** | **cell 7** | **cell 8** | **cell 9** | **cell 10** |
|---|---|---|---|---|---|---|---|---|---|---|
| 0 min | 0,0 | 0,0 | 0,0 | | 0,0 | 0,0 | 0,0 | 0,0 | 0,0 | |
| 5 min | -6,8 | 52,5 | 50,3 | | 21,4 | 6,7 | 8,5 | 67,7 | 18,9 | |
| 10 min | -4,3 | 57,9 | 83,9 | | 50,9 | 24,6 | 45,9 | 69,6 | 47,8 | |
| 15 min | 57,7 | 63,0 | 40,8 | | 8,6 | 50,4 | 31,5 | 76,9 | 100,0 | |
| 20 min | 60,6 | 100,0 | 33,5 | | 5,8 | 38,3 | 41,3 | 80,9 | 100,0 | |
| 25 min | 58,9 | 100,0 | 47,8 | | | 23,9 | 27,9 | 79,1 | 87,5 | |
| **bud growth** | | | | | | | | | | |
| 50 min | 71,1 | 63,5 | 88,0 | 25,9 | 55,7 | 70,3 | 92,2 | 94,4 | 87,5 | |
| 55 min | 100,0 | 77,9 | 83,9 | 34,7 | 57,5 | 86,9 | 97,4 | 92,4 | 82,7 | |
| 60 min | 100,0 | 76,1 | 79,5 | 53,1 | 55,5 | 74,9 | 100,0 | 100,0 | 100,0 | |
| 65 min | 100,0 | 87,9 | 92,1 | 74,0 | 22,9 | 79,5 | 100,0 | 99,7 | 80,6 | |
| 70 min | 100,0 | 92,2 | 100,0 | 72,2 | 28,9 | 70,8 | 100,0 | 99,5 | 91,5 | |
| 75 min | 73,7 | 83,0 | | 76,8 | 50,7 | 68,2 | 100,0 | 97,5 | 76,2 | |
| 80 min | 62,9 | 74,8 | | 82,5 | 59,0 | 77,8 | 100,0 | 94,1 | 82,8 | |
| **cytokinesis** | | | | | | | | | | |
| 0 min | 23,3 | | 25,8 | 20,9 | 51,5 | 57,5 | | 43,9 | 46,0 | 40,2 |
| 2 min | 27,2 | | 33,5 | 13,4 | 47,7 | 54,8 | 58,3 | 53,4 | 35,0 | 52,5 |
| 4 min | 47,5 | 23,0 | 25,4 | 5,5 | 47,5 | 48,3 | 67,0 | 53,2 | 42,3 | 43,3 |
| 6 min | 58,7 | 29,4 | 42,9 | 15,4 | 69,3 | | 75,9 | 50,3 | 38,6 | 46,9 |
| 8 min | 65,1 | 39,3 | 40,0 | 23,7 | 64,5 | | 78,7 | 66,0 | | |
| 10 min | 55,2 | 38,1 | 49,9 | 33,3 | 67,5 | | 73,5 | | | 76,0 |

### Bem1-CCG x $N_{ub}$-Ptc1

| **cell fusion** | **cell 1** | **cell 2** | **cell 3** | **cell 4** | **cell 5** | **cell 6** | **cell 7** | **cell 8** | **cell 9** | **cell 10** |
|---|---|---|---|---|---|---|---|---|---|---|
| 0 min | 0,0 | 0,0 | 0,0 | 0,0 | 0,0 | 0,0 | 0,0 | 0,0 | 0,0 | |
| 5 min | -14,6 | -0,7 | -11,2 | 25,1 | 25,1 | -3,4 | 4,5 | -2,3 | 7,7 | |
| 10 min | -6,3 | 16,1 | -26,1 | 11,5 | 47,6 | -17,5 | -8,1 | -1,6 | -11,3 | |
| 15 min | -34,4 | 3,3 | -53,9 | 46,8 | 7,9 | -11,8 | -4,5 | 52,3 | -29,2 | |
| 20 min | -16,2 | 7,0 | | | 49,9 | -97,3 | -8,2 | -21,8 | -87,8 | |
| 25 min | | -56,4 | | -21,9 | 42,7 | | | 12,9 | | |
| **bud growth** | | | | | | | | | | |
| 50 min | 22,8 | -56,4 | -116,7 | -21,9 | 70,1 | -21,8 | 26,1 | 12,9 | 19,6 | |
| 55 min | 29,0 | -70,4 | -105,1 | 12,7 | 22,8 | -9,6 | 47,8 | -6,9 | -1,7 | |
| 60 min | 21,7 | -94,6 | -90,9 | 41,9 | 19,1 | 1,2 | 35,1 | -17,4 | 33,7 | |
| 65 min | 15,3 | -93,5 | -129,2 | 20,2 | 16,5 | 23,6 | 28,9 | -53,1 | 34,7 | |
| 70 min | 32,4 | -114,8 | -105,3 | 19,7 | | 34,5 | 13,3 | 21,8 | 70,8 | |
| 75 min | 4,1 | | -114,6 | 13,5 | | 39,8 | 10,3 | | 54,1 | |
| 80 min | 10,9 | -65,8 | -111,0 | -8,6 | | 13,1 | 1,9 | 24,7 | 39,8 | |

| cytokinesis | | | | | | |
|---|---|---|---|---|---|---|
| 0 min | -5,9 | 13,6 | 22,9 | 10,8 | -3,8 | -8,5 |
| 2 min | -28,1 | 35,6 | -6,7 | -3,8 | 3,5 | 0,0 |
| 4 min | -9,1 | 20,2 | -8,1 | -10,1 | -48,2 | 2,7 |
| 6 min | -10,2 | 20,3 | -21,8 | 1,6 | 40,2 | -10,1 |
| 8 min | -14,0 | | -11,0 | 7,2 | 34,6 | 0,2 |
| 10 min | -17,9 | | -3,3 | -9,6 | 36,6 | -1,6 |

### Bem1-CCG x $N_{ub}$-Cdc24

| cell fusion | cell 1 | cell 2 | cell 3 | cell 4 | cell 5 | cell 6 | cell 7 | cell 8 | cell 9 | cell 10 |
|---|---|---|---|---|---|---|---|---|---|---|
| 0 min | 0,0 | 0,0 | 0,0 | | 0,0 | | 0,0 | 0,0 | 0,0 | |
| 5 min | -18,5 | 0,2 | 70,1 | | 24,4 | | 15,6 | 8,5 | 22,4 | |
| 10 min | 26,4 | 36,9 | 100,0 | | 55,2 | | 23,9 | 19,8 | 54,4 | |
| 15 min | 82,0 | 65,4 | 62,5 | | 56,5 | | 37,9 | 2,2 | 75,6 | |
| 20 min | 98,8 | 55,2 | | 75,3 | 98,5 | | 45,2 | 21,8 | 100,0 | |
| 25 min | | 38,2 | | | 100,0 | | 100,0 | 18,9 | | |
| **bud growth** | | | | | | | | | | |
| 50 min | 100,0 | 60,6 | 13,2 | 75,3 | 100,0 | | 76,8 | 100,0 | 100,0 | |
| 55 min | 100,0 | 42,5 | 22,9 | 96,4 | 100,0 | 13,5 | 81,5 | 100,0 | 100,0 | |
| 60 min | 100,0 | 91,6 | 82,0 | 100,0 | 98,0 | 18,0 | 56,4 | 100,0 | 100,0 | |
| 65 min | 100,0 | 100,0 | 100,0 | 100,0 | 61,5 | 100,0 | 44,0 | 100,0 | 100,0 | |
| 70 min | 100,0 | 100,0 | 100,0 | 74,3 | 62,2 | 66,6 | 25,0 | 100,0 | 100,0 | |
| 75 min | 100,0 | 100,0 | 100,0 | | 53,9 | 100,0 | 48,1 | 99,1 | 100,0 | |
| 80 min | 100,0 | 91,7 | 99,2 | | 40,3 | 100,0 | 55,2 | 100,0 | 100,0 | |
| **cytokinesis** | | | | | | | | | | |
| 0 min | 75,5 | 58,0 | 83,6 | 66,3 | 93,4 | | | | | |
| 2 min | 64,9 | 66,2 | 77,3 | 63,6 | 85,2 | | | | | |
| 4 min | 70,7 | 56,8 | 77,2 | 50,2 | 86,2 | | | | | |
| 6 min | 69,3 | 68,6 | 83,2 | 95,6 | 74,5 | | | | | |
| 8 min | 79,3 | 77,6 | 88,4 | 75,7 | 81,6 | | | | | |
| 10 min | 78,9 | 85,5 | 84,7 | 73,4 | 87,1 | | | | | |

### Bem1-CCG x $N_{ub}$-Cdc42

| cell fusion | cell 1 | cell 2 | cell 3 | cell 4 | cell 5 | cell 6 | cell 7 | cell 8 | cell 9 | cell 10 |
|---|---|---|---|---|---|---|---|---|---|---|
| 0 min | 0,0 | 0,0 | 0,0 | 0,0 | 0,0 | 0,0 | 0,0 | 0,0 | 0,0 | 0,0 |
| 5 min | -4,3 | | 12,9 | 19,8 | | -18,5 | -11,7 | 10,3 | | 14,5 |
| 10 min | -19,4 | -22,7 | 4,2 | 14,0 | 4,9 | 6,8 | 2,1 | 67,3 | | 14,4 |
| 15 min | | -14,1 | 35,8 | 26,3 | 12,6 | 17,5 | 23,3 | 100,0 | | 11,1 |
| 20 min | 17,8 | 100,0 | 100,0 | 26,0 | 4,6 | 21,3 | 100,0 | | | 29,2 |
| 25 min | 25,4 | 100,0 | | | -36,1 | 20,9 | 78,3 | | | 100,0 |

| | | | | | | | | | | |
|---|---|---|---|---|---|---|---|---|---|---|
| **bud growth** | | | | | | | | | | |
| 50 min | 17,8 | 22,2 | 24,0 | 53,5 | 39,7 | 37,1 | 39,8 | 100,0 | 36,2 | 39,5 |
| 55 min | 25,4 | 18,3 | 9,2 | 51,3 | 35,5 | 38,9 | 57,4 | 100,0 | 39,7 | 56,3 |
| 60 min | 28,0 | 31,0 | -15,0 | 53,0 | 48,0 | 45,6 | 100,0 | 95,0 | 54,1 | 54,5 |
| 65 min | 32,7 | 45,1 | 35,5 | 56,3 | 58,5 | 48,4 | 100,0 | 100,0 | 45,0 | 59,3 |
| 70 min | 35,9 | 37,5 | 45,8 | 57,1 | 59,8 | 58,6 | 100,0 | 94,2 | 45,9 | 61,0 |
| 75 min | 54,8 | 27,9 | 71,9 | 47,9 | 76,4 | 62,1 | 100,0 | 100,0 | 48,0 | 63,2 |
| 80 min | 50,8 | 22,4 | 69,4 | 53,9 | 64,9 | 65,4 | 100,0 | 99,3 | 60,1 | 71,1 |
| **cytokinesis** | | | | | | | | | | |
| 0 min | -5,8 | 12,7 | 18,6 | 19,2 | 13,5 | -30,5 | 14,6 | 85,6 | | |
| 2 min | | 10,3 | -0,8 | -6,4 | 14,2 | -10,7 | 10,5 | 64,0 | | |
| 4 min | -2,3 | 19,1 | 14,2 | -1,8 | 11,7 | -5,3 | 8,5 | 54,7 | | |
| 6 min | 2,3 | 21,2 | 4,9 | -22,9 | 13,8 | 11,6 | 10,8 | 74,6 | | |
| 8 min | 0,0 | 24,8 | 19,6 | -13,5 | 25,3 | -8,4 | 10,7 | 96,0 | | |
| 10 min | 10,7 | 29,6 | | -8,1 | 23,7 | -5,9 | 14,9 | 76,6 | | |

### Bem1-CCG x $N_{ub}$-Rga2

| **cell fusion** | **cell 1** | **cell 2** | **cell 3** | **cell 4** | **cell 5** | **cell 6** | **cell 7** | **cell 8** | **cell 9** | **cell 10** |
|---|---|---|---|---|---|---|---|---|---|---|
| 0 min | 0,0 | 0,0 | 0,0 | 0,0 | 0,0 | 0,0 | 0,0 | | | |
| 5 min | -12,0 | 1,1 | 28,3 | -9,5 | 12,6 | -7,1 | 4,8 | | | |
| 10 min | -30,1 | 5,3 | -2,5 | -13,0 | 19,9 | -14,9 | -8,9 | | | |
| 15 min | -17,5 | -43,6 | -23,2 | -43,6 | 45,0 | -30,9 | -47,7 | | | |
| 20 min | -16,3 | -51,5 | -12,8 | | 22,1 | -34,9 | | | | |
| 25 min | -39,6 | -44,6 | | 32,4 | -23,3 | | 7,9 | | | |
| **bud growth** | | | | | | | | | | |
| 50 min | 19,2 | 20,7 | 33,9 | 32,4 | 17,5 | 32,4 | -8,3 | | | |
| 55 min | 10,7 | 0,8 | 25,2 | 31,7 | 14,5 | -0,6 | 6,0 | | | |
| 60 min | 23,4 | 27,1 | 45,6 | 32,0 | -4,1 | 22,0 | -1,2 | | | |
| 65 min | 42,5 | 25,5 | 48,5 | 35,4 | 22,7 | 12,9 | 19,5 | | | |
| 70 min | 29,1 | 36,6 | 51,5 | 51,1 | 31,5 | 50,7 | 12,7 | | | |
| 75 min | 90,5 | 36,8 | 49,7 | 48,5 | 44,7 | 77,7 | 10,8 | | | |
| 80 min | 100,0 | 47,6 | 27,8 | 50,0 | 48,8 | 48,8 | 48,3 | | | |
| **cytokinesis** | | | | | | | | | | |
| 0 min | 13,7 | -19,4 | 11,6 | -7,1 | 2,0 | | | | | |
| 2 min | 11,3 | -18,9 | 6,4 | -18,4 | 7,7 | | | | | |
| 4 min | 19,1 | -14,2 | 9,2 | -16,9 | 12,5 | | | | | |
| 6 min | 18,9 | -19,1 | 9,2 | -15,3 | 17,3 | | | | | |
| 8 min | 6,7 | -14,2 | 3,0 | -11,0 | 6,2 | | | | | |
| 10 min | | -15,6 | 5,4 | | 13,0 | | | | | |

### Bem1-CCG x $N_{ub}$-Rsr1

| cell fusion | cell 1 | cell 2 | cell 3 | cell 4 | cell 5 | cell 6 | cell 7 | cell 8 | cell 9 | cell 10 |
|---|---|---|---|---|---|---|---|---|---|---|
| 0 min | 0,0 | 0,0 | 0,0 | 0,0 | 0,0 | 0,0 | 0,0 | 0,0 | 0,0 | 0,0 |
| 5 min | -18,2 | 17,2 | -35,3 | -16,8 | 69,5 | 9,9 | -0,1 | 5,3 | -23,8 | -10,6 |
| 10 min | -55,6 | 19,9 | -6,6 | -20,5 | 100,0 | 11,1 | -23,9 | 3,6 | -14,5 | -19,2 |
| 15 min | -117,5 | 51,5 | | -48,2 | | -9,6 | -26,8 | -15,0 | -27,6 | 56,2 |
| 20 min | -101,5 | 100,0 | | -64,3 | | -17,4 | -8,5 | -14,3 | -11,0 | 100,0 |
| 25 min | | | | | | | 37,6 | -20,5 | -39,5 | |
| **bud growth** | | | | | | | | | | |
| 50 min | -5,8 | 29,7 | 4,4 | 8,4 | 4,6 | 70,5 | 45,6 | 29,7 | -22,5 | 50,7 |
| 55 min | -6,3 | 53,9 | 11,5 | 25,6 | 19,4 | 83,1 | 40,0 | 27,5 | -10,8 | 78,7 |
| 60 min | 30,0 | 87,2 | 2,5 | 32,7 | 40,4 | 83,3 | 36,8 | 60,0 | -2,9 | 100,0 |
| 65 min | 27,8 | 100,0 | -3,1 | 18,3 | 37,2 | 88,8 | 53,1 | 50,9 | 25,6 | 68,4 |
| 70 min | 52,7 | 71,4 | -22,8 | 22,5 | 36,6 | 82,8 | 36,8 | 61,2 | 21,8 | 75,1 |
| 75 min | 61,6 | 87,4 | 6,0 | 2,0 | 29,8 | 57,9 | 22,6 | 48,2 | -6,2 | 85,1 |
| 80 min | 78,9 | 73,5 | 7,7 | -0,9 | 51,7 | 62,5 | 17,9 | 10,7 | 35,1 | 79,9 |
| **cytokinesis** | | | | | | | | | | |
| 0 min | | 1,1 | -13,2 | | -11,8 | -4,6 | -26,4 | -15,9 | | |
| 2 min | -16,1 | 2,5 | -11,6 | -22,9 | -4,1 | -14,3 | -19,4 | -15,6 | | |
| 4 min | | -15,7 | -12,7 | -23,8 | 1,9 | -11,1 | -14,8 | -5,6 | | |
| 6 min | -26,0 | -0,6 | -1,0 | -30,8 | 17,4 | -6,2 | -17,4 | -2,1 | | |
| 8 min | | 6,6 | -11,1 | -17,2 | 6,7 | -10,9 | -13,0 | -30,5 | | |
| 10 min | | -5,3 | -4,3 | -9,2 | | | -13,9 | | | |

### Bem1-CCG x $N_{ub}$-Bud6

| cell fusion | cell 1 | cell 2 | cell 3 | cell 4 | cell 5 | cell 6 | cell 7 | cell 8 | cell 9 | cell 10 |
|---|---|---|---|---|---|---|---|---|---|---|
| 0 min | 0,0 | 0,0 | 0,0 | 0,0 | 0,0 | 0,0 | 0,0 | | | |
| 5 min | -16,4 | | -12,5 | 25,9 | 4,4 | -19,3 | -1,5 | | | |
| 10 min | -21,8 | | -11,2 | 0,7 | | -29,3 | -4,2 | | | |
| 15 min | -42,0 | | 19,5 | -16,3 | | -11,7 | -41,1 | | | |
| 20 min | -52,0 | 56,5 | -5,8 | | | | -26,7 | | | |
| 25 min | | | | | | | | | | |
| **bud growth** | | | | | | | | | | |
| 50 min | 27,1 | 12,1 | 52,1 | -17,4 | 12,7 | 44,1 | 44,0 | | | |
| 55 min | 71,5 | 6,9 | 53,2 | -7,9 | 50,5 | 35,1 | 66,6 | | | |
| 60 min | 100,0 | 4,8 | 75,1 | 1,9 | 43,5 | 23,4 | 91,6 | | | |
| 65 min | 100,0 | 2,4 | 100,0 | 24,5 | 42,7 | 66,9 | 100,0 | | | |
| 70 min | 100,0 | 5,8 | 100,0 | -36,6 | 37,3 | 68,4 | 94,4 | | | |

| | | | | | | | |
|---|---|---|---|---|---|---|---|
| 75 min | 100,0 | 6,5 | 100,0 | -19,1 | 36,1 | 85,4 | 100,0 |
| 80 min | 100,0 | 13,9 | | -21,8 | 29,5 | 99,1 | 66,0 |
| **cytokinesis** | | | | | | | |
| 0 min | 33,4 | 0,0 | 1,6 | 11,4 | -12,8 | 18,7 | 0,0 |
| 2 min | 5,1 | | | -8,5 | -10,3 | 23,5 | 7,2 |
| 4 min | 1,9 | -1,6 | -11,3 | 6,8 | -12,9 | 18,7 | 13,2 |
| 6 min | -3,1 | -7,0 | 5,0 | 7,2 | -11,3 | 23,5 | 14,0 |
| 8 min | -0,7 | -7,4 | -18,2 | 1,1 | -11,9 | 32,1 | 21,0 |
| 10 min | -1,4 | -13,5 | -20,2 | 13,4 | -4,9 | 34,2 | 14,8 |

### Bem1-CCG x $N_{ub}$-Nba1

| cell fusion | cell 1 | cell 2 | cell 3 | cell 4 | cell 5 | cell 6 | cell 7 | cell 8 | cell 9 | cell 10 |
|---|---|---|---|---|---|---|---|---|---|---|
| 0 min | 0,0 | 0,0 | 0,0 | 0,0 | 0,0 | 0,0 | | | | |
| 5 min | -26,8 | -27,5 | -30,6 | -11,7 | 14,1 | 0,8 | | | | |
| 10 min | -43,6 | -32,3 | -46,9 | 36,7 | 30,0 | -5,4 | | | | |
| 15 min | -27,1 | -39,4 | -51,6 | 100,0 | 25,6 | 21,3 | | | | |
| 20 min | -41,6 | -69,2 | -22,9 | 49,2 | 29,9 | 70,0 | | | | |
| 25 min | -37,9 | 15,2 | -3,8 | 47,8 | -33,7 | | | | | |
| **bud growth** | | | | | | | | | | |
| 50 min | 6,5 | -1,4 | -0,6 | -9,6 | | 1,2 | | | | |
| 55 min | 0,8 | -4,4 | | 21,5 | | -35,1 | | | | |
| 60 min | 17,0 | -31,7 | | 17,0 | | -7,4 | | | | |
| 65 min | 2,1 | -28,2 | | 41,1 | | -23,8 | | | | |
| 70 min | 41,0 | -35,4 | 18,8 | | | 4,4 | | | | |
| 75 min | 31,9 | -50,7 | 33,0 | | | 0,2 | | | | |
| 80 min | | | | | | | | | | |
| **cytokinesis** | | | | | | | | | | |
| 0 min | 34,4 | 9,4 | 11,6 | 10,3 | | | 31,4 | 38,1 | 6,9 | |
| 2 min | 68,3 | 22,8 | 21,0 | 9,7 | 3,2 | 29,7 | 13,8 | 33,0 | 14,4 | |
| 4 min | 40,6 | 22,2 | 20,7 | 15,9 | 4,7 | 37,5 | 6,1 | 40,0 | 17,7 | |
| 6 min | 86,2 | 24,8 | 23,0 | 16,5 | 21,8 | 44,1 | 15,0 | 31,2 | 32,5 | |
| 8 min | 62,1 | 31,8 | 25,3 | 23,2 | 37,2 | 49,9 | 36,9 | 38,1 | 31,9 | |
| 10 min | 78,7 | | 20,8 | 27,0 | 28,9 | 58,2 | 26,8 | 43,1 | 26,0 | |

### Bem1-CCG x $N_{ub}$-Bem1

| cell fusion | cell 1 | cell 2 | cell 3 | cell 4 | cell 5 | cell 6 | cell 7 | cell 8 | cell 9 | cell 10 |
|---|---|---|---|---|---|---|---|---|---|---|
| 0 min | 0,0 | 0,0 | 0,0 | 0,0 | 0,0 | 0,0 | 0,0 | 0,0 | 0,0 | 0,0 |
| 5 min | -44,8 | -9,5 | -91,5 | 1,2 | -5,9 | -19,1 | -25,6 | -28,8 | | |
| 10 min | -2,5 | 14,6 | -75,3 | -16,8 | 0,4 | | -13,9 | | | 4,7 |
| 15 min | 28,1 | -10,4 | -45,2 | -11,9 | 17,2 | | | | | -18,8 |

| | | | | | | | | | | |
|---|---|---|---|---|---|---|---|---|---|---|
| 20 min | | | -54,3 | -33,6 | | | | | | -37,3 |
| 25 min | | | 13,1 | -43,3 | -19,0 | | | | | -19,1 |
| **bud growth** | | | | | | | | | | |
| 50 min | 38,1 | -10,8 | | 85,4 | 40,2 | -3,8 | -14,1 | -9,8 | -8,5 | -41,5 |
| 55 min | 57,9 | -29,7 | | 100,0 | 53,0 | 34,9 | 4,9 | -24,0 | 7,8 | -16,3 |
| 60 min | 97,7 | -2,6 | | 100,0 | 49,9 | 71,2 | 5,0 | 1,8 | 6,0 | -4,9 |
| 65 min | 100,0 | 2,3 | | | 53,4 | 65,1 | 16,9 | -6,0 | | 33,1 |
| 70 min | 73,2 | 6,9 | | | 59,4 | 55,0 | 6,3 | -8,5 | 72,9 | 98,7 |
| 75 min | 100,0 | 35,0 | | | 47,0 | 100,0 | -19,5 | -6,9 | 100,0 | 75,3 |
| 80 min | 100,0 | 30,5 | | | 42,6 | 100,0 | -19,3 | 22,4 | 100,0 | 77,4 |
| **cytokinesis** | | | | | | | | | | |
| 0 min | 46,3 | 22,8 | 43,1 | | 20,9 | 19,4 | | | | |
| 2 min | 42,9 | 24,9 | 33,0 | | 17,8 | 22,5 | 8,0 | | | |
| 4 min | 36,6 | 20,3 | 33,6 | | 22,0 | 24,1 | 17,0 | | | |
| 6 min | 39,6 | 22,0 | 31,2 | | 20,9 | 21,4 | 20,6 | | | |
| 8 min | 36,2 | 26,6 | 43,6 | | 12,9 | 22,5 | 25,1 | | | |
| 10 min | 34,8 | 21,1 | 39,6 | | 19,7 | 27,3 | 16,5 | | | |

### Bem1-CCG x $N_{ub}$-Exo70

| **cell fusion** | **cell 1** | **cell 2** | **cell 3** | **cell 4** | **cell 5** | **cell 6** | **cell 7** | **cell 8** | **cell 9** | **cell 10** |
|---|---|---|---|---|---|---|---|---|---|---|
| 0 min | 0,0 | 0,0 | 0,0 | 0,0 | 0,0 | 0,0 | | | | |
| 5 min | 8,0 | 6,9 | 6,2 | 30,2 | -20,1 | 5,3 | | | | |
| 10 min | 21,4 | 27,5 | -30,2 | 18,9 | -15,8 | 8,1 | | | | |
| 15 min | -14,4 | -34,6 | -66,0 | 2,0 | | -2,7 | | | | |
| 20 min | 15,7 | | -22,9 | -29,3 | | 7,9 | | | | |
| 25 min | | | | | | | | | | |
| **bud growth** | | | | | | | | | | |
| 50 min | 9,2 | -10,8 | 37,6 | 56,0 | 4,6 | 18,7 | | | | |
| 55 min | 10,6 | 46,1 | 26,5 | 98,3 | 8,6 | 32,4 | | | | |
| 60 min | 34,1 | 57,6 | 29,5 | 71,5 | 14,9 | 38,8 | | | | |
| 65 min | 24,1 | 56,9 | 55,0 | 70,8 | 37,0 | 32,7 | | | | |
| 70 min | 43,8 | 61,8 | 67,8 | 100,0 | 42,8 | 57,8 | | | | |
| 75 min | 48,0 | 78,8 | 92,3 | 62,0 | 20,4 | 49,7 | | | | |
| 80 min | 77,3 | 100,0 | 92,8 | 71,9 | 71,6 | 20,0 | | | | |
| **cytokinesis** | | | | | | | | | | |
| 0 min | 10,3 | -0,5 | 0,0 | 0,0 | 8,4 | 0,0 | 24,2 | | | |
| 2 min | 0,0 | -4,3 | 4,9 | 0,4 | 14,1 | 3,3 | 27,4 | | | |
| 4 min | -3,5 | -14,9 | -0,5 | -1,9 | 18,0 | 4,9 | 26,4 | | | |
| 6 min | -4,7 | 2,3 | 1,8 | 7,2 | 12,6 | 4,8 | 8,9 | | | |

| | | | | | | | |
|---|---|---|---|---|---|---|---|
| 8 min | -3,9 | 10,1 | 15,8 | 8,5 | 21,7 | 2,1 | 25,9 |
| 10 min | -11,0 | 5,7 | | 9,4 | 26,9 | 10,9 | 29,9 |

### Boi1-CCG x $N_{ub}$-Boi1

| cell fusion | cell 1 | cell 2 | cell 3 | cell 4 | cell 5 | cell 6 | cell 7 | cell 8 | cell 9 | cell 10 |
|---|---|---|---|---|---|---|---|---|---|---|
| 0 min | 0,0 | 0,0 | 0,0 | 0,0 | 0,0 | 0,0 | 0,0 | 0,0 | 0,0 | 0,0 |
| 5 min | -19,5 | 13,6 | 14,0 | 15,6 | -2,9 | -19,2 | 12,8 | 9,9 | 32,7 | -5,3 |
| 10 min | -9,3 | 11,1 | 24,8 | -2,5 | 5,7 | -39,9 | 5,3 | 4,8 | 29,3 | 6,8 |
| 15 min | -2,7 | 6,2 | 18,0 | 17,7 | 22,7 | -36,6 | 6,4 | 14,7 | 41,1 | 8,8 |
| 20 min | 38,2 | -4,1 | 8,2 | 9,9 | 5,1 | -26,2 | -12,9 | 13,4 | 43,4 | -8,7 |
| 25 min | -42,4 | -28,1 | -3,8 | 8,8 | -16,1 | -37,5 | 84,7 | 17,3 | 33,7 | 16,8 |
| **bud growth** | | | | | | | | | | |
| 30 min | 100,0 | 2,7 | 33,8 | 28,3 | 57,9 | 10,1 | 84,7 | 31,3 | 38,2 | 9,5 |
| 35 min | 100,0 | 13,4 | 53,6 | 33,4 | 66,6 | 4,6 | 72,2 | 60,1 | 19,9 | 55,8 |
| 40 min | 83,3 | 28,4 | 50,0 | 13,3 | 56,5 | 9,8 | 100,0 | 100,0 | 38,2 | 36,0 |
| 45 min | 100,0 | 31,3 | 54,1 | 18,0 | 78,0 | 25,6 | 100,0 | 81,3 | 29,8 | 41,8 |
| 50 min | 75,2 | 30,5 | 71,6 | 29,3 | 53,2 | 43,2 | 100,0 | 80,4 | 100,0 | 17,4 |
| 55 min | 60,2 | 22,2 | 56,4 | 44,7 | 55,5 | 45,6 | 100,0 | 100,0 | 100,0 | 25,4 |
| 60 min | 84,3 | 27,3 | 28,9 | 55,4 | 56,4 | 9,4 | 100,0 | 91,8 | 100,0 | 0,9 |
| **cytokinesis** | | | | | | | | | | |
| 117 min | | 22,5 | 20,0 | 6,2 | -6,7 | 47,1 | 9,3 | | | |
| 119 min | 14,2 | -1,2 | -9,1 | -22,6 | -37,2 | 29,0 | 14,4 | 6,0 | | |
| 121 min | -6,5 | -18,9 | -0,1 | -23,1 | -7,0 | 33,2 | 14,1 | 17,2 | | |
| 123 min | 5,1 | -0,6 | 5,0 | -27,2 | -23,8 | 37,9 | 4,3 | 33,0 | | |
| 125 min | 1,8 | -27,3 | -2,5 | -34,1 | -19,1 | 36,0 | 21,8 | 17,9 | | |
| 127 min | 9,4 | -21,9 | -18,0 | -39,4 | -19,8 | 41,2 | 15,6 | 10,0 | | |

### Boi1-CCG x $N_{ub}$-Nba1

| cell fusion | cell 1 | cell 2 | cell 3 | cell 4 | cell 5 | cell 6 | cell 7 | cell 8 | cell 9 | cell 10 |
|---|---|---|---|---|---|---|---|---|---|---|
| 0 min | 0,0 | 0,0 | 0,0 | 0,0 | 0,0 | 0,0 | 0,0 | 0,0 | 0,0 | |
| 5 min | -4,2 | -4,5 | 6,1 | -6,0 | -7,8 | -9,2 | 14,3 | -12,7 | 6,0 | |
| 10 min | -22,9 | -6,5 | 0,0 | -8,9 | -21,5 | -0,7 | 10,1 | -43,8 | 0,1 | |
| 15 min | -37,0 | -11,3 | -23,7 | -13,6 | -40,0 | -15,8 | 5,5 | -17,9 | -26,1 | |
| 20 min | -35,3 | -32,3 | -15,2 | | -52,7 | -35,3 | -6,4 | | -38,0 | |
| 25 min | -26,1 | -34,3 | | | | | 15,6 | | -18,1 | |
| **bud growth** | | | | | | | | | | |
| 30 min | -8,0 | -3,4 | -3,6 | 38,7 | -11,4 | | 26,0 | -1,3 | -3,1 | |
| 35 min | 16,9 | 7,8 | 10,9 | 3,9 | -25,0 | | 14,8 | 0,1 | 19,2 | |
| 40 min | 4,5 | 15,0 | -18,0 | 18,9 | -6,9 | | | 12,6 | | |
| 45 min | 0,4 | 17,3 | -20,6 | -9,7 | -1,6 | | 8,5 | 6,8 | 49,4 | |

| | | | | | | | | | |
|---|---|---|---|---|---|---|---|---|---|
| 50 min | -0,2 | 33,5 | -19,0 | -34,2 | -5,5 | | 8,4 | 12,6 | 56,6 |
| 55 min | -2,9 | 21,3 | -50,6 | | -13,1 | | 65,8 | -12,2 | |
| 60 min | | | | | | | | | |
| **cytokinesis** | | | | | | | | | |
| 117 min | 74,8 | 69,7 | 64,6 | 61,1 | 68,2 | | | | |
| 119 min | 77,5 | 66,6 | 66,9 | 63,3 | 65,9 | | | | |
| 121 min | 79,1 | 68,0 | 71,7 | 65,6 | 75,7 | | | | |
| 123 min | 83,4 | 68,4 | 71,8 | 69,3 | 78,6 | | | | |
| 125 min | 83,8 | 64,5 | 72,1 | 63,9 | 82,0 | | | | |
| 127 min | 80,5 | 69,2 | 73,2 | 72,4 | 86,0 | | | | |

### Boi1-CCG x $N_{ub}$-Ptc1

| **cell fusion** | **cell 1** | **cell 2** | **cell 3** | **cell 4** | **cell 5** | **cell 6** | **cell 7** | **cell 8** | **cell 9** | **cell 10** |
|---|---|---|---|---|---|---|---|---|---|---|
| 0 min | 0,0 | 0,0 | 0,0 | 0,0 | 0,0 | 0,0 | 0,0 | | | |
| 5 min | 0,4 | -12,2 | 40,9 | -26,2 | 17,4 | 4,5 | -1,0 | | | |
| 10 min | -30,5 | -1,9 | 30,5 | -16,7 | 20,8 | -19,4 | 38,2 | | | |
| 15 min | 2,9 | -30,7 | 43,0 | -46,6 | 9,7 | -11,4 | 7,2 | | | |
| 20 min | -30,2 | -9,7 | 3,5 | -54,7 | -8,6 | -46,3 | 22,2 | | | |
| 25 min | -52,9 | -60,4 | -22,6 | -16,7 | -15,4 | -47,5 | 27,5 | | | |
| **bud growth** | | | | | | | | | | |
| 30 min | -1,4 | -51,1 | -15,2 | 36,6 | 14,2 | -171,4 | | | | |
| 35 min | -26,4 | -39,4 | -19,8 | 58,4 | 76,5 | -126,9 | | | | |
| 40 min | -50,7 | -37,2 | 14,9 | 33,5 | 40,7 | -84,9 | | | | |
| 45 min | -46,6 | -14,5 | -5,1 | 35,8 | 87,8 | -122,7 | | | | |
| 50 min | -20,0 | -28,9 | 10,2 | 30,1 | 72,9 | -102,6 | | | | |
| 55 min | -52,1 | -25,4 | 9,6 | -15,8 | | | | | | |
| 60 min | | | | | | | | | | |
| **cytokinesis** | | | | | | | | | | |
| 117 min | -20,4 | -5,8 | 11,4 | 22,3 | -16,9 | -27,2 | | | | |
| 119 min | -15,4 | -27,9 | 8,3 | 1,1 | 15,4 | -13,5 | | | | |
| 121 min | -8,5 | -48,6 | 5,2 | -18,0 | 12,6 | -69,1 | | | | |
| 123 min | -11,8 | -55,5 | -22,9 | 0,9 | 21,4 | -7,4 | | | | |
| 125 min | -13,5 | -35,4 | -19,6 | -2,8 | -13,2 | -12,4 | | | | |
| 127 min | -14,5 | -48,6 | -9,5 | -3,9 | 6,5 | -37,5 | | | | |

### Epo1-CCG x $N_{ub}$-Boi1

| **cell fusion** | **cell 1** | **cell 2** | **cell 3** | **cell 4** | **cell 5** | **cell 6** | **cell 7** | **cell 8** | **cell 9** | **cell 10** |
|---|---|---|---|---|---|---|---|---|---|---|
| 0 min | 0,0 | 0,0 | 0,0 | 0,0 | 0,0 | 0,0 | 0,0 | 0,0 | 0,0 | 0,0 |
| 3 min | | 29,0 | -18,8 | -45,6 | -7,6 | -6,7 | 4,3 | -1,1 | | |
| 6 min | | 50,7 | -6,0 | -10,0 | -11,3 | 7,2 | 16,5 | -10,7 | | |

| | | | | | | | | | | |
|---|---|---|---|---|---|---|---|---|---|---|
| 9 min | | 21,6 | -11,2 | 0,3 | -13,5 | 7,6 | 6,5 | -4,2 | | |
| 12 min | | | -3,4 | -6,8 | -20,6 | 7,4 | 1,0 | -15,5 | | |
| 15 min | | | 4,5 | -5,9 | -22,8 | -3,6 | -0,6 | -5,1 | | |
| 18 min | | | -18,8 | -15,0 | -9,6 | -1,9 | -17,2 | -15,5 | | |
| 21 min | | | -2,2 | -12,2 | -14,4 | -9,5 | -13,7 | -11,6 | | |
| 24 min | | | | -12,6 | -17,2 | -28,2 | -9,5 | -19,8 | | |
| 27 min | | | | -11,0 | -13,3 | -10,6 | | -16,3 | | |
| **bud growth** | | | | | | | | | | |
| 30 min | -30,6 | 20,2 | -0,2 | -25,5 | 0,1 | -30,5 | | 8,3 | -0,9 | 24,0 |
| 33 min | | -33,2 | -13,7 | -22,7 | -6,9 | 10,3 | | 11,8 | -6,9 | 25,5 |
| 36 min | -74,5 | 47,2 | 2,6 | -21,8 | 28,1 | -21,4 | | 2,8 | -25,2 | 24,4 |
| 39 min | -24,4 | -2,4 | 3,2 | -13,9 | 32,9 | -13,3 | | -1,7 | -8,2 | 29,3 |
| 42 min | -57,7 | 63,9 | 0,8 | 1,2 | 26,6 | -13,7 | | 11,5 | -19,2 | 28,1 |
| 45 min | -24,0 | 70,2 | -1,6 | 19,4 | 27,9 | 12,6 | | 12,0 | -9,5 | 41,7 |
| 48 min | 10,5 | 93,4 | -16,0 | 16,3 | 17,5 | 17,9 | | 5,1 | -8,0 | 36,4 |
| 51 min | 53,6 | 100,0 | 2,9 | 38,3 | 29,2 | 17,9 | | 0,2 | -8,6 | 40,3 |
| 54 min | 31,0 | 100,0 | 27,6 | 33,5 | 8,4 | 12,5 | | -1,1 | -10,3 | 35,6 |
| **cytokinesis** | | | | | | | | | | |
| 117 min | -49,1 | 15,2 | -1,9 | -3,8 | | -15,7 | -22,3 | | | 26,2 |
| 120 min | -54,7 | 3,1 | -1,7 | -9,9 | | -14,0 | -16,5 | | | 24,8 |
| 123 min | -32,1 | -15,1 | -6,7 | 3,5 | | -12,6 | -8,7 | | | 25,3 |
| 126 min | -56,7 | 13,5 | -17,2 | 4,1 | | -24,8 | -17,4 | | | 25,6 |
| 129 min | -36,0 | 22,0 | 5,5 | -2,4 | | -12,2 | -16,7 | | | 26,9 |
| 132 min | -35,7 | 37,6 | -3,8 | 7,5 | | -16,5 | -4,0 | | | 25,9 |

### Epo1-CCG x $N_{ub}$-Ptc1

| cell fusion | cell 1 | cell 2 | cell 3 | cell 4 | cell 5 | cell 6 | cell 7 | cell 8 | cell 9 | cell 10 |
|---|---|---|---|---|---|---|---|---|---|---|
| 0 min | 0,0 | 0,0 | 0,0 | 0,0 | 0,0 | 0,0 | 0,0 | 0,0 | 0,0 | |
| 3 min | -1,3 | 27,6 | -14,8 | 0,0 | 20,0 | 1,7 | 18,0 | -0,6 | -6,7 | |
| 6 min | 2,2 | 49,8 | -0,7 | 4,8 | 50,6 | -9,9 | 25,0 | -17,4 | 7,2 | |
| 9 min | -10,6 | 26,5 | -16,8 | 51,3 | 56,4 | 5,3 | -74,9 | -17,2 | 7,6 | |
| 12 min | -5,5 | 23,3 | -31,0 | 3,1 | 13,3 | -1,7 | 73,1 | -23,0 | 7,4 | |
| 15 min | -6,1 | -14,8 | -19,3 | -26,6 | 21,3 | 1,1 | 53,8 | -4,5 | -3,6 | |
| 18 min | -26,7 | -0,7 | -16,6 | -7,4 | 58,1 | 0,0 | 18,5 | -20,3 | -1,9 | |
| 21 min | -9,2 | -16,8 | 2,0 | -24,5 | 58,0 | -1,1 | 3,0 | -13,1 | -9,5 | |
| 24 min | | -31,0 | -8,2 | -20,7 | 37,0 | | | -9,4 | -28,2 | |
| 27 min | | -19,3 | 26,5 | -6,7 | 47,5 | | | | -10,6 | |
| **bud growth** | | | | | | | | | | |
| 30 min | -0,8 | 41,0 | 23,3 | -18,3 | 37,0 | -32,9 | 34,6 | -37,2 | -30,5 | |
| 33 min | -12,8 | 50,9 | -14,8 | -4,5 | 47,5 | -49,3 | 7,0 | -11,6 | 10,3 | |

| | | | | | | | | | |
|---|---|---|---|---|---|---|---|---|---|
| 36 min | -26,6 | 45,3 | -0,7 | 4,8 | 10,0 | -34,8 | 45,3 | -16,9 | -21,4 |
| 39 min | -19,7 | 46,0 | -16,8 | -3,9 | 29,7 | -71,1 | 20,9 | -15,7 | -13,3 |
| 42 min | -26,5 | 50,9 | -31,0 | 11,7 | 67,6 | -60,3 | 3,9 | -6,7 | -13,7 |
| 45 min | -33,1 | 58,3 | -19,3 | 0,2 | -23,4 | -40,1 | 99,3 | -25,9 | 12,6 |
| 48 min | -32,1 | 42,9 | -16,6 | 8,5 | -42,2 | -35,8 | | -11,0 | 17,9 |
| 51 min | -56,7 | 65,2 | 2,0 | 21,4 | -30,0 | -10,7 | | -7,8 | 17,9 |
| 54 min | -43,7 | 63,2 | -8,2 | 31,5 | -39,2 | 13,9 | 22,6 | -4,4 | 12,5 |
| **cytokinesis** | | | | | | | | | |
| 117 min | -41,5 | 29,1 | 18,8 | -58,9 | -7,5 | | | -43,7 | -15,7 |
| 120 min | -36,6 | 21,8 | 23,3 | -100,5 | 14,8 | | | -33,9 | -14,0 |
| 123 min | -50,5 | 18,8 | 25,3 | -79,1 | -5,0 | | | -36,7 | -12,6 |
| 126 min | -40,1 | 23,3 | 19,5 | -108,7 | -1,9 | | | -45,1 | -24,8 |
| 129 min | -39,6 | 25,3 | 18,8 | -21,1 | -11,1 | | | -16,6 | -12,2 |
| 132 min | -39,0 | 19,5 | 23,0 | -20,2 | 0,3 | | | -18,5 | -16,5 |

**Table S2**

List of constructs used and created in this study.

| Plasmid | Description | Origin |
|---|---|---|
| pFA6a natNT2 | *natNT2, $Amp^R$* | Janke *et al.*, 2004 |
| pFA6a hphNT1 | *hphNT1, $Amp^R$* | Janke *et al.*, 2004 |
| pFA6a kanMX6 | *kanMX6, $Amp^R$* | Bähler *et al.*, 1998 |
| pFA6a CmLEU2 | *CmLEU2, $Amp^R$* | Schaub *et al.*, 2006 |
| pFA6a HIS3MX6 | *HISMX6, $Amp^R$* | Longtine *et al.*, 1998 |
| pRS 314 | *$Amp^R$, TRP1, CEN* | Sikorski and Hieter, 1989 |
| pRS 315 | *$Amp^R$, LEU2, CEN* | Sikorski and Hieter, 1989 |
| pRS 316 | *$Amp^R$, URA3, CEN* | Sikorski and Hieter, 1989 |
| pYM-N35 | *$P_{MET17}$, natNT2, $Amp^R$* | Janke et al., 2004 |
| $pP_{CUP1}N_{ub}$-HA kanMX4 | *$P_{CUP1}N_{ub}$-HA, $Amp^R$, kanMX4* | Dünkler *et al.*, 2012 |
| $pP_{CUP1}N_{ub}$-HA kanMX4 CEN | *$P_{CUP1}N_{ub}$-HA, $Amp^R$, kanMX4, CEN6/ARSH4* | Dünkler *et al.*, 2012 |
| pCRU 303 | *$C_{ub}$-R-URA3, $Amp^R$, HIS3* | Dünkler *et al.*, 2012 |
| pFA6a CRU HIS3MX6 | *$C_{ub}$-R-URA3, $Amp^R$, HISMX6* | This work |
| pBem1-CRU 303 | *BEM1-$C_{ub}$-R-URA3, $Amp^R$, HIS3* | This work |
| $pP_{MET}$ Bem1 CRU 313 | *$P_{MET17}$ BEM1-$C_{Ub}$-R-URA3, $Amp^R$, HIS3* | This work |
| $pP_{MET}$ $Bem1_{WK}$ CRU 313 | *$P_{MET17}$ $BEM1_{W192K}$-$C_{Ub}$-R-URA3, $Amp^R$, HIS3* | This work |
| pBud6-CRU 303 | *BUD6-$C_{ub}$-R-URA3, $Amp^R$, HIS3* | This work |
| $pP_{MET}$ $Bud6_{1-364}$ CRU 313 | *$P_{MET17}$ $BUD6_{1-364}$-$C_{ub}$-R-URA3, HIS3, CEN* | This work |
| $pP_{MET}$ $Bud6_{360-788}$ CRU 313 | *$P_{MET17}$ $BUD6_{360-788}$-$C_{ub}$-R-URA3, HIS3, CEN* | This work |
| pEpo1-CRU 303 | *EPO1-$C_{ub}$-R-URA3, $Amp^R$, HIS3* | Neller *et al.*, 2014 |
| pNba1-CRU 303 | *NBA1-$C_{ub}$-R-URA3, $Amp^R$, HIS3* | This work |
| pGps1-CRU 303 | *GPS1-$C_{ub}$-R-URA3, $Amp^R$, HIS3* | This work |
| $pP_{MET}$ $Gps1_{536-758}$ CRU 313 | *$P_{MET17}$ $GPS1_{536-758}$ -$C_{Ub}$-R-URA3, $Amp^R$, HIS3* | This work |
| pBoi1-CRU 303 | *BOI1-$C_{ub}$-R-URA3, $Amp^R$, HIS3* | Kustermann *et al.*, 2017 |
| pBoi2-CRU 303 | *BOI2-$C_{ub}$-R-URA3, $Amp^R$, HIS3* | Kustermann *et al.*, 2017 |
| pFir1-CRU 303 | *FIR1-$C_{ub}$-R-URA3, $Amp^R$, HIS3* | This work |
| pBem1-CCG 306 | *BEM1-mCHERRY-$C_{ub}$-R-GFP, $Amp^R$, URA3* | This work |
| pBoi1-CCG 306 | *BOI1-mCHERRY-$C_{ub}$-R-GFP, $Amp^R$, URA3* | Kustermann *et al.*, 2017 |
| pEpo1-CCG 306 | *EPO1-mCHERRY-$C_{ub}$-R-GFP, $Amp^R$, URA3* | Neller *et al.*, 2014 |
| $pGic2_{PBD}$-RFP | *$P_{GIC2}$-$GIC2_{1-208}$-RFP, $Amp^R$, His3, Ura3* | Tong et al., 2007 |
| $pP_{MET}Gic2_{PBD}$ 314 | *$P_{MET17}$-$GIC2_{1-208}$-9xMYC, $Amp^R$, TRP1* | This work |
| $pP_{MET}Gic2_{PBD}$ 315 | *$P_{MET17}$-$GIC2_{1-208}$-9xMYC, $Amp^R$, LEU2* | This work |
| $pGic2_{PBD}$-mCherry 306 | *$P_{GIC2}$-$GIC2_{1-208}$-mCHERRY, $Amp^R$, Ura3* | This work |
| $pP_{MET}Bem1$ 316 | *$P_{BEM1}$ BEM1, URA3, $Amp^R$, CEN* | This work |
| $pP_{MET}Bem1$ 313 | *$P_{MET17}$ GFP-BEM1, $Amp^R$, HIS3, CEN* | This work |
| $pP_{MET}Bem1_{1-414}$ 313 | *$P_{MET17}$ GFP-$bem1_{1-414}$, $Amp^R$, HIS3, CEN* | This work |
| $pP_{MET}Bem1_{145-551}$ 313 | *$P_{MET17}$ GFP-$bem1_{145-551}$, $Amp^R$, HIS3, CEN* | This work |

| | | |
|---|---|---|
| $pP_{MET}Bem1_{145\text{-}414}$ 313 | $P_{MET17}$ *GFP-bem1*$_{145\text{-}414}$, *Amp*$^R$, *HIS3, CEN* | This work |
| $pP_{MET}Bem1_{145\text{-}268}$ 313 | $P_{MET17}$ *GFP-bem1*$_{145\text{-}268}$, *Amp*$^R$, *HIS3, CEN* | This work |
| $pP_{MET}Bem1_{268\text{-}551}$ 313 | $P_{MET17}$ *GFP-bem1*$_{268\text{-}551}$, *Amp*$^R$, *HIS3, CEN* | This work |
| $pP_{MET}Bem1_{217\text{-}414}$ 313 | $P_{MET17}$ *GFP-bem1*$_{217\text{-}414}$, *Amp*$^R$, *HIS3, CEN* | This work |
| $pP_{MET}Bem1_{268\text{-}414}$ 313 | $P_{MET17}$ *GFP-bem1*$_{268\text{-}414}$, *Amp*$^R$, *HIS3, CEN* | This work |
| $pP_{MET}Bem1_{470\text{-}551}$ 313 | $P_{MET17}$ *GFP-bem1*$_{470\text{-}551}$, *Amp*$^R$, *HIS3, CEN* | This work |
| $pP_{MET}Bem1_{WK}$ 313 | $P_{MET17}$ *GFP-bem1*$_{W192K}$, *Amp*$^R$, *HIS3, CEN* | This work |
| $pP_{MET}Bem1_{ND}$ 313 | $P_{MET17}$ *GFP-bem1*$_{N253D}$, *Amp*$^R$, *HIS3, CEN* | This work |
| $pP_{MET}Bem1_{WK\ ND}$ 313 | $P_{MET17}$ *GFP-bem1*$_{W192K\ N253D}$, *Amp*$^R$, *HIS3, CEN* | This work |
| $pP_{MET}Bem1_{145\text{-}268\ WK}$ 313 | $P_{MET17}$ *GFP-bem1*$_{145\text{-}268\ WK}$, *Amp*$^R$, *HIS3, CEN* | This work |
| $pP_{MET}Bem1_{145\text{-}268\ ND}$ 313 | $P_{MET17}$ *GFP-bem1*$_{145\text{-}268\ ND}$, *Amp*$^R$, *HIS3, CEN* | This work |
| $pP_{MET}Bem1_{145\text{-}268\ WK\ ND}$ 313 | $P_{MET17}$ *GFP-bem1*$_{145\text{-}268\ WK\ ND}$, *Amp*$^R$, *HIS3, CEN* | This work |
| $pP_{MET}$Bem3-GFP 313 | $P_{MET17}$ *BEM3-GFP, Amp*$^R$, *HIS3, CEN* | This work |
| $pP_{MET}Bem3^{R950G}$-GFP 313 | $P_{MET17}$ *BEM3*$_{R950G}$*-GFP, Amp*$^R$, *HIS3, CEN* | This work |
| $pP_{MET}Bem3_{50\text{-}end}$-GFP 313 | $P_{MET17}$ *BEM3*$_{\mathbf{50\text{-}end}}$*-GFP, Amp*$^R$, *HIS3, CEN* | This work |
| $pP_{MET}Bem3_{101\text{-}end}$-GFP 313 | $P_{MET17}$ *BEM3*$_{\mathbf{101\text{-}end}}$*-GFP, Amp*$^R$, *HIS3, CEN* | This work |
| $pP_{MET}Bem3_{\Delta PX}$-GFP 313 | $P_{MET17}$ *BEM3*$_{\mathbf{\Delta 111\text{-}118}}$*-GFP, Amp*$^R$, *HIS3, CEN* | This work |
| $pP_{MET}Bem3_{GAP}$-GFP 313 | $P_{MET17}$ *BEM3*$_{\mathbf{751\text{-}end}}$*-GFP, Amp*$^R$, *HIS3, CEN* | This work |
| $pP_{MET}Bem3_{PHmut}$-GFP 313 | $P_{MET17}$ *BEM3*$_{R664S\ R645S\ K647D}$*-GFP, Amp*$^R$, *HIS3, CEN* | This work |
| $pP_{MET}$Bem3-GFP 315 | $P_{MET17}$ *BEM3-GFP, Amp*$^R$, *LEU2, CEN* | This work |
| $pP_{MET}Bem3_{R950G}$-GFP 315 | $P_{MET17}$ *BEM3*$_{R950G}$*-GFP, Amp*$^R$, *LEU2, CEN* | This work |
| pBem1 GFP 303 | $P_{BEM1}$ *BEM1-GFP, Amp*$^R$, *HIS3* | This work |
| $pBem1_{WK}$ GFP 303 | $P_{BEM1}$ *BEM1* $_{W192K}$*-GFP, Amp*$^R$, *HIS3* | This work |
| $pBoi1_{\Delta PxxP}$ 303 | $P_{BOI1}$ *BOI1*$_{\Delta 394\text{-}413}$, *Amp*$^R$, *HIS3* | This work |
| $pBoi1_{\Delta PxxP}$-GFP 303 | $P_{BOI1}$ *BOI1*$_{\Delta 394\text{-}413}$*-GFP, Amp*$^R$, *HIS3* | This work |
| pBoi1 303 | $P_{BOI1}$ *BOI1, Amp*$^R$, *HIS3* | This work |
| pBoi1-GFP 303 | $P_{BOI1}$ *BOI1-GFP, Amp*$^R$, *HIS3* | This work |
| pCla4 303 | $P_{CLA4}$ *CLA4, Amp*$^R$, *HIS3* | This work |
| $pCla4_{PPAA\ FL}$ 303 | $P_{CLA4}$ *CLA4*$_{F15A\ PP22.25AA\ F451L}$, *Amp*$^R$, *HIS3* | This work |
| $pCla4_{PPAA\ FL}$-GFP 303 | $P_{CLA4}$ *CLA4*$_{F15A\ PP22.25AA\ F451L}$*-GFP, Amp*$^R$, *HIS3* | This work |
| pCla4 GFP 303 | $P_{CLA4}$ *CLA4-GFP, Amp*$^R$, *HIS3* | This work |
| pSte20 303 | $P_{STE20}$ *STE20, Amp*$^R$, *HIS3* | This work |
| $pSte20_{PPAA}$ 303 | $P_{STE20}$ *STE20*$_{PP477.480AA}$, *Amp*$^R$, *HIS3* | This work |
| $pSte20_{FLPT}$ 303 | $P_{STE20}$ *STE20*$_{F470L\ P475T}$, *Amp*$^R$, *HIS3* | This work |
| $pP_{MET}$Ste20-CRU 313 | $P_{MET17}$ *STE20-C*$_{Ub}$*-R-URA3, Amp*$^R$, *HIS3* | This work |
| $pP_{MET}Ste20_{PPAA}$-CRU 313 | $P_{MET17}$ *STE20*$_{PP477.480AA}$*-C*$_{Ub}$*-R-URA3, Amp*$^R$, *HIS3* | This work |
| $pP_{MET}Ste20_{FLPT}$-CRU 313 | $P_{MET17}$ *STE20* $_{F470L\ P475T}$*-C*$_{Ub}$*-R-URA3, Amp*$^R$, *HIS3* | This work |
| pBem1-GFP 304 | *BEM1-GFP, Amp*$^R$, *TRP1* | This work |
| pCdc24-GFP 304 | *CDC24-GFP, Amp*$^R$, *TRP1* | This work |
| pNba1-GFP 304 | *NBA1-GFP, Amp*$^R$, *TRP1* | This work |
| pGps1-GFP 304 | *GPS1-GFP, Amp*$^R$, *TRP1* | This work |

| | | |
|---|---|---|
| pBoi1-GFP 304 | *BOI1-GFP, Amp*$^{R}$*, TRP1* | Kustermann *et al.*, 2017 |
| pP$_{MET}$SH3$_{Boi1}$-GFP 306 | *P*$_{MET17}$ *BOI1*$_{1\text{-}83}$*-GFP, Amp*$^{R}$*, URA3* | This work |
| pBoi2-GFP 304 | *BOI2-GFP, Amp*$^{R}$*, HIS3* | Kustermann *et al.*, 2017 |
| pP$_{MET}$SH3$_{Boi2}$-GFP 306 | *P*$_{MET17}$ *BOI2*$_{1\text{-}110}$*-GFP, Amp*$^{R}$*, URA3* | This work |
| pCla4-GFP 304 | *CLA4-GFP, Amp*$^{R}$*, TRP1* | This work |
| pSte20-GFP 304 | *STE20-GFP, Amp*$^{R}$*, TRP1* | This work |
| pShs1-GFP304 | *SHS1-GFP, Amp*$^{R}$*, TRP1* | This work |
| pMyo1-mCherry 306 | *MYO1-mCHERRY, Amp*$^{R}$*, URA3* | This work |
| pGEX-6PI | *P*$_{tac}$*GST, Amp*$^{R}$ | GE Healthcare |
| pGEX-6PI Bud6$_{1\text{-}364}$ | *P*$_{tac}$*GST-BUD6*$_{1\text{-}364}$*, Amp*$^{R}$ | This work |
| pGEX-6PI Bem1$_{140\text{-}271}$ | *P*$_{tac}$*GST-BEM1*$_{418\text{-}813}$*, Amp*$^{R}$ | This work |
| pGEX-6PI Nbp2 | *P*$_{tac}$*GST-NBP2, Amp*$^{R}$ | This work |
| pGEX-2T | *P*$_{tac}$*GST, Amp*$^{R}$ | GE Healthcare |
| pGEX-2T-SH3$_{Boi1}$ | *P*$_{tac}$*GST-BOI1*$_{1\text{-}154}$*, Amp*$^{R}$ | This work |
| pGEX-2T-SH3$_{Boi2}$ | *P*$_{tac}$*GST-BOI2*$_{1\text{-}160}$*, Amp*$^{R}$ | This work |
| pAc | *P*$_{T7}$*6HIS, Kan*$^{R}$ | Iffland *et al.*, 2000 |
| pAc Epo1$_{PxxP}$-SNAP | *P*$_{T7}$*-6his-EPO1*$_{640\text{-}670}$-SNAP, *Kan*$^{R}$ | This work |
| pAc Nba1$_{PxxP}$-SNAP | *P*$_{T7}$*-6his-NBA1*$_{202\text{-}289}$-SNAP, *Kan*$^{R}$ | |
| pAc Cla4$_{1\text{-}33}$-SNAP | *P*$_{T7}$*-6his-CLA4*$_{1\text{-}33}$-SNAP, *Kan*$^{R}$ | This work |
| pAc Cla4$_{1\text{-}33\ AA}$-SNAP | *P*$_{T7}$*-6his-CLA4*$_{1\text{-}33\ PP22.25AA}$-SNAP, *Kan*$^{R}$ | This work |
| pAc Cla4$_{1\text{-}33\ F15A\ AA}$-SNAP | *P*$_{T7}$*-6his-CLA4*$_{1\text{-}33\ F15A\ PP22.25AA}$-SNAP, *Kan*$^{R}$ | This work |
| pAc Cla4$_{1\text{-}33\ F15A}$-SNAP | *P*$_{T7}$*-6his-CLA4*$_{1\text{-}33\ F15A}$-SNAP, *Kan*$^{R}$ | This work |
| pAc Cla4$_{437\text{-}471}$-SNAP | *P*$_{T7}$*-6his-CLA4*$_{437\text{-}471}$-SNAP, *Kan*$^{R}$ | This work |
| pAc Cla4$_{437\text{-}471\ F451L}$-SNAP | *P*$_{T7}$*-6his-CLA4*$_{437\text{-}471\ F451L}$-SNAP, *Kan*$^{R}$ | This work |
| pP$_{MET}$Cla4$_{1\text{-}471}$-9xMyc 314 | *P*$_{MET17}$ *CLA4*$_{1\text{-}471}$ *9xMYC, Amp*$^{R}$*, TRP1* | This work |
| pP$_{MET}$Cla4$_{1\text{-}471\ F15A\ AA/AA}$-9xMyc 314 | *P*$_{MET17}$ *CLA4*$_{1\text{-}471\ F15A\ PP22.25AA\ PP458.461AA}$ *9xMYC, Amp*$^{R}$*, TRP1* | This work |
| pP$_{MET}$Cla4$_{1\text{-}471\ F15A\ AA/PP\ F451L}$-9xMyc 314 | *P*$_{MET17}$ *CLA4*$_{1\text{-}471\ F15A\ PP22.25AA\ F451L}$ *9xMYC, Amp*$^{R}$*, TRP1* | This work |
| pP$_{MET}$Ste20-9xMyc 314 | *P*$_{MET17}$ *STE20-9xMYC, Amp*$^{R}$*, TRP1* | This work |
| pP$_{MET}$Ste20$_{F470I\ P475T}$-9xMyc 314 | *P*$_{MET17}$ *STE20* $_{F470I\ P475T}$ *-9xMYC, Amp*$^{R}$*, TRP1* | This work |
| pP$_{MET}$Ste20$_{PP477.480AA}$-9xMyc 314 | *P*$_{MET17}$ *STE20* $_{PP477.480AA}$*-9xMYC, Amp*$^{R}$*, TRP1* | This work |
| pML107 | *P*$_{TDH3}$*SpCAS9, Amp*$^{R}$*, LEU2* | Laughery *et al.*, 2015 |
| pML107 Bem1-748 | *P*$_{SNR52}$*sgRNA-BEM1*$^{748}$*, P*$_{TDH3}$*SpCAS9, Amp*$^{R}$*, LEU2* | This work |
| pML107 Bem1-561 | *P*$_{SNR52}$*sgRNA-BEM1*$^{561}$*, P*$_{TDH3}$*SpCAS9, Amp*$^{R}$*, LEU2* | This work |
| pML107 Fir1-1573 | *P*$_{SNR52}$*sgRNA-FIR1*$^{1573}$*, P*$_{TDH3}$*SpCAS9, Amp*$^{R}$*, LEU2* | This work |
| pML107 Boi1-792 | *P*$_{SNR52}$*sgRNA-BOI1*$^{792}$*, P*$_{TDH3}$*SpCAS9, Amp*$^{R}$*, LEU2* | This work |
| pML107 Boi2-906 | *P*$_{SNR52}$*sgRNA-BOI2*$^{906}$*, P*$_{TDH3}$*SpCAS9, Amp*$^{R}$*, LEU2* | This work |

**Table S3**

List of *S. cerevisiae* strains used and created in this study.

| Strain | Name | Relevant genotype | Origin |
|---|---|---|---|
| JD47 | JD47 | MATa, *his3-Δ200, leu2-3, 112 lys2-801, trp1-Δ63, ura3-52* | Dohmen et al., 1995 |
| JD53 | JD53 | MATα, *his3-Δ200, leu2-3, 112 lys2-801, trp1-Δ63, ura3-52* | Dohmen et al., 1995 |
| JD51 | JD51 | MATa/α, *his3-Δ200/his3-Δ200, leu2-3/leu2-3, 112 lys2- 801/112 lys2-801, trp1-Δ63/trp1-Δ63, ura3-52/ura3-52* | Dohmen et al., 1995 |
| ULM53 | ULM53 | MATα, *his3-Δ200, leu2-3, 112 lys2-801, trp1-Δ63, ura3-52* | This work |
| NJY319 | Bem1CRU | JD47, *BEM1::BEM1-*$C_{ub}$*-R-URA3 HIS3* | This work |
| YAD3026 | Bem1$_{WK}$ | JD47, *BEM1::bem1*$_{W192K}$ | This work |
| STY1133 | Bem1$_{WK}$CRU | JD47, *BEM1::bem1*$_{W192K}$-$C_{ub}$*-R-URA3 HIS3* | This work |
| ARY41 | Bem1$_{\Delta PB}$CRU | JD47, *BEM1::bem1*$_{1\text{-}479}$-$C_{ub}$*-R-URA3 HIS3* | This work |
| STY1118 | Bem1CRU $N_{ub}$-Nba1 Δboi2 | 1n; *BEM1::BEM1-*$C_{ub}$*-R-URA3 LEU2,* $P_{NBA1}$::$P_{CUP1}$$N_{ub}$*-HA kanMX, BOI2::hphNT1,* | This work |
| STY1119 | Bem1CRU $N_{ub}$-Nba1 Δboi2 Boi1$_{\Delta PxxP}$ | 1n; *BEM1::BEM1-*$C_{ub}$*-R-URA3 LEU2,* $P_{NBA1}$::$P_{CUP1}$$N_{ub}$*-HA kanMX, BOI1::natNT2, BOI2::hphNT1,* $P_{BOI1}$::$P_{BOI1}$*boi1*$_{\Delta 394\text{-}413}$ *HIS3* | This work |
| STY1129 | Bem1CRU $N_{ub}$-Bem1 Δboi2 | 1n; p$P_{MET17}$ *BEM1-*$C_{ub}$*-R-URA3 CEN TRP1,* $P_{Bem1}$::$P_{CUP1}$$N_{ub}$*-HA kanMX, BOI2::hphNT1,* | This work |
| STY1130 | Bem1CRU $N_{ub}$-Bem1 Δboi2 Boi1$_{\Delta PxxP}$ | 1n; p$P_{MET17}$ *BEM1-*$C_{ub}$*-R-URA3 CEN TRP1,* $P_{BUD6}$::$P_{CUP1}$$N_{ub}$*-HA kanMX, BOI1::natNT2, BOI2::hphNT1,* $P_{BOI1}$::$P_{BOI1}$*boi1*$_{\Delta 394\text{-}413}$ *HIS3* | This work |
| STY1151 | Bem1CRU $N_{ub}$-Bud6 | JD47, *BEM1::BEM1-*$C_{ub}$*-R-URA3 LEU2,* $P_{BUD6}$::$P_{CUP1}$$N_{ub}$*-HA kanMX,* | This work |
| STY1149 | Bem1CRU $N_{ub}$-Bud6 Δboi2 | 1n; *BEM1::BEM1-*$C_{ub}$*-R-URA3 LEU2,* $P_{BUD6}$::$P_{CUP1}$$N_{ub}$*-HA kanMX, BOI2::hphNT1,* | This work |
| STY1150 | Bem1CRU $N_{ub}$-Bud6 Δboi2 Boi1$_{\Delta PxxP}$ | 1n; *BEM1::BEM1-*$C_{ub}$*-R-URA3 LEU2,* $P_{BUD6}$::$P_{CUP1}$$N_{ub}$*-HA kanMX, BOI1::natNT2, BOI2::hphNT1,* $P_{BOI1}$::$P_{BOI1}$*boi1*$_{\Delta 394\text{-}413}$ *HIS3* | This work |
| NJY355 | Bud6CRU | JD47, *BUD6::BUD6-*$C_{ub}$*-R-URA3 HIS3* | This work |
| YWY45 | Bud6$_{1\text{-}364}$CRU | JD47, p$P_{MET17}$ *BUD6*$_{1\text{-}364}$-$C_{ub}$*-R-URA3 CEN HIS3* | This work |
| YWY45 | Bud6$_{360\text{-}788}$CRU | JD47, p$P_{MET17}$ *BUD6*$_{360\text{-}788}$-$C_{ub}$*-R-URA3 CEN HIS3* | This work |
| UNY94 | Gps1CRU | JD47, *GPS1::GPS1-*$C_{ub}$*-R-URA3 HIS3* | This work |
| YWY57 | Gps1$_{536\text{-}758}$CRU | JD47, p$P_{MET17}$*GPS1*$_{536\text{-}758}$-$C_{Ub}$*-R-URA3 CEN HIS3* | This work |
| STY484 | Gps1$_{536\text{-}758}$CRU $N_{ub}$-Boi1 | JD53, p$P_{MET17}$*GPS1*$_{536\text{-}758}$-$C_{Ub}$*-R-URA3 CEN HIS3,* $P_{BOI1}$::$P_{CUP1}$$N_{ub}$*-HA kanMX* | This work |
| STY485 | Gps1$_{536\text{-}758}$CRU $N_{ub}$-Boi1 Δnba1 | JD53, p$P_{MET17}$*GPS1*$_{536\text{-}758}$-$C_{Ub}$*-R-URA3 CEN HIS3,* $P_{BOI1}$::$P_{CUP1}$$N_{ub}$*-HA kanMX, NBA1::natNT2* | This work |
| STY487 | Gps1$_{536\text{-}758}$CRU $N_{ub}$-Guk1 | JD47, p$P_{MET17}$*GPS1*$_{536\text{-}758}$-$C_{Ub}$*-R-URA3 CEN HIS3,* p$P_{CUP1}$$N_{ub}$*-HA-GUK1 CEN kanMX* | This work |
| STY487 | Gps1$_{536\text{-}758}$CRU $N_{ub}$-Guk1 Δnba1 | JD47, p$P_{MET17}$*GPS1*$_{536\text{-}758}$-$C_{Ub}$*-R-URA3 CEN HIS3,* p$P_{CUP1}$$N_{ub}$*-HA-GUK1 CEN kanMX, NBA1::natNT2* | This work |
| YWY63 | Nba1CRU | JD47, *NBA1::NBA1-*$C_{ub}$*-R-URA3 HIS3* | This work |

| | | | |
|---|---|---|---|
| NJY317 | Boi1CRU | JD47, *BOI1::BOI1-$C_{ub}$-R-URA3 HIS3* | Kustermann *et al.*, 2017 |
| NJY318 | Boi2CRU | JD47, *BOI2::BOI2-$C_{ub}$-R-URA3 HIS3* | Kustermann *et al.*, 2017 |
| STY1307 | Fir1CRU | JD47, *$P_{FIR1}$::$P_{MET17}$ natNT2, FIR1::FIR1-$C_{ub}$-R-URA3 HIS3* | This work |
| YAD3556 | Fir1$_{\Delta PxxP}$CRU | JD47, *$P_{FIR1}$:: $P_{MET17}$ natNT2, FIR1::fir1$_{PKRSPLR/AKSSLS}$-$C_{ub}$-R-URA3 HIS3* | This work |
| STY48 | Epo1CRU | JD47, *$P_{FIR1}$:: $P_{MET17}$ natNT2, EPO1::EPO1-$C_{ub}$-R-URA3 HIS3* | Neller *et al.*, 2014 |
| YAD2288 | Epo1$_{\Delta PxxP}$CRU | JD47, *$P_{FIR1}$:: $P_{MET17}$ natNT2, EPO1::epo1$_{\Delta 654\text{-}661}$-$C_{ub}$-R-URA3 HIS3* | This work |
| NJY168 | Cla4CRU | JD47, *CLA4::CLA4-$C_{ub}$-R-URA3 HIS3* | This work |
| NJY169 | Ste20CRU | JD47, *STE20::STE20-$C_{ub}$-R-URA3 HIS3* | This work |
| YAD1072 | $P_{MET}$Bem1-CCG | JD47, *$P_{BEM1}$:: $P_{MET17}$ natNT2, BEM1::BEM1-Cherry-$C_{ub}$-R-GFP URA3* | This work |
| #73 | $P_{MET}$Boi1-CCG | JD47, *$P_{BOI1}$:: $P_{MET17}$ natNT2, BOI1::BOI1-Cherry-$C_{ub}$-R-GFP URA3* | Kustermann *et al.*, 2017 |
| YAD974 | $P_{MET}$Epo1-CCG | JD47, *$P_{EPO1}$:: $P_{MET17}$ natNT2, EPO1::EPO1-Cherry-$C_{ub}$-R-GFP URA3* | Neller *et al.*, 2014 |
| SGY218 | Δbem1<br>pBem1 316 , pRS315 | JD47, *bem1::natNT2,* p*RS315 CEN LEU2* p*$P_{BEM1}$-BEM1 CEN URA3* | This work |
| SGY190 | Δbem1 Δbem2<br>pBem1 316 | JD47, *bem1::natNT2, bem2::CmLEU2,* p*$P_{BEM1}$-BEM1 CEN URA3* | This work |
| SGY211 | Δbem1 Δbem3<br>pBem1 316 | JD47, *bem1::natNT2, bem3::CmLEU2,* p*$P_{BEM1}$-BEM1 CEN URA3* | This work |
| SGY217 | Δbem1 Δbem3<br>pBem1 316<br>pRS314 | JD47, *bem1::natNT2, bem3::CmLEU2,* p*$P_{BEM1}$-BEM1 CEN URA3,* p*RS314 CEN TRP1* | This work |
| SGY216 | Δbem1 Δbem3<br>pBem1 316 ,<br>p$P_{MET}$Gic2$_{PBD}$ | JD47, *bem1::natNT2, bem3::CmLEU2,* p*$P_{BEM1}$-BEM1 CEN URA3,* p*$P_{MET17}$-GIC2$_{1\text{-}208}$-9xMYC CEN TRP1* | This work |
| SGY222 | Δbem1 Δbem3<br>pBem1 316<br>pRS313 | JD47, *bem1::natNT2, , bem3::CmLEU2* p*$P_{BEM1}$-BEM1 316 CEN URA3,* p*RS313 HIS3* | This work |
| SGY223 | Δbem1 Δbem3<br>pBem1 316<br>p$P_{MET17}$-Bem3 | JD47, *bem1::natNT2, bem3::CmLEU2,* p*$P_{BEM1}$-BEM1 316 CEN URA3,* p*$P_{MET17}$GFP-BEM3 CEN HIS3* | This work |
| SGY65 | Δbem1 pBem1 316<br>pRS313 | JD47, *bem1::natNT2,* p*$P_{BEM1}$-BEM1 CEN URA3,* p*RS313 HIS3* | This work |
| SGY66 | Δbem1 pBem1 316<br>p$P_{MET}$Bem1 | JD47, *bem1::natNT2,* p*$P_{BEM1}$-BEM1 CEN URA3,* p*$P_{MET17}$-GFP-BEM1 CEN HIS3* | This work |
| SGY70 | Δbem1 pBem1 316<br>p$P_{MET}$Bem1$_{145\text{-}268}$ | JD47, *bem1::natNT2,* p*$P_{BEM1}$-BEM1 CEN URA3,* p*$P_{MET17}$- GFP-bem1$_{145\text{-}268}$ CEN HIS3* | This work |
| YAD3026 | Bem1$_{WK}$ | JD47, *BEM1::bem1$_{W192K}$* | This work |
| STY1217 | Bem1$_{ND}$ | JD47, *BEM1::bem1$_{N253A}$* | This work |
| STY1221 | Bem1$_{WK/ND}$ | JD47, *BEM1::bem1$_{W192K\ N253A}$* | This work |
| YAD3027 | Bem1$_{WK}$ | ULM53, *BEM1::bem1$_{W192K}$* | This work |
| STY1218 | Bem1$_{ND}$ | ULM53, *BEM1::bem1$_{N253A}$* | This work |
| STY1222 | Bem1$_{WK/ND}$ | ULM53, *BEM1::bem1$_{W192K\ N253A}$* | This work |
| STY1230 | Bem1$_{WK}$<br>p$P_{MET}$Gic2$_{PBD}$ | JD47, *BEM1::bem1$_{W192K}$ ,* p*$P_{MET17}$GIC2$_{1\text{-}208}$-9xMYC CEN TRP1* | This work |
| STY1228 | Bem1$_{ND}$<br>p$P_{MET}$Gic2$_{PBD}$ | JD47, *BEM1::bem1$_{N253A}$ ,* p*$P_{MET17}$GIC2$_{1\text{-}208}$-9xMYC CEN TRP1* | This work |
| STY1232 | Bem1$_{WK/ND}$ | JD47, *BEM1::bem1$_{W192K\ N253A}$ ,* | This work |

| | | | |
|---|---|---|---|
| | pP$_{MET}$Gic2$_{PBD}$ | pP$_{MET17}$GIC2$_{1-208}$-9xMYC CEN TRP1 | |
| LRY418 | Δboi1 Δboi2 Boi1 pP$_{MET}$Gic2$_{PBD}$ | JD47, BOI1::natNT2, BOI2::hphNT1, P$_{BOI1}$::P$_{BOI1}$BOI1 HIS3, pP$_{MET17}$-GIC2$_{1-208}$-9xMYC CEN LEU2 | This work |
| LRY419 | Δboi1 Δboi2 Boi1$_{ΔPxxP}$ pP$_{MET}$Gic2$_{PBD}$ | JD47, BOI1::natNT2, BOI2::hphNT1, P$_{BOI1}$::P$_{BOI1}$ BOI1$_{Δ394-413}$ HIS3, pP$_{MET17}$-GIC2$_{1-208}$-9xMYC CEN LEU2 | This work |
| SGY156 | Δbem1 Bem1 pP$_{MET}$Gic2$_{PBD}$ Shs1-mCherry | JD47, BEM1::natNT2 P$_{BEM1}$::P$_{BEM1}$BEM1 HIS3, pP$_{MET17}$-GIC2$_{1-208}$-9xMYC CEN LEU2, SHS1::SHS1-mCherry URA3 | This work |
| SGY151 | Δbem1 Bem1$_{ND}$ pP$_{MET}$Gic2$_{PBD}$ Shs1-mCherry | JD47, BEM1::natNT2, P$_{BEM1}$::P$_{BEM1}$BEM1 $_{N253A}$ HIS3, pP$_{MET17}$-GIC2$_{1-208}$-9xMYC CEN LEU2, SHS1::SHS1-mCherry URA3 | This work |
| SGY152 | Δbem1 Bem1$_{WK}$ pP$_{MET}$Gic2$_{PBD}$ Shs1-mCherry | JD47, BEM1::natNT2, P$_{BEM1}$::P$_{BEM1}$BEM1 $_{W192K}$ HIS3, pP$_{MET17}$-GIC2$_{1-208}$-9xMYC CEN LEU2, SHS1::SHS1-mCherry URA3 | This work |
| SGY153 | Δste20 Ste20$_{FLPT}$ pP$_{MET}$Gic2$_{PBD}$ Shs1-mCherry | JD47, STE20::natNT2, P$_{STE20}$::P$_{STE20}$STE20 $_{F470L\ P475T}$ HIS3, pP$_{MET17}$-GIC2$_{1-208}$-9xMYC CEN LEU2, SHS1::SHS1-mCherry URA3 | This work |
| SGY154 | Δcla4 Cla4$_{PPAA\ FL}$ pP$_{MET}$Gic2$_{PBD}$ Shs1-mCherry | JD47, CLA4::natNT2, P$_{CLA4}$::P$_{CLA4}$CLA4 $_{F15A\ PP22.25AA\ F451L}$ HIS3, pP$_{MET17}$-GIC2$_{1-208}$-9xMYC CEN LEU2, SHS1::SHS1-mCherry URA3 | This work |
| SGY371 | Δste20 Ste20 pP$_{MET}$Gic2$_{PBD}$ Shs1-mCherry | JD47, STE20::natNT2, P$_{STE}$20::P$_{STE20}$STE20, HIS3, pP$_{MET17}$-GIC2$_{1-208}$-9xMYC CEN LEU2, SHS1::SHS1-mCherry URA3 | This work |
| SGY158 | Δste20 Δcla4 Cla4$_{PPAA\ FL}$ pP$_{MET}$Gic2$_{PBD}$ Shs1-mCherry | JD47, STE20::hphNT1, CLA4::natNT2, P$_{CLA4}$::P$_{CLA4}$CLA4 $_{F15A\ PP22.25AA\ F451L}$ HIS3, pP$_{MET17}$-GIC2$_{1-208}$-9xMYC CEN LEU2, SHS1::SHS1-mCherry URA3 | This work |
| SGY366 | Δste20 Δcla4 Ste20 Cla4 pP$_{MET}$Gic2$_{PBD}$ Shs1-mCherry | JD47, STE20::hphNT1, CLA4::natNT2, P$_{CLA4}$::P$_{CLA4}$CLA4 HIS3, P$_{STE}$20::P$_{STE20}$STE20 TRP1, pP$_{MET17}$-GIC2$_{1-208}$-9xMYC CEN LEU2, SHS1::SHS1-mCherry URA3 | This work |
| SGY369 | Δste20 Δcla4 Ste20 $_{FLPT}$ Cla4 $_{PPAA\ FL}$ pP$_{MET}$Gic2$_{PBD}$ Shs1-mCherry | JD47, STE20::hphNT1, CLA4::natNT2, P$_{CLA4}$::P$_{CLA4}$CLA4 $_{F15A\ PP22.25AA\ F451L}$ HIS3, P$_{STE}$20::P$_{STE20}$STE20 $_{F470L\ P475T}$ TRP1, pP$_{MET17}$-GIC2$_{1-208}$-9xMYC CEN LEU2, SHS1::SHS1-mCherry URA3 | This work |
| SGY368 | Δcla4 Cla4 pP$_{MET}$Gic2$_{PBD}$ Shs1-mCherry | JD47, CLA4::natNT2, P$_{CLA4}$::P$_{CLA4}$CLA4 HIS3, pP$_{MET17}$-GIC2$_{1-208}$-9xMYC CEN LEU2, SHS1::SHS1-mCherry URA3 | This work |
| LRY443 | Δboi1 Δboi2 Boi1 Bem1-GFP 304 Shs1-mCherry | JD47, BOI1::natNT2, BOI2::hphNT1, P$_{BOI1}$::P$_{BOI1}$ BOI1 HIS3, BEM1::BEM1-GFP TRP1, SHS1::SHS1-mCherry URA3 | This work |
| LRY444 | Δboi1 Δboi2 Boi1$_{ΔPxxP}$ Bem1-GFP 304 Shs1-mCherry | JD47, BOI1::natNT2, BOI2::hphNT1, P$_{BOI1}$::P$_{BOI1}$ BOI1$_{Δ394-413}$ HIS3, BEM1::BEM1-GFP TRP1, SHS1::SHS1-mCherry URA3 | This work |
| SGY265 | Δcyk3 Δhof1 pHof1 316 | JD47, CYK3::kanMX, HOF1::natNT2 pP$_{HOF1}$HOF1 CEN URA3, pRS315 | This work |
| SGY271 | Δcyk3 Δhof1 Δste20 pHof1 316 | JD47, CYK3::kanMX, HOF1::natNT2 pP$_{HOF1}$HOF1 CEN URA3, STE20::HISMX pRS314 TRP1, pRS315 LEU2 | This work |
| SGY273 | Δcyk3 Δhof1 Ste20$_{FLPT}$ pHof1 316 | JD47, CYK3::kanMX, HOF1::natNT2 pP$_{HOF1}$HOF1 CEN URA3, STE20::HISMX P$_{STE20}$::P$_{STE20}$STE20 TRP1, pRS315 LEU2 | This work |

| | | | |
|---|---|---|---|
| SGY51 | pP$_{MET}$GFP-Bem1$_{145-268}$ | JD47, p*P*$_{MET17}$*GFP-bem1*$_{145-268}$ *CEN HIS3* | This work |
| SGY27 | pP$_{MET}$GFP-Bem1$_{145-268\ ND}$ | JD47, p*P*$_{MET17}$*GFP-bem1*$_{145-268\ ND}$ *CEN HIS3* | This work |
| SGY63 | pP$_{MET}$GFP-Bem1$_{145-268\ WK}$ | JD47, p*P*$_{MET17}$*GFP-bem1*$_{145-268\ WK}$ *CEN HIS3* | This work |
| YAD860 | Bem1-GFP | JD47, *BEM1::BEM1-GFP TRP1* | This work |
| STY560 | Bem1-GFP Δgps1 | JD47, *BEM1::BEM1-GFP TRP1, GPS1::HISMX* | This work |
| STY528 | Bem1-GFP Nba1$_{\Delta PxxP}$ | JD47, *BEM1::BEM1-GFP TRP1, NBA1::nba1*$_{PPR280,282AAA}$ | This work |
| YAD3474 | Bem1-GFP Fir1$_{\Delta PxxP}$ | JD47, *BEM1::BEM1-GFP TRP1, FIR1::fir1*$_{PKRSPLR\ 525,531\ AKSSALS}$ | This work |
| YAD3476 | Bem1-GFP Nba1$_{\Delta PxxP}$ Fir1$_{\Delta PxxP}$ | JD47, *BEM1::BEM1-GFP TRP1, NBA1::nba1*$_{PPR280,282AAA}$, *FIR1::fir1*$_{PKRSPLR\ 525,531\ AKSSALS}$ | This work |
| UNY318 | Bem1-GFP Boi1$_{WK}$ Boi2$_{WK}$ | JD47, *BEM1::BEM1-GFP TRP1, BOI1::boi1*$_{W53K}$, *BOI2::boi2*$_{W83K}$ | This work |
| ARY60 | Cdc24-GFP | JD47, *CDC24::CDC24-GFP TRP1* | This work |
| STY792 | Cdc24-GFP Δgps1 | JD47, *CDC24::CDC24-GFP TRP1 GPS1::HISMX* | This work |
| STY833 | Cdc24-GFP Nba1$_{\Delta PxxP}$ | JD47, *CDC24::CDC24-GFP TRP1, NBA1::nba1*$_{PPR280,282AAA}$, | This work |
| YAD3473 | Cdc24-GFP Fir1$_{\Delta PxxP}$ | JD47, *CDC24::CDC24-GFP TRP1, FIR1::fir1*$_{PKRSPLR\ 525,531\ AKSSALS}$ | This work |
| YAD3475 | Cdc24-GFP Nba1$_{\Delta PxxP}$ Fir1$_{\Delta PxxP}$ | JD47, *CDC24::CDC24-GFP TRP1, NBA1::nba1*$_{PPR280,282AAA}$, *FIR1::fir1*$_{PKRSPLR\ 525,531\ AKSSALS}$ | This work |
| UNY317 | Cdc24-GFP Boi1$_{WK}$ Boi2$_{WK}$ | JD47, *CDC24::CDC24-GFP TRP1, BOI1::boi1*$_{W53K}$, *BOI2::boi2*$_{W83K}$ | This work |
| STY1134 | Cdc24-GFP Bem1$_{WK}$ | JD47, *CDC24::CDC24-GFP TRP1, BEM1::bem1*$_{W192K}$ | This work |
| STY578 | Boi1-GFP | JD47, *BOI1::BOI1-GFP TRP1* | This work |
| STY563 | Boi1-GFP Δgps1 | JD47, *BOI1::BOI1-GFP TRP1, GPS1::HISMX* | This work |
| STY834 | Boi1-GFP Nba1$_{\Delta PxxP}$ | JD47, *BOI1::BOI1-GFP TRP1, NBA1::nba1*$_{PPR280,282AAA}$, | This work |
| YAD3467 | Boi1-GFP Fir1$_{\Delta PxxP}$ | JD47, *BOI1::BOI1-GFP TRP1, FIR1::fir1*$_{PKRSPLR\ 525,531\ AKSSALS}$ | This work |
| YAD3471 | Boi1-GFP Nba1$_{\Delta PxxP}$ Fir1$_{\Delta PxxP}$ | JD47, *BOI1::BOI1-GFP TRP1, NBA1::nba1*$_{PPR280,282AAA}$, *FIR1::fir1*$_{PKRSPLR\ 525,531\ AKSSALS}$ | This work |
| STY804 | P$_{MET}$SH3$_{Boi1}$-GFP | JD47, *ura3-52::URA3 P*$_{MET17}$*boi1*$_{1-83}$*-GFP* | This work |
| STY836 | P$_{MET}$SH3$_{Boi1}$-GFP Nba1$_{\Delta PxxP}$ | JD47, *ura3-52::URA3 P*$_{MET17}$*boi1*$_{1-83}$*-GFP, NBA1::nba1*$_{PPR280,282AAA}$ | This work |
| YAD3466 | P$_{MET}$SH3$_{Boi1}$-GFP Fir1$_{\Delta PxxP}$ | JD47, *ura3-52::URA3 P*$_{MET17}$*boi1*$_{1-83}$*-GFP, FIR1::fir1*$_{PKRSPLR\ 525,531\ AKSSALS}$ | This work |
| YAD3472 | P$_{MET}$SH3$_{Boi1}$-GFP Nba1$_{\Delta PxxP}$ Fir1$_{\Delta PxxP}$ | JD47, *ura3-52::URA3 P*$_{MET17}$*boi1*$_{1-83}$*-GFP, NBA1::nba1*$_{PPR280,282AAA}$, *FIR1::fir1*$_{PKRSPLR\ 525,531\ AKSSALS}$ | This work |
| YAD3228 | P$_{MET}$SH3$_{Boi1}$-GFP Boi1$_{WK}$ Boi2$_{WK}$ | JD47, *ura3-52::URA3 P*$_{MET17}$*boi1*$_{1-83}$*-GFP, BOI1::boi1*$_{W53K}$, *BOI2::boi2*$_{W83K}$ | This work |
| YWY83 | Boi2-GFP | JD47, *BOI2::BOI2-GFP TRP1* | This work |
| STY1110 | Boi2-GFP Δgps1 | JD47, *BOI2::BOI2-GFP TRP1, GPS1::HISMX* | This work |

| | | | |
|---|---|---|---|
| STY835 | Boi2-GFP<br>Nba1$_{\Delta PxxP}$ | JD47, *BOI2::BOI2-GFP TRP1,*<br>*NBA1::nba1*$_{PPR280,282AAA}$ | This work |
| YAD3032 | P$_{MET}$SH3$_{Boi2}$-GFP | JD47, *ura3-52::URA3 P*$_{MET17}$*boi2*$_{1\text{-}160}$*-GFP* | This work |
| STY1109 | P$_{MET}$SH3$_{Boi2}$-GFP<br>Δgps1 | JD47, *ura3-52::URA3 P*$_{MET17}$*boi2*$_{1\text{-}160}$*-GFP,*<br>*GPS1::HISMX* | This work |
| YAD3033 | P$_{MET}$SH3$_{Boi2}$-GFP<br>Nba1$_{\Delta PxxP}$ | JD47, *ura3-52::URA3 P*$_{MET17}$*boi2*$_{1\text{-}160}$*-GFP,*<br>*NBA1::nba1*$_{PPR280,282AAA,}$ | This work |
| STY576 | Nba1-GFP | JD47, *NBA1::NBA1-GFP URA3,* | This work |
| STY577 | Nba1-GFP<br>Δgps1 | JD47, *NBA1::NBA1-GFP URA3,*<br>*GPS1::HISMX* | This work |
| SGY144 | Ste20-GFP | JD47, *STE20::STE20-GFP TRP1* | This work |
| SGY198 | Ste20-GFP<br>Bem1$_{WK}$ | JD47, *STE20::STE20-GFP TRP1*<br>*BEM1::natNT2 P*$_{BEM1}$*::P*$_{BEM1}$*BEM1*$_{W192K}$ *HIS3* | This work |
| SGY143 | Cla4-GFP | JD47, *CLA4::CLA4-GFP TRP1* | This work |
| SGY197 | Cla4-GFP<br>Bem1$_{WK}$ | JD47, *CLA4::CLA4-GFP TRP1*<br>*BEM1::natNT2 P*$_{BEM1}$*::P*$_{BEM1}$*bem1*$_{W192K}$ *HIS3* | This work |
| SGY199 | *Boi1-GFP*<br>Bem1$_{WK}$ | JD47, *BOI1::BOI1-GFP TRP1*<br>*BEM1::natNT2 P*$_{BEM1}$*::P*$_{BEM1}$*bem1*$_{W192K}$ *HIS3* | This work |
| LRY318 | Boi1-GFP<br>Δboi2 | JD47, *BOI1::natNT2, BOI2::hphNT1,*<br>*P*$_{BOI1}$*::P*$_{BOI1}$ *BOI1-GFP HIS3* | This work |
| LRY319 | Boi1$_{\Delta PxxP}$ –GFP<br>Δboi2 | JD47, *BOI1::natNT2, BOI2::hphNT1,*<br>*P*$_{BOI1}$*::P*$_{BOI1}$ *BOI1*$_{\Delta 394\text{-}413}$ *–GFP HIS3* | This work |
| LRY72 | Cla4$_{1\text{-}471}$-9xMyc | JD47, *CLA4::natNT2,*<br>p*P*$_{MET17}$*-cla4*$_{1\text{-}471}$*-9xMYC CEN TRP1* | This work |
| LRY73 | Cla4$_{1\text{-}471\ PPAA\ \text{-}}$9xMyc | JD47, *CLA4::natNT2,*<br>p*P*$_{MET17}$*-cla4*$_{1\text{-}471\ PP22.25AA}$*-9xMYC CEN TRP1* | This work |
| LRY74 | Cla4$_{1\text{-}471\ F15A\ PP22.25AA\ PPAA}$ -9xMyc | JD47, *CLA4::natNT2,* p*P*$_{MET17}$*-Cla4*$_{1\text{-}471\ F15A\ PP22.25AA\ PP458.461AA}$*-9xMYC CEN TRP1* | This work |
| LRY75 | Cla4$_{1\text{-}471\ F15A\ PP22.25AA\ F451L}$ -9xMyc | JD47, *CLA4::natNT2,* p*P*$_{MET17}$*-cla4*$_{1\text{-}471\ F15A\ PP22.25AA\ F451L}$*-9xMYC CEN TRP1* | This work |
